# Single-cell genomic atlas of great ape cerebral organoids uncovers human-specific features of brain development

**DOI:** 10.1101/685057

**Authors:** Sabina Kanton, Michael James Boyle, Zhisong He, Malgorzata Santel, Anne Weigert, Fatima Sanchis Calleja, Leila Sidow, Jonas Fleck, Patricia Guijarro, Dingding Han, Zhengzong Qian, Michael Heide, Wieland Huttner, Philipp Khaitovich, Svante Pääbo, Barbara Treutlein, J. Gray Camp

**Affiliations:** Max Planck Institute for Evolutionary Anthropology, Leipzig, Germany; Department of Biosystems Science and Engineering, ETH Zürich, Basel, Switzerland; CAS Key Laboratory of Computational Biology, CAS-MPG Partner Institute for Computational Biology, Shanghai, China.; Max Planck Institute of Molecular Cell Biology and Genetics, Dresden, Germany; Center for Neurobiology and Brain Restoration, Skolkovo Institute of Science and Technology, Moscow, Russia.; Institute of Molecular and Clinical Ophthalmology, Basel, Switzerland.

## Abstract

The human brain has changed dramatically since humans diverged from our closest living relatives, chimpanzees and the other great apes^1–5^. However, the genetic and developmental programs underlying this divergence are not fully understood^6–8^. Here, we have analyzed stem cell-derived cerebral organoids using single-cell transcriptomics (scRNA-seq) and accessible chromatin profiling (scATAC-seq) to explore gene regulatory changes that are specific to humans. We first analyze cell composition and reconstruct differentiation trajectories over the entire course of human cerebral organoid development from pluripotency, through neuroectoderm and neuroepithelial stages, followed by divergence into neuronal fates within the dorsal and ventral forebrain, midbrain and hindbrain regions. We find that brain region composition varies in organoids from different iPSC lines, yet regional gene expression patterns are largely reproducible across individuals. We then analyze chimpanzee and macaque cerebral organoids and find that human neuronal development proceeds at a delayed pace relative to the other two primates. Through pseudotemporal alignment of differentiation paths, we identify human-specific gene expression resolved to distinct cell states along progenitor to neuron lineages in the cortex. We find that chromatin accessibility is dynamic during cortex development, and identify instances of accessibility divergence between human and chimpanzee that correlate with human-specific gene expression and genetic change. Finally, we map human-specific expression in adult prefrontal cortex using single-nucleus RNA-seq and find developmental differences that persist into adulthood, as well as cell state-specific changes that occur exclusively in the adult brain. Our data provide a temporal cell atlas of great ape forebrain development, and illuminate dynamic gene regulatory features that are unique to humans.

## MAIN TEXT

Bulk genomic measurements in primary brain tissue from humans, chimpanzees and other apes have identified molecular features that appear specific to the human brain^9–13^. These studies have been limited to a snapshot of adult brain tissues, or average measurements across heterogeneous cell populations. Time course measurements of rhesus macaque brain development provide insights into developmental divergence in primates^14^, but it has been difficult to perform similar experiments in great apes due to the lack of available tissue. Cerebral organoids^15^ grown from human and other great ape induced pluripotent stem cells (iPSCs)^16^ offer the exciting potential to study the evolution of human brain development in controlled culture environments. Previously, we and others have shown that human and chimpanzee cerebral organoids recapitulate many aspects of *in vivo* cortex development^17–22^. In particular, low-throughput single-cell transcriptomics on cortical-like regions within human and chimpanzee cerebral organoids revealed that gene expression patterns of early fetal neocortex development were largely recapitulated in the organoids^17, 19^, and comparative analyses revealed changes between human and chimpanzee^21^. Higher throughput scRNA-seq methodologies enable genomic dissection of individual organoids with the potential to study gene expression landscapes across multiple brain regions^23, 24^ and from multiple individuals. Here, we set out to use high-throughput single-cell RNA-seq, together with accessible chromatin profiling, to understand human cerebral organoid development from pluripotency, and to explore how human cortical gene expression programs have diverged from the other great apes. We further analyze adult prefrontal cortex tissue using single-nucleus RNA-Seq to reveal the potential and limits of cerebral organoids to study human-specific expression patterns observed in the mature brain.

We first used droplet-based scRNA-seq (10X genomics) to profile cell composition across a time course of human organoid development (pluripotency: 0 days (d); embryoid body: 4d; neuroectoderm: 10d; neuroepithelium: 15d; organoid stages: 1, 2, and 4 months (m)) from two human pluripotent stem cell lines (H9, embryonic stem cell (ESC), 23,226 cells; 409b2, iPSC, 20,272 cells; Fig. 1a,b; Extended Data Fig. 1). Marker gene analysis of two-dimensional t-SNE projections of the data from each time point separately, as well as all time points combined, revealed distinct progenitor, neuronal, astrocyte, and mesenchymal populations that emerged across the time course, with intermixing of iPSC and ESC-derived cells (Fig. 1c; Extended Data Fig. 2). We generated pseudocells by combining nearest neighbors in the high dimension gene expression space, which resulted in a more robust transcript estimation (on average ∼6,000 genes detected per pseudocell compared to ∼3,000 genes per single cell). We then constructed a force-directed k-nearest neighbors graph^25^ to visualize the temporal progression of the data (Fig. 1d, Extended Data Fig. 3). We track a progression through pluripotent, neuroectodermal, and neuroepithelial stem cell states during the first 15 days of differentiation. By 1 month, cells diversify into neural progenitors from multiple brain regions including the forebrain (dorsal and ventral telencephalon, diencephalon), midbrain (mesencephalon), and hindbrain (rhombencephalon). A small subpopulation resembling retinal progenitors of the developing eye field is also present, but these cells were only detected in an iPSC 409b2-derived organoid. In addition, a non-neuronal mesenchymal population appears from both cell lines early in the differentiation time course. By 2 months, excitatory and inhibitory neuronal fates have differentiated from progenitors of multiple brain regions, and by 4 months astrocytes have emerged (Fig. 1e). These observations were based on the supervised analysis of known marker genes and inspection of *in situ* patterns from the Allen Developing Mouse Brain Atlas, comparisons to bulk RNA-seq data from microdissected regions of the developing human brain (BrainSpan data^26^) and single-cell reference maps of cell prototypes from the dorsal and ventral telencephalon^27^, as well as the analysis of spliced and unspliced transcripts using RNA velocity^28^ (Fig. 1e,f; Extended Data Fig. 3-4). Together, this data provides a temporally and pseudotemporally resolved gene expression atlas of the earliest stages of human brain development.

**Figure 1:**
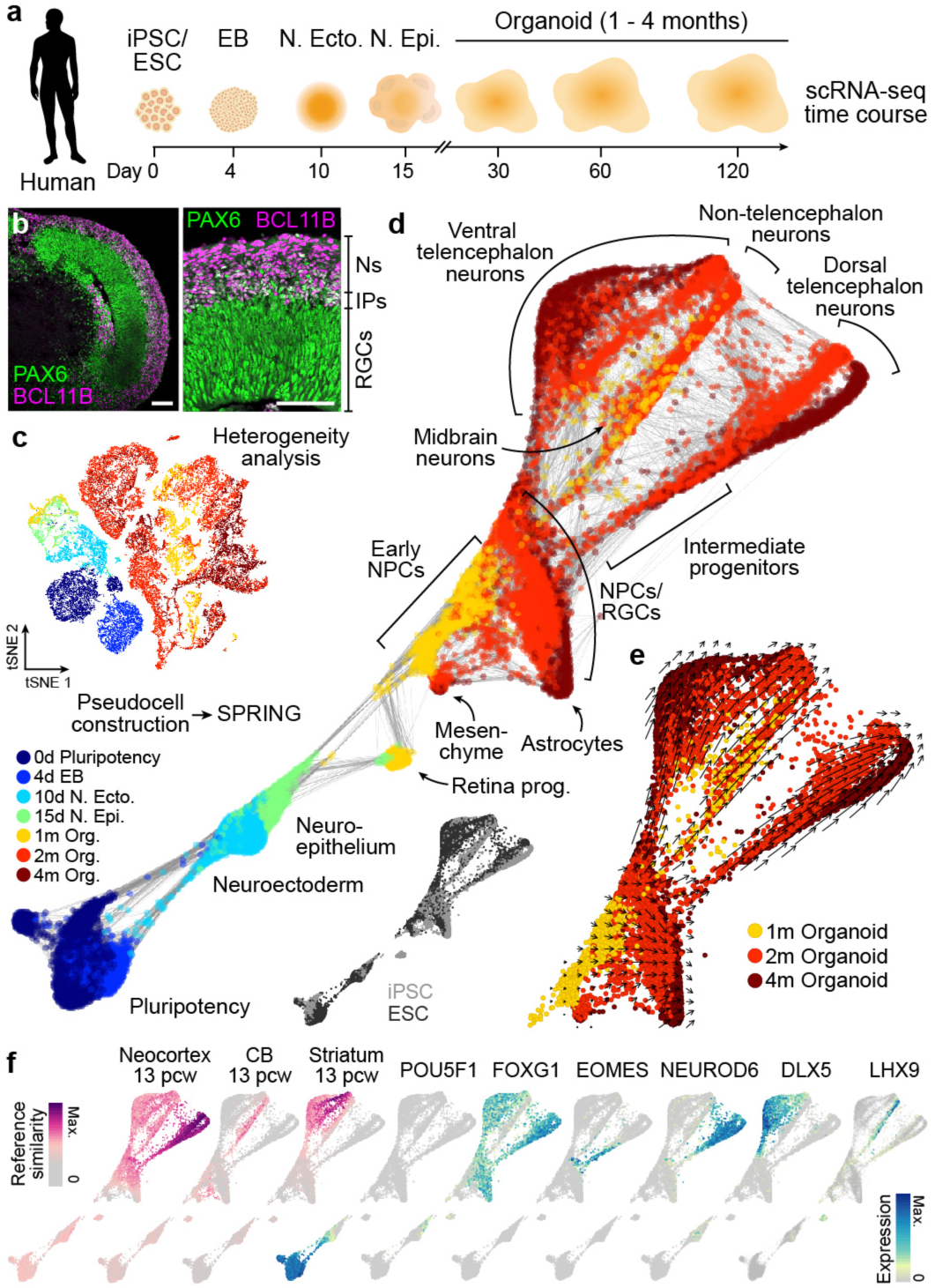
Reconstructing human cerebral organoid differentiation from pluripotency. (a) scRNA-seq was performed on iPSC- and ESC-derived cells at different time points during cerebral organoid differentiation from pluripotency. (b) Immunohistochemical staining for PAX6 (green) and BCL11B/CTIP2 (pink) of a 63 day human organoid from iPSC line 409b2 with a zoom into a cortical-like region (scale bars 100 μm). (c) All time points were combined and cell heterogeneity was assessed using t-distributed stochastic neighbor embedding (tSNE). See Extended Data Fig. 2. (d) Pseudocells were constructed by pooling nearest neighbors and the entire differentiation trajectory was reconstructed using SPRING^25^. Pseudocells are colored by time point or cell line (inset). (e) Tracking unspliced and spliced transcripts using RNA velocity^28^ supports differentiation of progenitor cells into distinct regions of the developing human brain. (f) Left, magenta colored, SPRING plot colored by reference similarity spectrum (RSS) to bulk RNA-seq data generated from diverse brain regions at different time points (Allen Brain Atlas). Shown are the tissues and time points with maximum correlation. Right, cyan-blue colored, SPRING plot colored by marker gene expression.

We next analyzed the reproducibility of these gene expression patterns across PSC lines from different human individuals (Fig. 2a; Extended Data Fig. 5). In addition to the 2 lines (iPSC 409b2 and ESC H9) described above, we generated single-cell transcriptomic data from 2 month old organoids from 5 additional iPSC lines (Sc102a1, 9,525 cells; Wibj2, 13,356; Kucg2, 4,395; Hoik1, 2,660; Sojd3, 3,830), resulting in a total of 62,305 cells from 20 organoids. We identified cells on the neuronal lineage (49,153 cells), for which we constructed pseudocells as described above. We then quantified the similarity (Pearson correlation) of each pseudocell transcriptome to each time point and brain region bulk RNA-seq reference transcriptome from the developing human brain (BrainSpan^29^). We used these similarities to calculate a reference similarity spectrum (RSS) score for each pseudocell, used SPRING to reconstruct the relationships between pseudocells based on RSS, and projected all single cells to the SPRING-based pseudocell embedding. This analysis revealed neuronal differentiation trajectories representing ventral and dorsal telencephalon, as well as distinct populations of cortical excitatory (GLI3, EOMES, NEUROD6), ventral telencephalon inhibitory (DLX1, SOX6, GAD1/2), diencephalon excitatory, diencephalon inhibitory (with Cajal-Retzius cell signatures), mesen- (or midbrain) and rhombencephalon (hindbrain) excitatory, and mesen- and rhombencephalon inhibitory neurons (Fig. 2b-f). Notably, the use of RSS as input for the SPRING analysis instead of the transcriptomes resulted in a well-integrated projection of the data from all human individuals without the need for further integration approaches (Extended Data Fig. 5). Cell annotations were also confirmed through comparisons to voxel maps of *in situ* hybridization patterns from the developing mouse brain (Extended Data Fig. 5). Molecular signatures of the annotated cell types match with those in published scRNA-seq data sets of human cerebral organoids and fetal human brain tissues (Extended Data Fig. 6)^21, 27^. We found that each iPSC line contributed cells to multiple differentiation trajectories, however the proportions of cells in each trajectory varied across organoid and iPSC line (Fig. 2d; Extended Data Fig. 5). For example, over 90% of cells from the line Kucg2 were on the cortical excitatory (dorsal) trajectory in each of the 3 organoids, whereas Hoik1-derived organoids predominantly contained cells from non-telencephalic regions. This tendency of iPSC lines to form different compendiums of brain regions and cell types is consistent with prior work in the literature^23, 30^. Nonetheless, the brain region-specific gene expression patterns across the lines were highly correlated (median of Pearson correlation of pseudotemporally dependent genes: 0.91 and 0.90 for dorsal and ventral trajectories, respectively) and cells from each region clustered together (Fig. 2g-i). This data suggests that even though there is variation in the relative proportion of cell types that form in each organoid, the gene expression patterns within each brain region are largely reproduced across diverse human pluripotent stem cell lines, thus providing a baseline for identifying human-specific gene expression.

**Figure 2:**
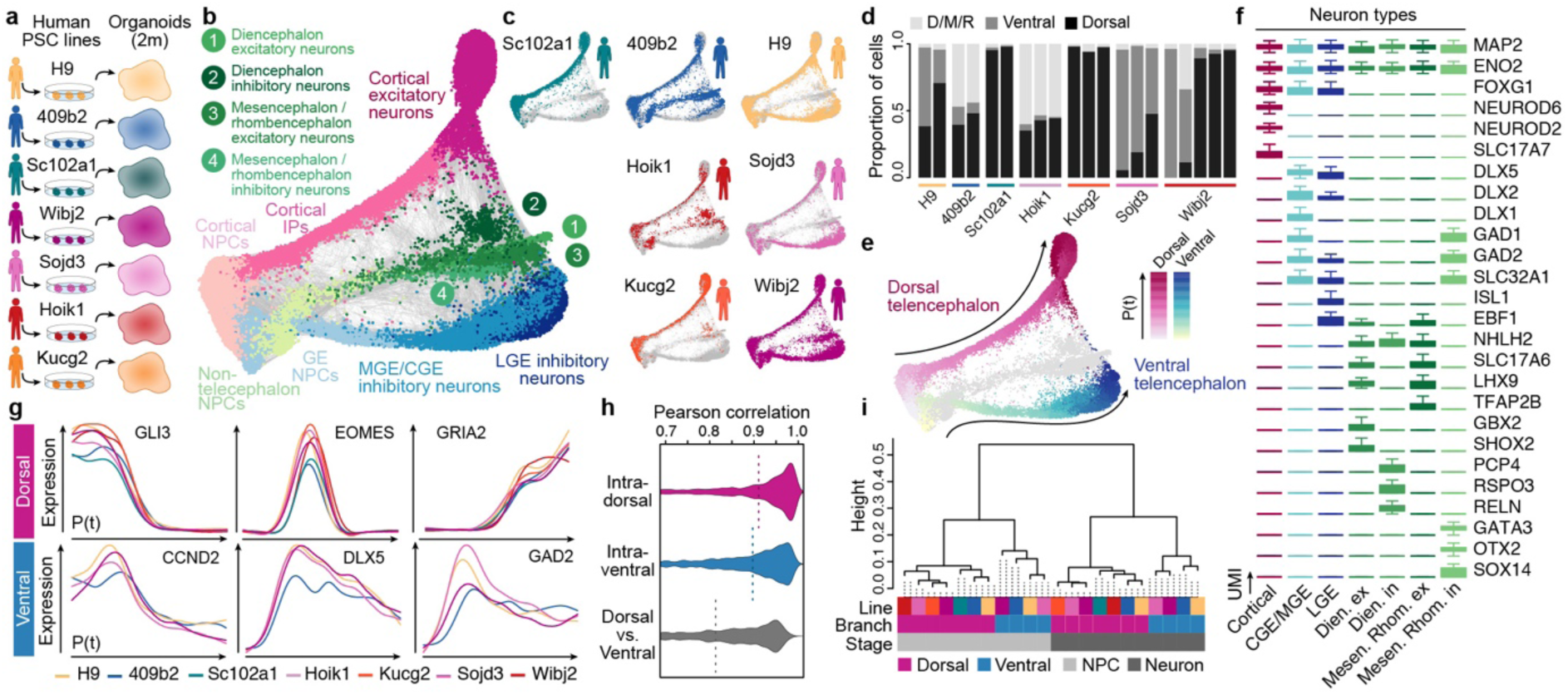
Assessing the reproducibility of gene expression patterns in the human organoid forebrain from iPSC lines from different individuals. (a) To assess the reproducibility of gene expression patterns in organoids, scRNA-seq was performed on 2-month-old human organoids from 6 iPSC lines and 1 ESC (H9) line. (b-c) SPRING reconstruction based on the reference similarity spectrum (RSS) of organoid cells with plots colored by (b) cell types and by (c) line. (d) Proportion of cells per organoid that are within the dorsal telencephalon, ventral telencephalon, or diencephalon, mesencephalon and rhombencephalon neuronal branches. The data shows that there is variation in the types of cells that form in each organoid. (e) Pseudotime along the dorsal telencephalon and ventral telencephalon branch. (f) Boxplots (interquartile range with minimum and maximum, outliers removed) showing expression of marker genes for major neuron populations that emerge in the human cerebral organoids. CGE/MGE, caudal/medial ganglionic eminence; LGE, lateral ganglionic eminence; Dien. ex., diencephalon excitatory; Dien. in., diencephalon inhibitory; Mesen. Rhom. ex., Mesencephalon / rhombencephalon excitatory; Mesen. Rhom. in., Mesencephalon / rhombencephalon inhibitory. (g) Pseudotemporal expression patterns of neuronal differentiation markers for the dorsal (cortex, upper) and ventral telencephalon trajectories (lower) for each line. (h) Correlations of pseudotime-dependent gene expression patterns between cells within dorsal (upper) or ventral (middle) telencephalon branches, and between dorsal and ventral cells from the same line (lower). (i) Dendrogram based on pairwise correlations between cells from different lines/branches/stages based on pseudotime-dependent gene expression patterns. The clustering shows that differences between progenitors and neurons, as well as the variation between those cell types in different brain regions, are larger than variation between cell lines.

We next used chimpanzee organoids to identify features that differ from early human brain development. As for humans, we generated an atlas of gene expression across chimpanzee organoid development from pluripotency to 4 months in culture (Fig. 3a, Extended Data Fig. 7). Similar to human, chimpanzee organoids were morphologically complex with cortical-like regions containing apically located PAX6-positive progenitor cells and basally located neurons (Fig. 3b), with intermediates in between^19^. From the scRNA-seq data, we identified dorsal and ventral telencephalon differentiation trajectories, as well as rhombencephalon cell populations in chimpanzee organoids with a graph topology and gene expression patterns that were very similar to those observed in human (Fig. 3c-e). One difference was that upper and deep layer neurons in these chimpanzee organoids appeared to diversify and mature at an earlier stage along the cortical excitatory trajectory (Extended Data Fig. 8). We used a time warping algorithm to align the iPSC-to-cortical excitatory neuron pseudotimes from human and chimpanzee and observed that the later time points in chimpanzee failed to map to a human pseudocell counterpart (Fig. 3f,g). This observation suggested that neurons within chimpanzee organoid cortical regions may develop at a faster rate than in humans. In support of this observation we found that human dorsal telencephalon pseudocells projected to earlier parts of the developmental trajectory reconstructed in fetal human brain tissues^27^ than the chimp counterpart (Fig. 3h). In addition, we found that neuron maturation scores based on the cumulative expression of neuron projection, synapse assembly, and neurotransmitter secretion genes increased to higher levels in chimpanzee relative to human neurons over the time course (Fig. 3i, Extended Data Fig. 8). We also observed significantly more astrocytes relative to the number of radial glia (RG) cells in chimpanzee in 2- and 4-month organoids compared to humans (Fig. 3j). To determine the heterogeneity in organoid maturation across iPSC lines, we analyzed single-cell RNA-seq data (Smart-seq2) from additional human (15 individuals, 52 organoids) and chimpanzee (11 individuals, 38 organoids) organoids^19, 21^. Indeed, we found that there is heterogeneity in terms of upper and deep layer bifurcation timing that could be dependent on iPSC lines or organoid protocols. However, we found significant consistency across lines, organoids, and protocols in our assessment of neuron maturation based on gene expression (Extended Data Fig. 8). To determine if this difference in maturation timing is specific to humans, we generated cerebral organoids from macaque ESCs and analyzed 2 and 4 month organoids using single-cell transcriptomics (Extended Data Fig. 9). We found that upper and deep layer neurons diverge as early as 2 months, and that neurons mature over an even shorter time frame than in chimpanzees. Also, more upper layer neurons were detected in fetal macaque brain tissues compared with fetal human brain tissues with similar ages (Extended Data Fig. 8). This is consistent with expectations from previous reports comparing human and macaque brain development *in vivo*^14^ and *in vitro* 2D cultures^18, 31^. Together, this data suggests that delayed maturation of the human brain^11, 29, 32, 33^ can be traced back to very early stages of brain development.

**Figure 3:**
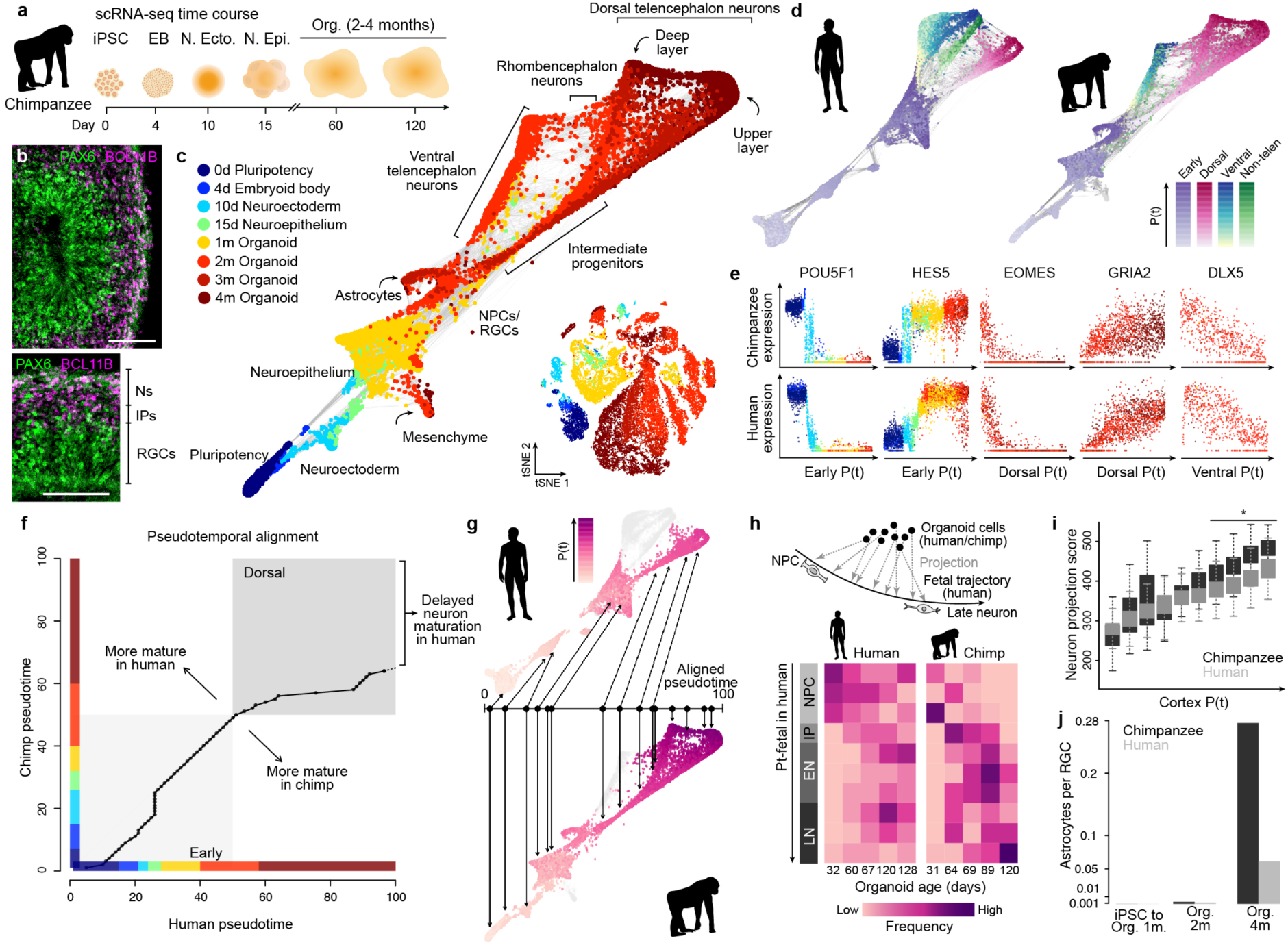
Chimpanzee cerebral organoid reconstructions reveal heterochrony in early cortex development. (a) scRNA-seq was performed on chimpanzee iPSC-derived cells at different time points during cerebral organoid differentiation from pluripotency. (b) Immunohistochemical staining for PAX6 (green) and BCL11B/CTIP2 (pink) of a 63-day chimpanzee organoid from iPSC line SandraA with a zoom into a cortical-like region (scale bars 100 μm). (c) All time points were combined and cell heterogeneity was assessed using tSNE (Extended Data Fig. 7). Pseudocells were constructed by pooling nearest neighbors and the entire differentiation trajectory was reconstructed using SPRING. Cells and pseudocells are colored by time point. (d) SPRING plots of human (left) and chimpanzee (right), colored by stage and lineage pseudotimes. (e) Marker gene expression along pseudotime trajectories in chimpanzee (upper) and human (lower). (f) Alignment of human and chimpanzee pseudotimes after combining pseudocells from the early stages and the dorsal forebrain lineage. The later chimpanzee pseudotime points fail to align with human pseudocells. (g) SPRING plots of human (upper) and chimpanzee (lower) organoid development, colored by the aligned pseudotimes with chimpanzee pseudotime as the template. (h) Projection of human and chimpanzee organoid cells to human fetal brain data reveals higher similarity of chimpanzee organoid cells to later stages of development compared to human organoid cells. (i) Boxplots (interquartile range with minimum and maximum, outliers removed) showing neuron projection scores defined as the sum expression of genes related to neuron projection in human and chimpanzee along the unaligned cortical pseudotimes. (j) Number of astrocytes captured by scRNA-seq in organoids at different time points, normalized by the number of radial glia for each respective time point.

We next wanted to detect gene expression changes in the developing cortex that have occurred since humans diverged with chimpanzees (Fig. 4a). We first inspected the expression patterns of human genes resulting from duplication or rearrangement that do not exist in other apes (Extended Data Fig. 10)^34–37^. We found that 22 of these 41 genes are detected in our human cerebral organoid data, and four of them (ARHGAP11B, FAM72B, FAM72C, FAM72D) are highly-specific to G2M phase progenitors of the dorsal and ventral telencephalon (Extended Data Fig. 10). ARHGAP11B has previously been shown to regulate basal radial glia cell proliferation and self-renewal^38^ and our data highlights the specificity of expression to a distinct phase of the cell cycle, and shows that expression is highly specific to RG progenitors along the time course of cortex development.

To identify quantitative gene expression differences between the primates, we first aligned all human, chimpanzee and macaque reads to a consensus genome and then aligned pseudotimes of dorsal telencephalon progenitor to early-born deep layer neuron trajectory between the species (Fig. 4b; Extended Data Fig. 11). We searched for genes that vary in expression along the pseudotime in each species, and find that 76.6% of these pseudotemporally dynamic genes have a conserved expression pattern. They represent ancestral gene regulatory programs that have been preserved in the primate developing cortex. We then searched for genes that are differentially expressed specifically on the human branch, and identified 98 genes, 96 of which clustered into seven different pseudotemporal patterns (Fig. 4c,d). Notably, clusters 1, 2, and 3 were enriched in human RGs, IPs, and neurons, respectively, and projections onto the entire human and chimpanzee cortical differentiation trajectory from pluripotency revealed specificity of differential expression to these cortical excitatory cell populations (Fig. 4e,f). Surprisingly, we observed that most of the human-specific deviations from chimpanzee and macaque were expression gains, rather than the loss of expression, in humans (Fig. 4g). This gain of expression skew was also observed for chimpanzee-specific changes relative to human and macaque (Extended Data Fig. 11). Our interpretation is that it could be more deleterious to lose a highly conserved gene expression pattern than it is to gain the expression of a new gene. Genes with gain of human-specific expression have no specific gene ontology enrichment, but are predicted to be involved in diverse cell biological processes including RG proliferation, neuron migration, neurite formation and are localized to different components of maturing neurons including axons, dendrites, and synapses (Fig. 4h). When comparing our results to previously published datasets generated using different single-cell RNA-seq methods (Fluidigm C1, Smart-seq2), as well as to human and macaque fetal data, we find strong overlap between the datasets (Extended Data Fig. 11). We also performed a similar comparison between human and chimp using cells of ventral telencephalon identity, and find 92 human-chimp DE genes, with 17% being distinct from what was observed in the cortex (Extended Data Fig. 12). Together, this analysis identifies human-specific gene expression changes that can be specific to certain cell states within the developing human forebrain.

**Figure 4:**
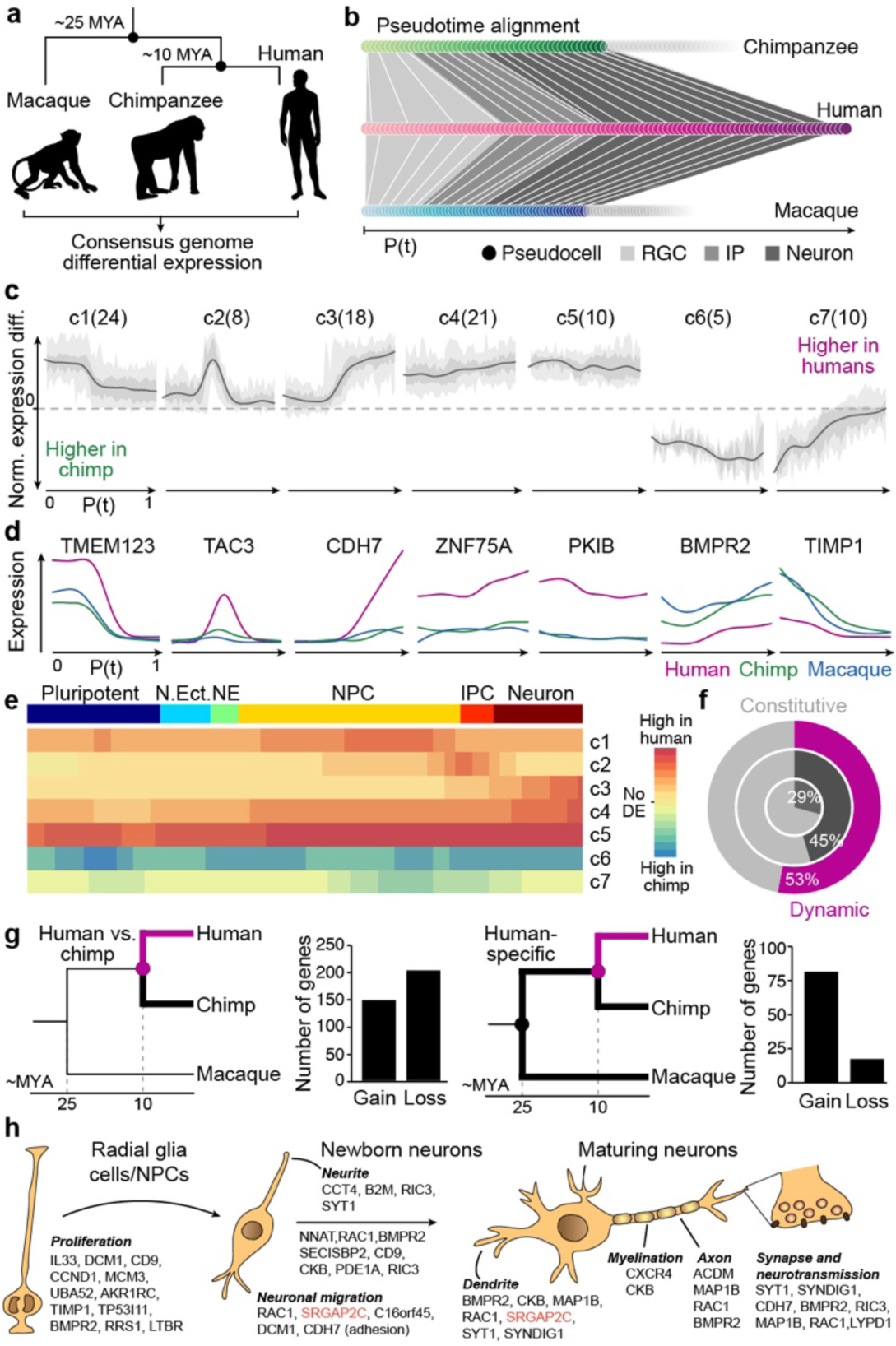
Identification of human-specific changes in corticogenic gene expression. (a) scRNA-seq was performed on 2-4 month-old organoids from human, chimpanzee and macaque lines. Reads were aligned to a consensus genome, to analyze human-specific gene expression. (b) A schematic of time warping alignment of pseudotimes of the cortical progenitor to deeper layer neuron trajectories in human, chimpanzee and macaque in order to perform differential gene expression analysis. (c) Differential expression analysis of the aligned data reveals 7 clusters of genes with distinct human-specific pseudotemporal expression patterns. Average differential expressions of cluster genes are plotted over pseudotime for organoid dorsal forebrain pseudocells with 50% and 95% confidence intervals shown in dark and light grey, respectively. The number of genes per cluster are shown in parenthesis. (d) Pseudotemporal expression pattern of exemplary genes with human-specific expression changes for each of the 7 clusters, comparing human (pink), chimpanzee (green) and macaque (blue) expression. (e) Average human-chimpanzee differential expression patterns along the trajectory from pluripotent cells to cortical neurons shown for the 7 clusters of genes with human-specific pseudotemporal expression changes in organoid cells. (f) Proportions of genes with pseudotime-dependent (dynamic, dark) or constitutive expression patterns (grey), in all (inner), expression-controlled background (middle) and human-specific differentially expressed (outer) genes. (g) Number of differentially expressed genes in a direct human vs. chimpanzee (left) or human vs. chimpanzee and macaque (right) comparison grouped by gain or loss of expression in humans. A gain of expression specifically in humans is more likely than a loss of expression pattern conserved in all primates. (h) Functional annotations of genes with human-specific expression patterns based on GO annotations related to brain development and neurogenesis. Only the human-specific DE genes with consistent human-chimpanzee or human-macaque DE detected in at least one of the three C1-based scRNA-seq data sets are shown (Extended Data Fig. 11). Human-specific duplicated genes are marked in red.

To identify potential mechanisms that could underlie the human-specific gain and loss of gene expression in the cortex, we performed bulk and single-cell accessible chromatin profiling (scATAC-seq, Fluidigm C1) along the differentiation time course from pluripotency to 4 month organoids in human and chimpanzee (Fig. 5a). For the organoid time points, we enriched for dorsal telencephalon by using microdissected cortical regions as input for scATAC-seq (Extended Data Fig. 13). Aggregating the data from each single cell revealed a strong signal-to-noise ratio, and the organoid data was highly similar to bulk fetal brain DNase hypersensitivity^39^ (Fig. 5b), and overlapped forebrain regulatory regions (Extended Data Fig. 13). Accessible regions were scanned for transcription factor binding motifs and k-mers (7 nucleotides in length) to identify features that differ among cells and correlate with accessibility variation^40^. These features were used to visualize cell similarity in a two-dimensional t-SNE projection, which separated iPS, EB, neuroectodermal, neuroepithelial, cortical neural progenitor (NPC) and neuronal cells in both human and chimpanzee (Fig. 5c; Extended data Fig. 13). We then ordered cells in pseudotime using diffusion maps, allowing us to monitor transcription factor binding motifs and chromatin accessibility dynamically over the differentiation path from pluripotency to cortical neurons (Fig. 5d; Extended Data Fig. 13-14). In this way, we found that the majority of genes not expressed in cerebral organoids have inaccessible promoters in organoids (Extended Data Fig. 13).

**Figure 5:**
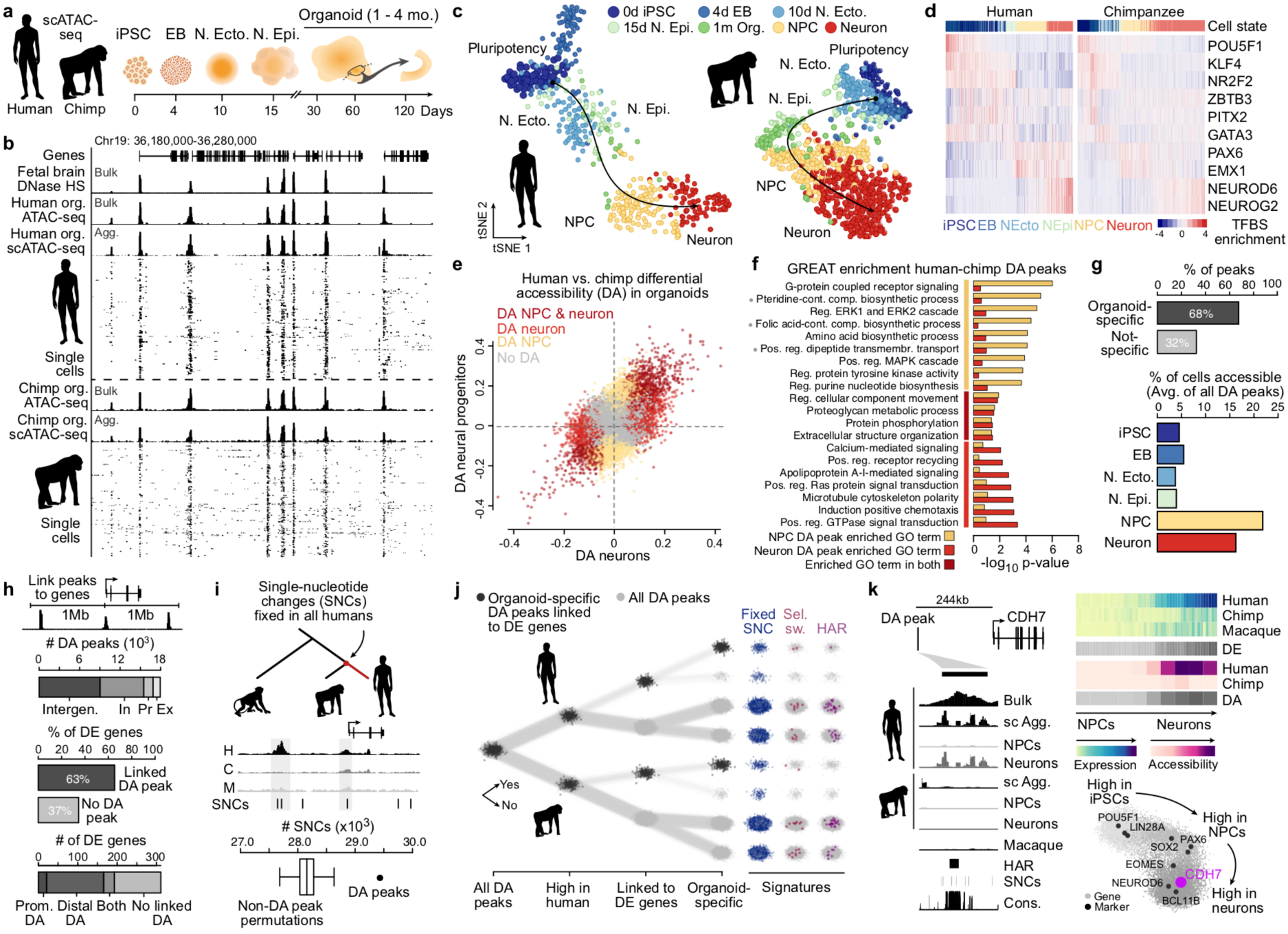
Single-cell ATAC-seq reveals dynamics of chromatin accessibility during cortex development. (a) scATAC-seq was performed at different time points of human and chimpanzee cerebral organoid development from pluripotency to 4 month-old organoids. For the organoid time points, microdissected cortical regions were used as input for scATAC-seq. (b) Bulk, single-cell and aggregated (Agg.) single-cell ATAC-seq profiles from 2-to-4-month-old human and chimpanzee organoids at a representative locus. Data from fetal brain DNase hypersensitivity is shown as a comparison. (c) t-SNE projections of features that differ in accessibility among cells within scATAC-seq peaks per cell^40^ from human (left) and chimpanzee (right) with cells color coded by time point, and 2-4 month-old organoid cells colored by cell state (NPC, neuron). (d) Heatmaps showing binding motif enrichment for selected transcription factors (rows) in all cells (columns) ordered in pseudotime for human (left) and chimpanzee (right). Pseudotime order was constructed using diffusion maps (Extended Data Figure 14). (e) Scaled differential accessibility (DA) between human and chimpanzee NPCs (y axis) and neurons (x axis). Positive values represent higher accessibility in humans. Points represent DA peaks and are color coded by their cell state specificity: NPC (gold), neurons (light red), or DA in both (dark red). (f) Barplot showing the enrichment of selected biological process gene ontology (GO) terms associated with DA peaks between human and chimpanzee in NPCs (gold) or neurons (light red) relative to all accessible organoid peaks. Gray dots next to a GO term name indicate significantly enriched terms after multiple hypothesis test correction (foreground/background hypergeometric test, FDR q<0.05 and 2-fold region-based enrichment). (g) Top, barplot showing the percentage of human-chimp organoid DA peaks that are accessible only at the cerebral organoid stage (“organoid-specific”) or accessible at the cerebral organoid stage and an earlier stage of differentiation (“not-specific”). Bottom, barplot showing the percentage of human cells from each cell state that are accessible at DA peaks. (h) A DA peak was linked to the nearest expressed gene if it fell within 1 Mb of that gene’s transcription start site. Top, stacked barplot shows the number of DA peaks located in exonic (Ex), promoter (Pr), intronic (In), or intergenic (Intergen) regions. Middle, barplot shows the percentage of DE genes linked with DA peaks. Bottom, stacked barplot shows the proportion of DE genes with a DA peak at the promoter region (Prom. DA), distal to the promoter region (Distal DA), at both promoter and distal regions (Both), or no linked DA peak. (i) Comparison of the number of single nucleotide changes (SNCs) derived and fixed in all humans overlapping DA peaks and non-DA peaks (randomly sampled to match the number and average accessibility of DA peaks). (j) DA peaks are annotated as accessible in human or chimp (first bifurcation: up, more accessible in human; down, more accessible in chimp), linked to a differentially expressed gene between human and chimp (second bifurcation: up, yes; down, no), or having specific expression in organoids relative to other time points during organoid development (third bifurcation: up, yes; down: no). On the right, sites are highlighted that show evolutionary signatures including fixed SNCs (blue), selective sweeps (pink) or human accelerated regions (HAR, purple). (k) Cadherin 7 (CDH7) has human-specific expression that is found in human neurons and has a nearby DA site that overlaps fixed SNCs and a HAR. Signal tracks are shown for human bulk organoid ATAC-seq (bulk), as well as aggregated scATAC-seq organoid cells (sc Agg.), and aggregated scATAC-seq for NPCs and neurons, respectively, for human and chimpanzee. Aggregated scATAC-seq data from macaque organoids are also shown. The bottom-right depicts a gene correlation network from the human cerebral organoid time course scRNA-seq data with CDH7 highlighted.

We next searched for differential accessibility (DA) between human and chimpanzee cortical NPCs and neurons. We identified 8,099 peaks (7.4% of all accessible peaks) that gained accessibility in humans relative to chimpanzee, whereas 9,836 peaks (9% of all accessible peaks) lost accessibility (Fig. 5e). Some of these peaks (2,219, 12.4% of DA peaks) are DA in both NPCs and neurons, however most are specific to either NPCs (9,659, 53.8% of DA peaks) or neurons (6,057, 33.8% of DA peaks) and are enriched for various biological processes relative to all accessible organoid peaks (Fig. 5f). Notably, the majority of DA regions are specifically accessible in organoids relative to the earlier developmental stages (Fig. 5g) and many have been shown to drive reporter expression in the mouse developing forebrain (Extended Data Fig. 15)^41^. Consistent with other analyses of gene regulatory evolution^42, 43^, most DA peaks are located in intergenic or intronic non-protein coding regions of the genome (Fig. 5h). The majority of genes that are differentially expressed between human and chimpanzee along the dorsal telencephalon trajectory have one or more human-chimp DA peaks nearby (63% of differentially expressed protein-coding genes, Fig. 5h). We indeed found that genes with differential expression between human and chimpanzee were significantly more likely to have a nearby differentially accessible region than genes that are not differentially expressed between the species (Extended Data Fig. 15, Kolmogorov–Smirnov test, p<0.05). DA peaks are also significantly enriched for single nucleotide changes (SNCs) that are fixed in all humans and distinct from chimpanzee and other primates^44^ (Fig. 5i). Furthermore, these SNCs generate new or disrupt transcription factor binding sites for TFs that are expressed in organoids (Extended Data Fig. 15).

We annotated organoid-specific peaks that are DA between humans and chimpanzees and are nearby differentially expressed genes with various evolutionary signatures (Fig. 5j). This analysis identified potential regulatory regions that have human-derived fixed SNCs^44^, have undergone accelerated evolution in humans^45–47^, or overlap conserved regions that have been deleted in humans^48^. For instance, we identified 62 human accelerated regions that overlap DA peaks (32 in human DA peaks, 30 in chimp DA peaks), with one of these sites being nearby a gene with human-specific expression. In this case, the potential regulatory region is 244 Kb away from cadherin 7 (CDH7), a gene with higher expression specifically in human cortical neurons, and has increased accessibility in human neurons relative to chimpanzee and macaque (Fig. 5k). We also find DA regions nearby two genes, Ly6/PLAUR domain-containing protein 1 (LYPD1) and Ras-related C3 botulinum toxin substrate 1 (RAC1), that have human-specific expression in NPCs and neurons, respectively. LYPD1 is involved in neurotransmitter receptor-binding and anxiety-related behaviors^49^ and RAC1 is a GTPase involved in diverse processes including glucose uptake and cytoskeletal reorganization and genetic variants in this gene can lead to micro- or macrocephaly^50^ (Extended Data Fig. 15). In addition, we identify 22 regions that are accessible in chimpanzee NPCs or neurons that are highly conserved in mammals, but the DNA has been deleted in humans (so-called human conserved deletion, hCONDELs)^48^ and 1 of these are located nearby a DE gene (FADS1, Supplementary Table 10).

Finally, we wanted to know if the human-specific gene expression patterns observed in the developing brain were stage-specific or if they persist into adulthood. We generated single-nucleus RNA-Seq data from postmortem prefrontal cortex tissue of three human, chimpanzee/bonobo and macaque individuals (50,035 in human, 33,847 in chimp/bonobo and 50,403 in macaque). We obtain spatial information by isolating nuclei from sequential sections sliced from basal to apical positions, which allows us to link cell-type specific differences to cortical layering (Fig. 6a)^9^. By integrating the species using canonical correlation analysis and clustering ^51^, we recover expected cell classes such as excitatory and inhibitory neurons, astrocytes, oligodendrocytes, microglia and endothelial cells (Fig 6b-d). For our purposes, we focused on these broad cell classifications but note that subtypes could be more finely resolved and characterized (Extended Data Fig. 16). Different cell classes and cell types show distinct distribution along layers (Extended Data Fig. 16), which is consistent with previous reports^9^. Notably, we find that the portion of the transcriptome that is specific to neurons is more highly conserved based on sequence constraint than that of other cell types (Wilcoxon’s rank sum test, P<0.001, Fig 6e). Indeed, neuronal markers also show higher conservation than genes with higher expression in earlier pluripotent and progenitor states (Extended Data Fig. 16). Inhibitory neuron markers show slightly higher conservation than excitatory neuron markers in both adult brain and organoids (Fig. 6e and Extended Data Fig. 16), and inhibitory neurons in the adult cortex and organoid ventral telencephalon had fewer human-specific DE genes in comparison with excitatory neurons in the adult cortex and organoid dorsal telencephalon trajectory, respectively. We also find that astrocytes have slightly more differential expression than neurons or oligodendrocytes in adults (Fig. 6e,f; Extended Data Fig. 16). Together, these observations suggest different levels of evolutionary constraint on specific cell types in the cortex.

**Figure 6:**
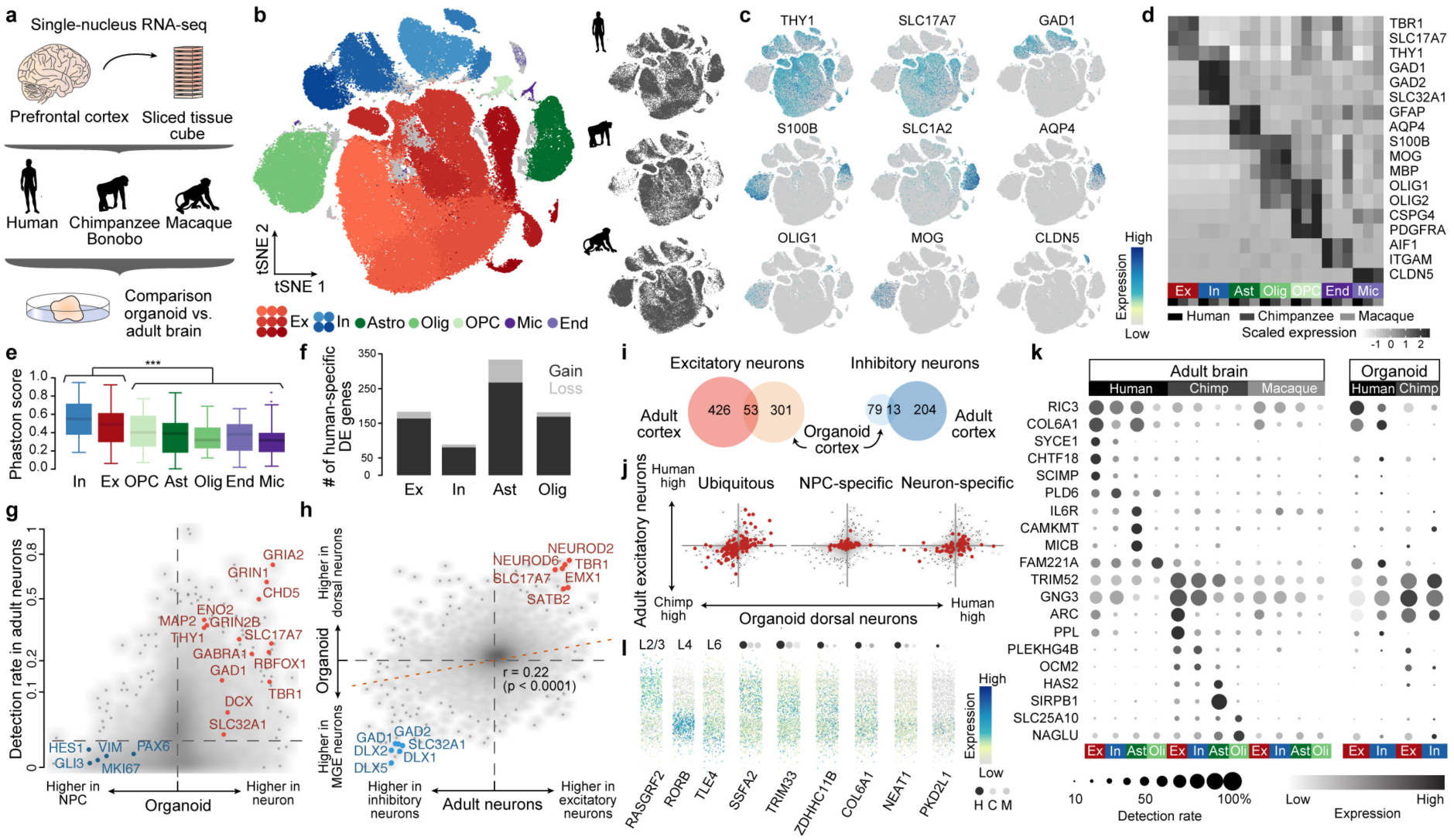
Single-nucleus RNA-Seq of adult prefrontal cortex reveals shared and distinct patterns of gene expression change from development to adulthood. (a) Single-nucleus RNA-Seq was performed on sliced tissue cubes dissected from adult frozen prefrontal cortex tissue from human, chimpanzee/bonobo and macaque. (b) tSNE clustering on CCA integrated data shows different subclasses of major cell classes present in the different species. (c) Feature plots show expression of canonical marker genes for major cell classes based on non-integrated expression values. (d) Average cluster expression of canonical cell type markers separated by species reveal similar patterns of marker gene expression for seven cell classes across species. (e) Genomic conservation based on average phastCon scores of markers for seven cell classes (***: two-sided Wilcoxon’s rank sum test, P<0.0001). (f) Number of genes with human-specific differential expression in each of four major cell classes. Fractions with human-specific gain of expression are shown in dark, and ones with human-specific loss of expression are shown in light. (g) Detection rate in adult tissue of genes being differentially expressed between NPCs and neurons in organoids. (h) Consistency of genes differing between dorsal and ventral forebrain neurons from organoids and excitatory and inhibitory neurons in adult tissue. (i) Overlap of genes with human-chimpanzee differential expression detected in adult neurons and organoid trajectories. (j) Comparison of human-chimpanzee differential expression in adult excitatory neurons and that in organoid dorsal neurons for the robust DE genes detected in the organoid dorsal forebrain trajectory. Three categories of DE genes are highlighted: ubiquitous DE in organoids (left), DE only in NPCs (middle) and DE only in neurons (right). (k) Dotplot showing expression patterns and detection rates across adult and organoid cell classes, for differentially expressed genes in adult cell classes. (l) Predicted laminar expression patterns of six example human-specific DE genes in adult excitatory neurons. Dots on top show their expression in excitatory neurons in the three species, with sizes proportional to their detection rates and darkness showing their average expression levels. Three non-DE canonical layer markers (RASGRF2 for layer 2, RORB for layer 4, TLE4 for layer 6) are also shown.

A substantial fraction of the genes expressed in dorsal and ventral telencephalic organoid neurons are also detected in excitatory and inhibitory neurons in the adult cortex, respectively (Fig. 6g,h). Notably, we find that 53 and 13 genes are commonly detected as DE in the organoid and adult excitatory and inhibitory neurons, respectively, compared to hundreds of genes that are specifically DE in the adult stage (Fig. 6i). Genes with human-chimp DE detected ubiquitously in the organoid dorsal telencephalon show stronger consistency with DE in adult excitatory neurons than genes with DE that is specific to either organoid NPCs or neurons, with NPC-specific DE genes having the weakest consistency in adult (Fig. 6j). In addition, DE genes restricted to organoids or adult show higher expression levels at the stage where DE is detected (Extended Data Fig. 16). There are interesting examples of genes with human-specific DE in adult cell classes (Fig. 6k,l), including genes that are DE in developing and adult neurons, such as COL6A1 which has been shown to have a protective role limiting autophagy and apoptosis in aging neurons^52^ and RIC3 which regulates the number and maturation of acetylcholine-gated ion channels in neurons^53^. We also find genes with human-specific DE in excitatory neurons showing significant layer specificity (Fig. 6l), suggesting their functions in specific subpopulations of cells at specific layer structures. Together, these analyses suggest that, with some exceptions, cortical cell type-specific transcriptome differences between human and chimpanzee are dynamic and linked to developmental stages.

To summarize, we identified patterns of dynamic gene expression and chromatin accessibility differences between human and chimpanzee cerebral organoid development from pluripotency through neuroepithelium, into multiple regions of the ape brain. We provide strong evidence that despite differences in brain region composition, gene expression patterns in the organoid forebrain are largely reproducible across iPSC lines from different individuals. We find that delayed maturation of the human brain begins during the very early stages of brain development. Moreover, we resolve differential gene expression to dynamic cell states upon the ontogenetic path from pluripotency to cortical neurons, and identify regulatory regions that could underlie human-specific innovations in gene expression. Finally, we map human-specific gene expression to cell types in the prefrontal cortex, and identify gene expression patterns that are specific to the adult brain, as well as patterns that can already be detected during development. The data generated in this study are available for exploration via a public interactive browser (https://bioinf.eva.mpg.de/shiny/sample-apps/scApeX/). Taken altogether, these data illuminate features of individual cell states that are uniquely human, and provides an extensive resource to guide exploration into the gene regulatory mechanisms that distinguish the developing human and chimpanzee brains.

## AUTHOR CONTRIBUTIONS

SK, MB grew organoids with assistance from AW, LS, MH. SK performed scRNA-seq and snRNA-Seq with assistance from MS. MB performed scATAC-seq. ZH, MB, and SK analyzed the data. FSC, MH performed immunohistochemical stainings. JF compared organoid scRNA-seq data to mouse voxel maps. PG dissected and sliced tissue for snRNA-Seq. DH and ZQ performed bulk RNA-Seq of adult tissue. SK, MB, ZH, BT, JGC designed the study, and wrote the manuscript with support from PK, WH, SP.

## AUTHOR INFORMATION

Conflict of interest: The authors declare no conflicts of interest.

## ACKNOWLEDGEMENTS

We thank D. Wollny, A. Brazovskaya, K. Köhler, T. Schaffer, B. Schellbach, A. Weihmann, R. Schultz, I. Bünger, M. Dannemann, R. Snabel, B. Vernot, W. Hevers, M. Schörnig, J. Kelso of MPI-EVA, and K. Sekine of Yokohama University for their help with this project. We thank A. Fischer, M. Halbwax, K. Köhler, Anne Weigert and the Tchimpounga Sanctuary for support with generating the JoC iPSC line, Lea Berninger and Jula Peters (MPI-CBG, Dresden) for contributing Sc102a1 and SandraA organoids. Karyotyping was supported by the Stem Cell Engineering Facility, a core facility of CMCB at Technische Universität Dresden. Sorting was in part performed at the CUDZ at the veterinary medicine faculty at the University of Leipzig. This work was supported by the Max Planck Society (BT), Chan Zuckerberg Initiative (BT/JGC), and European Research Council (ANTHROPOID, JGC; ORGANOMICS; BT). SK was supported by the Boehringer Ingelheim Fonds.

## ACCESSION CODE

The single-cell RNA-seq data is being deposited to EMBL-EBI ArrayExpress with the accession number E-MTAB-7552.

## METHODS

### Pluripotent stem cell lines and organoid culture

We acquired 6 human induced pluripotent stem cell (iPSC) lines (Sojd3, Hoik1, Kucg2, Wibj2 from the HipSci resource^54^; h409b2 from the RIKEN BRC cell bank^17^; Sc102a1 from System Biosciences), one human ES cell line (H9, WiCell)^55^, three chimpanzee iPSC lines (SandraA^19^; PR818-5^19^, originally generated by the Gage lab and kindly provided to us by the R. Livesey group; JoC, generated in this study), one bonobo iPSC line (Bokela, generated in this study) and one ES macaque cell line (MN1^18^, kindly provided through the R. Livesey group from Eliza Curnow). The iPSC line JoC (chimpanzee, Tchimpounga Sanctuary) was reprogrammed from blood cells (primary lymphocytes) using plasmid based reprogramming^56^ and Bokela (bonobo, Zoo Leipzig) was reprogrammed from fibroblasts using the StemMACS mRNA transfection kit (Miltenyi Biotec). Cell lines were validated for pluripotency markers by immunhohistochemical stainings using the Human Pluripotent Stem Cell 3-Color Immunohistochemistry Kit (R&D Systems, SC021) and were differentiated into the three different germ layers using the Human Pluripotent Stem Cell Functional Identification kit (R&D Systems) and StemMACS Trilineage Differentiation Kit (Miltenyi Biotec). Karyotyping was carried out using Giemsa banding at the Stem Cell Engineering facility, a core facility of CMCB at Technische Universität Dresden, and karyotypes were found to be normal. Cell lines were cultivated using standard feeder-free conditions in mTeSR1 (StemCell Technologies) and StemMACS iPS-Brew XF (Myltenyi Biotec) on matrigel-coated plates and differentiated into cerebral organoids using a whole organoid differentiation protocol (Lancaster et al. 2014). iPS Brew was used for cultivation of macaque ESCs as well as for EB generation during organoid differentiation for these batches (Supplementary Table 1). Cell lines were regularly tested for mycoplasma using PCR validation (Venor GeM Classic, Minerva Biolabs) and found to be negative.

### Single-cell RNA-seq data generation

A summary of all single-cell experiments can be found in Supplementary Table 1. For organoid experiments (1 month, 2 months, 3 months, 4 months), whole organoids were dissociated for generating single cell gene expression libraries. Briefly, organoids were transferred to HBSS (without Ca^2+^ and Mg^2+^,-/-) and cut into two pieces to clear away debris from the center of the organoid (2-3 washes in total). Organoid pieces were then dissociated using Neural dissociation kit (P) using Papain-based dissociation (Miltenyi Biotec). Organoid pieces were incubated in Papain at 37 °C (enzyme mix 1) for an initial 15 min. followed by addition of Enzyme A (enzyme mix 2) to the Papain mix. Organoid pieces were then triturated using wide bore 1000ml tips and incubated for additional intervals of 5-10 min with triturations between the incubation steps, amounting to a total Papain incubation time of approximately 45 min. Cells were filtered through a 30 μm strainer and washed, centrifuged for 5 min at 300xg and washed 3 times with HBSS (-/-). Cells were then analyzed using Trypan Blue assay, counted using the automated cell counter Countess (Thermo Fisher), and diluted for an appropriate concentration to obtain approximately 6000 cells per lane of a 10X microfluidic chip device. Typically, cells from one organoid were loaded per lane in the microfluidic device, and in some cases organoids from different lines were pooled onto the same lane and demultiplexed based on single-nucleotide polymorphisms. For 1 month organoids, three pooled 409b2 and one H9 organoid were dissociated and cells from the two cell lines were mixed at equal ratios to be loaded on the chip. For as set of 2 month HipSci organoid data, organoids were dissociated for all four HipSci cell lines and pooled at equal ratios to be loaded on one lane of the microfluidic device aiming for 10k cells. Fluidigm C1 data (Supplementary table 1) were generated as previously described ^19^ and cells from chimpanzee SandraA 75d organoids were microdissected regions from vibratome slices for which single cell suspensions were generated as described above. Single cells were then sorted into 96-well plates using a FACS Aria III sorter and further processed using the SmartSeq2 protocol^57^ to generate cDNA and the NexteraXT kit (Illumina) to generate sequencing libraries. All libraries (10X and Fluidigm C1/SmartSeq2) were sequenced on Illumina’s Hiseq2500 platform in paired-end mode (100 bp Fluidigm C1/SmartSeq2; 26+8bp, 100bp 10x).

### Early stages of organoid differentiation (iPS cells to neuroepithelium)

For iPSC/ESC single-cell experiments, cells were detached from cell culture dishes using TrypLExpress (Thermo Fisher) incubation for 5 min. followed by addition of mTeSR1. Cells were centrifuged for 5 min. at 200xg and resuspended in mTeSR1, filtered through a 20 μm strainer and washed with mTeSR1. Cells were then centrifuged again for 5 min. at 200xg and resuspended in mTeSR1, counted, diluted to the same concentration and mixed at equal ratios for the three cell lines to be loaded on the 10X microfluidic chip aiming for 10,000 cells. Thirty embryoid bodies (EBs), 15 neuroectoderms, and 1-3 neuroepithelium of each cell line were pooled for each dissociation, respectively. Cells were obtained by papain dissociation as described above for organoid dissociation, with slightly shorter incubation times in enzyme mix 1 (approximately 30 min.). For 10X experiments, cells from the three different cell lines were diluted and mixed at equal ratios to be loaded on the microfluidic chip device.

Single-cell experiments were conducted using the 10X Chromium Single Cell 3’ v2 Kit following the manufacturer’s instructions. Briefly, cells were mixed with reverse transcription mix, gel beads and oil were loaded on the chip device to be coencapsulated into droplets, which underwent first strand cDNA synthesis thereby tagging mRNAs with a unique molecular identifier (UMI) and a unique cell barcode. All following steps were conducted in bulk by breaking the droplets and cleaning up and amplifying the cDNA. Single-cell libraries were then constructed by fragmentation, end repair and adapter ligation and amplification using library specific index sequences as provided by 10X Genomics. Quantification and quality control of libraries was performed using High Sensitivity DNA assays for Agilent’s Bioanalyzer and sequenced on a HiSeq2500 in Rapid or HighOutput sequencing mode. Typically, one 10X library was sequenced on one lane of a sequencing flow cell, with the exception of the HipSci organoids for which three pooled libraries (each library contained pooled cells from four dissociated HipSci organoids from different cell lines) were sequenced on two lanes of a flow cell. See Table S1 for more details.

### Immunohistochemistry

Organoids were washed in PBS prior to fixing in 4% PFA for 2-4 hours (h). The excess of fixative was removed with three PBS washes and organoids were then transferred to a 30% sucrose solution for 24-48 h for cryoprotection. Finally, organoids were transferred to plastic cryomolds (Tissue Tek) and embedded in OCT compound 4583 (Tissue Tek) for snap-freezing on dry ice. For immunohistochemical stainings, organoids were sectioned in slices of 20 µm thickness using a Leica CM3050 S cryostat and Microm HM 560 (Thermo Fisher Scientific) at -15 to -20°C. Organoid sections were quickly washed in PBS to remove any residual OCT. Then, sections were incubated in antigen retrieval solution (HistoVT One, Nacalai Tesque) at 70°C for 20 min. Excess solution was washed away with PBS and the tissue was incubated in blocking-permeabilizing solution (0.3% Triton, 0.2% Tween-20 and 5% Normal Goat Serum in PBS) for 1h at room temperature. Afterwards, sections were incubated overnight at 4°C in blocking-permeabilizing solution containing antibodies anti-PAX6 (mouse, 1:1000, Thermo Fisher Scientific, MA1-109; rabbit, 1:300, Covance, PRB-278P) and anti-CTIP2 (rat, 1:1000, Abcam, AB18465), anti-SATB2 (rabbit, 1:500, Abcam, Ab92446; mouse, 1:500, Abcam, Ab51502), anti-Tbr2 (mouse, 1:500, MPI-CBG Antibody Facility^35^). On the next day, sections were rinsed three times in PBS before incubation for 1h at room temperature in secondary antibody solution, which contained blocking-permeabilizing solution, DAPI (1:3000), Alexa Fluor 488-conjugated anti-rabbit antibody (goat, 1:1000, Thermo Fisher, A11008), Alexa Fluor 546-conjugated anti-mouse antibody (goat, 1:500, Thermo Fisher Scientific, A-21123), Alexa Fluor 647-conjugated anti-rat antibody (goat, 1:500, Thermo Fisher Scientific, A-21247) and Alexa Fluor 488-conjugated - anti-mouse (A21202) and anti-rat antibody (A21208), Alexa Fluor 555-conjugated anti-rabbit antibody (A31572), Alexa Flour 647-conjugated anti-mouse antibody (A31571) (all donkey-derived, 1:500, Molecular Probes). Finally, remainders of secondary antibody solution were washed off with PBS before covering with ProLong Gold Antifade Mountant medium (Thermo Fisher Scientific). Stained organoid cryosections were imaged using a confocal laser scanning Olympus Fluoview FV1200 microscope and Zeiss LSM 880 Airy upright microscope. Whole-section tilescans composed of 3 different z-plane images (z-step = 5-8 µm) were acquired using a 10X magnification objective, Plan-Apochromat 10x/0.45 M27 and Plan-Apochromat 20x/0.8 M27 objectives. Images were then stitched, stacked and further processed using the Olympus Fluoview 4.2b software and ImageJ (Fiji).

### Single cell RNA-seq data preprocessing

We used Cell Ranger, the set of analysis pipelines suggested by 10X Genomics, to demultiplex raw base call files to FASTQ files and align reads to the human genome and transcriptome (hg38, provided by 10X Genomics) with the default alignment parameters. Pooled samples, including samples from different species or human lines, were then demultiplexed using a two-step procedure based on the read mapping results. In the first step, the genome alignment between human (hg38) and chimpanzee (panTro5) was downloaded from UCSC Genome Browser. Sites with diverged bases between human and chimpanzee were obtained based on the genome alignment. Reads covering the species-diverged sites were collected for each reported cell, with the number of bases matching each species counted. Cells with more than 80% reads covering the species-diverged sites matching with one species were assigned as cells from the species. For those samples with human cells from different lines pooled, a second step of demultiplexing was done using demuxlet^58^, based on the genotyping information of lines downloaded from HipSci websites (Kucg2, Wibj2, Hoik1, Sojd3) or called using bcftools based on the unpooled scRNA-seq data (H9, 409b2). Cells with the best singlet likelihood no less than 50 higher than the second best singlet likelihood and estimated mixture ratio less than 30% were labeled as their best-matched lines. All cells failing to pass any of the above threshold were classified as doublets and excluded from the following analysis.

Seurat^59^ was then applied for further data processing. Cells with more than 6,000 or less than 200 detected genes, as well as those with mitochondrial transcripts proportion higher than 5% were excluded. After the log-normalization, confounding factors including the number of detected genes and proportions of mitochondrial transcripts were also regressed out. Highly variable genes were then obtained as genes with dispersion higher than 0.5 and normalized expression level between 0.0125 and 3, followed by principal component analysis (PCA) based on the z-transformed expression levels of the identified highly variable genes (Supplementary Table 2). The top-20 PCs were used to do clustering using Seurat. Additional quality controls of the measured cells were based on primary cell type predictions by using public human fetal brain scRNA-seq data (Nowakowski data set)^27^. In brief, a Lasso logistic regression model was built, using gene expression ranks of the Nowakowski data set as the training set, to predict the primary cell type identity of each single cell in two-month-old and four-month old organoids. Cells which were predicted to be of ‘glycolysis’ identity were excluded, so as cells in the Seurat clusters where more than 80% of cells were predicted as of ‘glycolysis’ identity. Heterogeneity analysis of human (Extended Data Figure 1, Extended Data Figure 2, Supplementary Tables 3 and 4) and chimpanzee (Extended Data Figure 6, Supplementary Tables 5 and 6) full lineage data was performed using t-stochastic neighbor embedding based on the top principal components identified (top 20 PCs for human, top 15 PCs for chimpanzee). Cluster identities were assigned based on cluster gene markers (Supplementary Table 6) as determined by FindAllMarkers function in Seurat (min percentage of cells expressed = 0.25 and log fold change threshold = 0.25) and gene expression of known marker genes. For human data, cells from 409b2 and H9 and were integrated using canonical correlation analysis (CCA) as implemented in Seurat (v3). Briefly, data were normalized and the top 2000 highly variable genes for 409b2 and H9 cells were determined using the vst method. The datasets were integrated based on the top 20 CCs using the Seurat method by identifying anchors and integrating the datasets. The resulting integrated data were scaled and principal component analysis was performed. Clustering was performed based on the top 20 PCs and using a resolution of 0.6. Feature plots show non-integrated expression values. Cluster markers were determined using Wilcoxon test considering only genes that show a minimum log fold expression change of 0.25 in at least a fraction of 0.25 of cells in the clusters using the non-integrated expression values.

### Reference similarity spectrum (RSS) and construction of pseudocell transcriptomes

The reference similarity spectrum (RSS) of one cell to the Human Developing Brain (HDB) atlas was defined as the normalized similarity between gene expression levels of the cell and gene expression levels of each of the 237 fetal samples with RNA-seq data in the BrainSpan database in Allen Brain Atlas (Extended Data Fig. 3). To increase discrimination of different reference samples, only the highly variable genes of the HDB data set (see Supplementary Table 2), defined based on expression variation-mean comparison of the reference data set, were used for the RSS calculation. Between each cell and each sample in the HDB data set, Pearson correlation coefficient (PCC) was calculated across the HBD-highly variable genes. Z-transformation was then applied to PCCs between each cell to the 237 fetal HDB samples to get the normalized similarities.

To construct pseudocells, single cells were firstly grouped based on their sample sources and Seurat clusters. Within each group of cells, i.e. those cells from the same sample and in the same Seurat cluster, cells were selected randomly with a selection probability of 20%. The selected cells were called pseudocell seeds or territory capitals (Extended Data Fig. 3). The ten nearest neighbors (NN) of each seed, based on Euclidean distances of the top-20 PCs, were then assigned to the seed, forming a pseudocell territory. If one cell was assigned to multiple pseudocell territories, one territory was chosen randomly. The expression level of one gene in each pseudocell was then calculated as the average gene expression level across cells in the pseudocell territory.

### Visualization, lineage identification and pseudotime estimation of pseudocells for reconstructing human cerebral organoid differentiation from pluripotency

First, PCA was applied to a pseudocell expression matrix using the z-transformed expression levels of the highly variable genes as input. Euclidean distance between the top 10 PCs of each pair of pseudocells was calculated and a KNN-network (K=100) was then calculated with the constraint to only consider pseudocells from the same or nearby stages when screening for nearest neighbors. The kNN-network was visualized using SPRING^25^. To construct the pseudotime course of human cerebral organoid differentation from pluripotency, the Walktrap community identification algorithm (implemented in the R package igraph) was applied to the above kNN-network to identify network communities. The resulting communities were manually aggregated into four groups to minimize branches in each group. A diffusion map algorithm (implemented in R package destiny^60^) was applied to pseudocells in each of the four groups, with the expression levels of the highly variable genes of pseudocells as the input. The ranks in DC1 were used as the pseudotimes. We used an F-test based ANOVA analysis to identify genes with pseudotime-dependent expression patterns. In brief, we established a natural splined linear regression model (ns function in the R package splines) with six degrees of freedom (df), with expression levels as the response variable and pseudotimes as the independent variable, for each of the highly variable genes. An F-test was applied, to compare variation explained by the splined linear model with that of the residuals normalized by degrees of freedom. Bonferroni correction was performed across tested genes, with a corrected p-value threshold 0.01 to identify genes with pseudotime-dependent expression. The analysis was applied to the four groups of pseudocells separately.

### Visualization, lineage identification, pseudotime estimation of cells in human two-month-old cerebral organoids from different individuals

Pseudocells were constructed for the human two-month-old organoids as above with constraint on samples and based on cells with predicted primary cell types as one of radial glia, intermediate progenitor (IPC), excitatory neuron, and inhibitory neuron. RSS to the BrainSpan fetal samples was calculated for each pseudocell, with distance between two pseudocells defined as the correlation distance between RSS of the two pseudocells. The kNN-network (k=20) was then constructed and SPRING was used to determine coordinates of pseudocells for visualization. To further discriminate pseudocells representing different neuronal lineages, a Walktrap algorithm for network community identification was applied to the RSS-based kNN-network (k=100). Communities that were significantly connected and showing concordant marker expression or similarity spectrum were aggregated, which resulted in three progenitor-to-neuron trajectories. Based on gene expression level ranks of cells in the three defined trajectories, two Lasso logistic regression models were trained, one for classification of cortical and ventral lineage, while the other one for classification of all the three trajectories. The first model was applied to pseudocells in the community C6 which was significantly connected to cortical and ventral trajectories, while the second model was applied to pseudocells in the community C4, which was significantly connected with both the non-telencephalon pseudocells and community C4. With a unique lineage label defined for each pseudocell, a 1*500 self-organizing-map (SOM) model was trained for each of the three trajectories, using RSS of pseudocells within the lineage as the training data. The index of neuron that one pseudocell was assigned to was used as its pseudotime. Diffusion map analysis was also applied to pseudocells at each trajectory, with highly variable gene expression as the input, with ranks of DC1 defined as alternative pseudotime of pseudocells. Pseudotimes obtained by the two methods are highly correlated (Spearman correlation is 0.91 and 0.92 for the dorsal and ventral telencephalon trajectories, respectively).

To project the single-cell data to the cell embedding space that was defined for pseudocells, two support vector regression (SVR) models (implemented in the R package e1071), each of which was for one dimension of the embedding, were trained using RSS of pseudocells as the training set. The trained models were applied to RSS of single cells for their predicted coordinates. Such coordinates were further refined by pushing each cell to its nearest pseudocell with smallest correlation distance of RSS to be 70% closer.

Similarly, a support vector machine (SVM) model was trained (implemented in the R package e1071) using RSS of pseudocells for the three trajectories, and applied to RSS of single cells for their lineage identity. After that, the corresponding SOM model for pseudotime estimation was applied to RSS of each single cell for its estimated pseudotime.

### Dynamic time warping (dtw)-based alignment of pseudotime courses

We used a dynamic time warping algorithm to align different pseudotime courses. In brief, each pseudotime course was evenly broke into 50 blocks. Average gene expression levels of pseudocells or cells within each block was calculated. Pairwise distances between blocks from the two courses were calculated as the Pearson correlation distance, i.e. 1-PCC, across the highly variable genes in cells of both pseudotime courses. Suppose di,j represents the distance between the i-th block in the reference pseudotime course and the j-th block in the query pseudotime course. We defined D as the alignment distance matrix, where

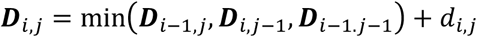

A trace-back procedure was then performed to get the alignment. Three modes of alignment were implemented. In the first mode, the ‘fixed-end’ alignment, the initialization of D was done as:

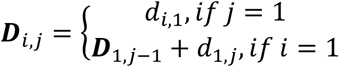

In the other two modes, the ‘fixed-start’ and ‘end-to-end’ alignments, D was initialized as:

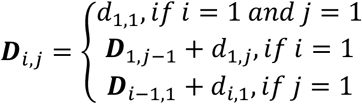

In the trace-back step, a ‘fixed-end’ and ‘end-to-end’ alignment was started from DM,N, where M and N are the numbers of blocks at the reference and query pseudotime courses, respectively. On the other hand, the trace-back step was started from D_m,N,_ where *m* = *argmin*_*i*_(D_*i,n*_). In our study, the ‘fixed-end’ alignment was used to align the cortical and ventral lineage pseudotime course of human organoid cells; the ‘fixed-start’ alignment was used to align pseudotime courses of human and chimpanzee cortical pseudocells; the ‘end-to-end’ was used in the truncated alignment of pseudotime courses of different species.

### Reconstruction of chimpanzee cerebral organoid differentiation from pluripotency

We applied a similar procedure as mentioned above describing the reconstruction of human cerebral organoid differentiation from pluripotency to reconstruct the organoid differentiation trajectory from chimpanzee single-cell RNA-seq data. In brief, the single-cell RNA-seq reads were mapped to the human-chimpanzee-macaque consensus genome and counted using Cell Ranger. Seurat was used for further preprocessing including gene expression normalization, confounding factor regression, PCA and clustering, Cells from organoid samples with predicted primary cell type identity of ‘glycolysis’, as well as cells within clusters with more than 80% cells having ‘glycolysis’ identity, were excluded. Pseudocells were then constructed with a seed selection probability of 20% and constraints on samples and Seurat clusters. PCA was applied to expression levels of highly variable genes across pseudocells, and pairwise distances of pseudocells were calculated as the Euclidean distances between the top-10 PCs. The kNN network (k=100) of pseudocells was constructed, linking every pseudocell with its 100-nearest pseudocells representing the same or nearby stages. Three-month-old and four-month-old organoids were seen as the same stage. The Walktrap network community identification algorithm was applied and the resulted community labels (walktrap communities) of pseudocells were compared with the predicted community labels (projected communities) based on a Lasso logistic regression model trained by ranks of gene expression levels of the human pseudocells representing the human organoid differentiation from pluripotency as described above. Any walktrap community with < 1000 kNN connections with other communities was discarded. One of the four labels: early, cortical, ventral, non-telencephalon was assigned to one walktrap community if more than 95% of pseudocells within the community were from the same group according to their projected communities. For one community with more than 10% of pseudocells with projected communities belonging to both ventral and midbrain-hindbrain groups, the non-telencephalon label was only assigned to pseudocells with projected communities in the non-telencephalon group. The diffusion map algorithm was applied to each of the four pseudocell groups, using the expression levels of highly variable genes as input, to estimate their pseudotimes. For the cortical, ventral and midbrain-hindbrain groups, the ranks of DC1 was used as the pseudotimes. For the early group, a principal curve (implemented in the R package princurve) was fitted in the DC1-DC2 space. The order of pseudocells projecting to the resulted principal curve was used as the pseudotimes.

### Human-chimpanzee-macaque consensus genome

The construction of the consensus genome was performed using the procedure as described^9, 61^. In brief, the chained and netted pairwise genome alignments of the human (hg38) and chimpanzee (panTro5) genomes, and the human and macaque (rheMac8) genomes, were downloaded from UCSC Genome Browser. Based on the downloaded pairwise genome alignments, a multiple genome alignment of human-chimpanzee-macaque was constructed using multiz. Based on the human-chimpanzee-macaque genome alignment, we constructed the consensus genome by masking all discordant sites including mismatches, insertion/deletion (indels), as well as 6-bp flanking regions of indels on the human genome. The obtained consensus genome was indexed with gene annotation in GENCODE v27 for read mapping to the consensus genome with Cell Ranger.

### Pseudotime estimation of cerebral organoid cells in different species

Single cell RNA-seq data of organoids with ages from two-month-old to four-month-old in human, chimpanzee, and macaque were mapped to the human-chimpanzee-macaque consensus genome and counted using Cell Ranger. Further preprocessing using Seurat was applied separately for data from the three species. Only cells with predicted primary cell type identities as radial glia, intermediate progenitors, excitatory neurons, or inhibitory neurons were included in the later analysis. Pseudocells were constructed for humans and chimpanzees, both with a coarse grain ratio of 20% and constraints on samples and Seurat clusters. The RSS to the HDB data set was calculated for each pseudocell, and the SVM model for lineage estimation was applied to estimate the lineage identity of each pseudocell. Focusing on the cortical lineage, a diffusion map analysis was applied to cortical pseudocells of the three species, respectively. The ranks of DC1 were used as the pseudotimes of the pseudocells. In macaque, similar procedure was applied directly to single cells without pseudocell construction.

### Truncated dtw-based alignment of pseudotime courses reprgesenting neural progenitors and deeper layer neurons in different species

We used the first DC discriminating BCL11B+ and SATB2+ cortical neurons (DC3 in chimpanzee, DC4 in macaque) to identify upper layer (UL) neurons, as the pseudocells in the branch with highest expression level of SATB2. To identify potential upper layer neurons in human, we first retrieved markers of upper and deeper layer (DL) excitatory neurons^27^. The sum expression levels of UL and DL markers was then calculated for each pseudocell in human and chimp, with the UL-specificity score (*sUL*) being defined as the UL/DL markers expression ratio. The distribution of *sUL* in UL neurons in chimpanzee was used to determine the threshold to discriminate UL neurons from other cell types (*sUL* > 0). All UL neurons in the three species were excluded from the following analysis.

To correct for the DL neuron maturation timing differences between human and the other two species, a two-step pseudotime course alignment strategy was used. The first step, namely the trimming step, aims to determine the pseudotime points in chimpanzee and macaque which correspond to the latest pseudotime point in human. In brief, an SVR model with Gaussian-kernel was firstly constructed, with chimpanzee or macaque pseudotimes as the response variables and the RSS as the dependent variables. Two models were trained with the chimpanzee pseudocells and macaque cells respectively, and applied to the human pseudocells to predict their corresponding chimpanzee and macaque pseudotime points. Two constrained B-splines regression models (*F_HC_*, *F_HM_*) were then fitted (implemented in the R package *cobs*): human pseudotimes of human pseudocells (*t_h_*) versus their predicted chimpanzee 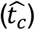 or macaque 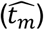 pseudotimes, with constraints of *F*_*HC*_(*t*_*h*_ = 0) = *F*_*HM*_(*t*_*h*_ = 0) = 0. *F*_*HC*_(*t*_*h*_=1) and *F*_*HM*_(*t*_*h*_=1) were used as the pseudotime thresholds to select chimpanzee pseudocells and macaque cells. Chimpanzee DL neurons with pseudotime *t_C_* > *F_HC_*(*t_h_*=1), as well as macaque DL neurons with pseudotime *t_M_* > *F_HM_*(*t_h_*=1), were excluded in following analysis. The second step, namely the alignment step, was then applied to the remaining pseudotime courses of the three species. An ‘end-to-end’ dtw-based alignment, as described above, was used to align the human pseudotimes with pseudotimes of each of the other two species using the human pseudotime course as the template.

### Conserved developmental trajectories from NPCs to neurons in primates

Genes with pseudotime-dependent expression changes in organoids were identified in each of the three species, using the F-test based ANOVA analysis as described above. Those genes with significant pseudotime-dependent expression changes (BH-corrected P<0.05) in all the three species were defined as the genes with universal pseudotime-dependent expression changes, or pseudotime-dependently expressed genes. To estimate the similarities of the expression trajectories among the three species for those genes, Pearson’s correlation coefficient (PCC) was calculated for each gene between each pair of species, across its interpolated expression levels at 50 evenly distributed points along the aligned pseudotimes based on a natural spline regression model (*df*=6). To determine the threshold of a conserved trajectory, we performed 100 pseudotime permutations of pseudocells in the three species. Pairwise PCCs between species were calculated for each of the pseudotime-dependently expressed genes based on the randomized pseudotimes. Minimal PCC of each gene based on each permutation was obtained, and the PCC threshold was determined as the average of the second highest minimal PCC among permutations across all genes of interest. Pseudotime-dependently expressed genes with PCC higher than the threshold between any species pairs were defined as genes with conserved expression pseudotemporal patterns in primates.

### Identification, clustering, and species specificity of differentially expressed genes between humans and chimpanzees

To compare transcriptome changes of the developmental trajectory from cortical neural progenitors to deeper layer neurons between human and chimpanzee, an F-test based comparison was applied to the expression profile along pseudotimes of the two species. In brief, for each gene, a natural spline linear regression model (*df*=6) was constructed for human and chimpanzee pseudocells along the aligned pseudotime course, without discriminating human and chimpanzee samples, and used as the null model (m_0_). The alternative natural spline linear regression model was also constructed, with each species having its own slopes and intercept (m_1_). The residuals of the variation, which cannot be explained by each model, were compared by an F test. Non-ribosomal genes with BH-corrected P<0.01 were identified as differentially expressed genes (DE genes) between human and chimpanzee along the developmental trajectory from cortical neural progenitors to deeper layer neurons (Supplementary Table 7).

To estimate the robustness of the identified differential expression (DE) to the number of used lines, as well as the pseudocell distribution along the pseudotime course, we used a series of replaceable pseudocell sampling procedure with constraints. In brief, in each round of replaceable pseudocell sampling, the candidate pseudocells to be selected are restricted to be those from a certain number of human cell lines. In addition, the subsampling in human pseudocells is performed to recapitulate the pseudocell distribution along the aligned pseudotime of chimpanzee pseudocells, i.e. each of the ten pseudotime bins contains the same number of human and chimpanzee pseudocells. This sampling procedure was performed 100 times for each possible number of human lines, ranging from one to seven. DE analysis, as described above, was applied to compare gene expressions of human pseudocells in each sampling with the chimpanzee pseudocells.

Robust DE genes were determined as DE genes which can be detected in at least 80% of tests performed with replaceable pseudocell samplings with any number of used human cell lines.

A similar strategy was also used to estimate the false positive human-chimpanzee DE genes due to differences between cell lines. In each sampling, two lines were randomly selected as group one, and a certain number of lines, ranging from one to five, were selected from the remaining lines as group two. For each group, pseudocells were randomly sampled from the selected lines to recapitulate the pseudocell distribution along the aligned pseudotime of chimpanzee pseudocells. Such sampling was performed 100 times for each possible number of lines used in group two. The transcriptome trajectory from cortical neural progenitors to deeper layer neurons in macaque organoids was used as the evolutionary outgroup to determine species specificity of the identified human-chimpanzee DE genes. First, the cumulative expression divergences of each gene between human and macaque (*d_HM_*), and between chimpanzee and macaque (*d_CM_*), were calculated. The cumulative expression divergence was calculated by summing up absolute values of average expression differences between species at the 50 pseudotime bins of equal sizes along the aligned pseudotimes. The human-chimpanzee DE of one gene is seen as human-specific if *d*_*HM*_ − *d*_*CM*_ > max (*d*_*HM*_, *d*_*CM*_)/2. Genes with chimpanzee-specific DE were identified in the same way. Genes with human-specific DE were then clustered based on their human-chimpanzee DE along pseudotimes. Average expression differences between human and chimpanzee at each of the 50 pseudotime bins along the pseudotimes was calculated for each gene with human-specific DE (denoted as *d_t_* at pseudotime bin *t*), and then normalized as 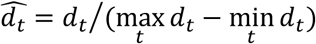. Hierarchical clustering (Ward algorithm) was then used to cluster those genes into nine clusters, with distances between genes calculated as the Euclidean distances between their normalized DE spectrums. Clusters with fewer than five genes were discarded. We annotated genes with human-specific expression patterns using the Homo sapiens Gene Ontology Annotation file (validation date: 21/04/2017) provided by the Gene Ontology Consortium.

### Processing of the Fluidigm C1 based scRNA-seq data of cerebral organoids

In addition to the newly generated Fludigm C1 (SmartSeq2)-based scRNA-seq data, we further retrieved published sequencing data of 786 and 344 single cells from human and chimpanzee cerebral organoids^17, 19^, in the format of FASTQ files from GEO accession numbers GSE75140 and GSE86207 (CMK data set). All the reads were mapped to the human-chimpanzee-macaque consensus genome using STAR (v2.6.1d) with ‘--quantMode’ parameter set to ‘*TranscriptomeSAM*’ and GENCODE v27 annotation provided. Gene expression levels in each cell were quantified as TPM by RSEM (v1.3.1). Additionally, we retrieved the recently published gene expression matrix representing 3211 cells from human and chimpanzee cerebral organoids (excluding redundant cells from GSE75140 and GSE86207) and 4854 cells from human and macaque fetal brains^21^.

Based on the resulting gene expression profile, RSS to the fetal Brainspan data set was calculated as described above for each cell, with 248 genes with significant differential expression between cortical neurons measured by Smart-seq and Smart-seq2 excluded from the references. Distances between organoid cells were calculated as the Pearson’s coefficient distances between RSS of cells. Distances between cells from fetal brains were calculated in the same way. The resulted distance matrices of all organoid cells and fetal brain cells were used as the input to generate tSNE embeddings. kNN-network (k=50) was generated for organoid cells and fetal brain cells separated based on the RSS-based distances, and a Walktrap algorithm for network community identification was applied to identify cell clusters, which were further annotated based on their marker genes. Based on the cell type annotation, the diffusion map analysis, with the RSS profiles as input, was applied to the dorsal forebrain NPCs and neurons in organoids and fetal brains, respectively. The ranks of DC1 were used as the pseudotimes.

To validate the human-chimpanzee differential expression identified in our droplet-based scRNA-seq data using the C1-based cerebral organoid data, the organoid dorsal telencephalon pseudotemporal trajectory was firstly split into ten intervals. In each pseudotemporal interval, the human-chimpanzee DE was calculated as the log2- transformed fold change (log2FC) between the average expression of human and chimpanzee cells in the interval. Here, the CMK data set and other data sets which used a distinct quantification method were processed separately. A similar strategy was also applied to the aligned droplet-based human and chimpanzee pseudotemporal trajectories. Generalized log2-transformed fold change (gLog2FC), defined as the average log2FC across the pseudotemporal intervals, as well as the maximum log2FC across the intervals (mLog2FC), was further calculated for each human-chimpanzee robust DE genes in organoids. A DE gene is seen as being consistent in the two data sets if both gLog2FC and mLog2FC of the C1-based and droplet-based human-chimpanzee comparisons are of the same signs (refer as consistent DE genes). The pseudotemporal intervals with the maximum fold change in the droplet-based and C1-based trajectories were also obtained and compared for the consistent DE genes. This procedure was also applied to compare human-macaque differential expression of the human-specific DE genes along the droplet-based pseudotemporal trajectory and the C1-based fetal brain pseudotemporal trajectory.

### Single-cell and bulk ATAC-seq data generation

Organoids of 2 to 4 months old were washed twice with PBS in a Tissue-Tek Cryomold (Sakura), then embedded in 4% low-melting agarose (Sigma) and sliced into 150 μm sections using a vibrating microtome (Ci 7000 smz, Camden Instruments). Slices were placed on microscope slides containing Differentiation medium with vitamin A (Diff +VA) and inspected under a stereomicroscope to dissect cortical regions. Selected regions were washed twice in 500 μL PBS and incubated at 37°C in 500 μL Accutase (Sigma) plus 0.5 μL DNase I (New England Biolabs) for ∼45 minutes. Trituration was performed for additional mechanical dissociation. Cells were passed through a 30 μm pre-separation filter (Miltenyi Biotec), washed with Diff+VA medium, and spun down at 300 x g (Heraeus Megafuge 40R, Thermo Scientific) for 5 minutes. The cell pellet was resuspended in 200 μl of Diff+VA medium. Cells were viewed under a microscope to ensure a single cell suspension was obtained, and then counted using a Countess Automated Cell Counter (Invitrogen). Single cell suspensions for the early stages of organoid differentiation (iPS cells to neuroepithelium) were obtained as described above.

From the cell suspension, 50,000 cells were used as input for bulk ATAC-seq as described^62^. The remaining cells were diluted to a final concentration of 300 cells/μl and used for microfluidics based single-cell ATAC-seq as described^63^. Briefly, cells were mixed with Suspension Reagent (Fludigm) at a 3:2 ratio and loaded onto a primed medium (10-17 μm) integrated microfluidic circuit (Fludigm) for capturing. Cell capture sites were examined under a microscope and noted for containing 0, 1, or multiple cells. Lysis, transposition, and amplification were performed on the Fluidigm C1 platform. DNA from each cell was transferred to an individual well of a 96-well plate and barcoded with unique combinations of 24 adapter-index i7 and 16 adapter-index i5 primers ^63^. Quantification and library size distribution was assessed on an Agilent 2100 Bioanalyzer using High Sensitivity DNA chips. Excessive primer contamination was removed using SPRIselect (Beckman Coulter Life Sciences) size selection. Up to 192 cells were pooled and sequenced in paired-end, dual-index mode for 50+8+50+8 cycles on one lane of an Illumina HiSeq 2500. A summary of all single-cell experiments can be found in Supplementary Table 1.

### Single-cell and bulk ATAC-seq data processing

Base calling was performed using Bustard (Illumina), adapter trimming with leeHom^64^, and demultiplexing with deML^65^. Reads were aligned to hg19 for human, panTro4 for chimp, and rheMac8 for macaque using bowtie2 with a maximum fragment length of 2000. PCR duplicates were marked and removed using Picard tools (http://broadinstitute.github.io/picard). Samtools^66^ was used to retain properly paired reads with mapping quality greater than 30, while reads mapping to the mitochondrial genome, Y chromosome, and blacklisted genomic regions that show excessively high read mapping, several of which correspond to nuclear mitochondrial DNA segments (identified in Buenrostro et al.^63^ and the ENCODE Project^67^) were removed. For scATAC-seq, single cell BAM files were merged, excluding data from any capture site with 0 or more than 1 cell, to create an aggregated BAM file. Peaks, which represent regions enriched in mapped pair-end sequences, were called using MACS2^68^ with options nomodel, nolambda, keep-dup all, and call-summits. Peak summits were extended by ±250 bp. In the event of overlapping peaks, the peak with the lowest p-value was kept. A single-cell ATAC-seq consensus peak set was obtained by requiring a peak to be accessible in a minimum of 5% of cells. Data visualization was carried out using the Integrative Genomics Viewer (IGV)^69^.

### Enrichment for validated human VISTA enhancers

We overlapped scATAC-seq peaks detected in human cerebral organoids with positive human VISTA enhancers using bedtools *intersect*. For each tissue annotated in the VISTA Enhancer Browser, we counted the number enhancers that did or did not overlap a peak.

We compared these values to the number of all other tissue elements that did or did not overlap a peak. Fisher’s exact tests were performed to determine which tissues’ enhancers had a higher likelihood of being represented. The significance values were corrected for multiple testing using the *qvalue* package in R.

### Cell state identification using single cell ATAC-seq on cerebral organoids and pseudotime estimation

The accessibility at each site in the consensus peak set for every single cell was used to create a count matrix. Cells with fewer than 5000 reads and less than 5% of reads in peaks were filtered out from further analyses. chromVAR^40^ was used to scan the peaks for transcription factor binding motif occurrences, using a curated collection of 1,765 human motifs from the cisBP database, and to identify significantly variable motifs among cells. In addition to TF binding motifs, peaks were scanned for 7-mers. Cell similarity was visualized in a two-dimensional t-SNE plot using the bias-corrected deviations in accessibility for 7-mers with a variability score greater than 1.5.

Each cell’s t-SNE coordinates and the consensus peaks were passed to Cicero^70^ and the densityPeak algorithm was used to identify two clusters of cells Statistically significant differences in TF motif accessibility between the two clusters was calculated using chromVAR, and those motifs corresponding to marker TFs known to distinguish neural progenitors and neurons was used to for cell state identification. Statistically significant differences in accessibility of additional annotations between the two clusters were used to support cell state identities. These annotations included differentially accessible chromatin peaks identified as being enriched in developing mouse brain radial glial cells or excitatory neurons ^71^, as well as accessibility in peaks nearby genes showing pseudotime-dependent expression in cortical neural progenitors or cortical neurons identified as part of this study.

We identified differentially accessible (DA) peaks between the two clusters using the command differentialGeneTest in Cicero. A count matrix was generated with featureCounts ^72^ using the top 250 DA peaks in each cluster. This count matrix was used as input for a diffusion map in order to obtain a pseudotemporal ordering of the cells^73^. Projecting transcription factor binding motif deviation Z-scores on the cells revealed a gradient of known neural progenitor to neuronal markers along the first diffusion map component and we took a cell’s rank along this component as its pseudotime value.

DA peaks identified between the two clusters were used as input test regions for GREAT (version 3.0.0)^74^ with all accessible organoids peaks serving as background regions. We used the default basal plus extension genomic association rule with its default values. All gene ontology (GO) Biological Process terms and their associated hypergeometric p-values were exported. For each term, we plotted the p-value obtained using cluster 1 (identified as NPCs) DA peaks and the p-value obtained using cluster 2 (identified as neurons) DA peaks as input. Terms with a p-value < 0.05 were considered enriched. Informative enriched terms were highlighted based on their significance value in one cell state relative to the other, and for small differences between the cell states when highlighting terms enriched in both.

### Single cell ATAC-seq pseudotime estimation for cells in early states of differentiation and cerebral organoids

Similar to the analysis of the cerebral organoids, we used chromVAR to calculate bias-corrected deviations in accessibility for TF motifs and 7-mers for each cell. Here, we included the scATAC-seq consensus peak sets called in the iPSC, embryoid body, neuroectoderm, and neuroepithelial time points, in addition to the scATAC-seq consensus peak set from the cerebral organoid time point. In the event of overlapping peaks, the peak with strongest signal was retained. Cells with fewer than 5,000 reads and less than 5% of reads in peaks (fraction of reads in peaks, FRiP) were removed from further analyses (Supplementary Table 8). Cell similarity was visualized in a two dimensional t-SNE plot using the bias-corrected deviations in accessibility for 7-mers.

As the cerebral organoid cells’ pseudotimes were previously resolved, we focused on ordering the earlier stages. For this we used Cicero’s differentialGeneTest to identify DA peaks among the iPSC, embryoid body, neuroectoderm, and neuroepithelial time points. A count matrix was generated using the top 250 DA peaks in each time point and used as input for a diffusion map. Projecting TF motif deviation Z-scores of the cells revealed a gradient of pluripotent to more differentiated marker TFs along the first three diffusion map components. We fit a principle curve through the map, and used the pluripotent cells as a starting point to guide the curve. The rank of a cell along this curve was used as its pseudotime. We then added the cerebral organoid cells pseudotime ranks to the end of this earlier stage resolved pseudotime. We used the pheatmap R package to visualize the dynamics of significantly variable motifs across pseudotime.

### Annotation of Accessible Chromatin Peaks

Peaks were linked to an expressed protein-coding gene using the nearest (maximum distance 1 Mb) transcription start site of the canonical transcript as defined by GENCODE (comprehensive gene annotation, release 19). Promoter regions were defined as 1000bp upstream a TSS, and distal regions refer to non-promoter regions. Exon and intron annotations were also obtained from GENCODE (comprehensive gene annotation, release 19). BEDtools^75^ was used to annotate peaks for several evolutionary signatures, including: human accelerated regions^45–47^; selective sweeps compared to great apes^76^ and archaic humans^77^; single nucleotide changes (SNC) in modern humans that happened since the split with great apes and before or after the split with the ancestor of Neandertals and Denisovans, first identified in Prüfer et al. 2014^44^ and updated for this analysis using the most current 1000 Genomes Phase 3 allele frequencies, with a global allele frequency ≥99.5% defined as fixed in all modern humans; small insertions and deletions (up to 5 nucleotides) fixed in modern humans that happened since the split with great apes and before or after the split with the ancestor of Neandertals and Denisovans^78^; and, human deletions that are highly conserved in mammals (hCONDELs, Supplementary Table 10)^48^.

### Identification of genomic regions with differential accessibility between human and chimpanzee organoid neural progenitors and neurons

To compare the chromatin accessibility of NPCs and neurons in cerebral organoids between human and chimpanzee and identify putative regulatory regions that may contribute to transcriptome divergence between human and chimpanzee, we applied a likelihood ratio test based on a generalized linear model with binomial error distribution to each regulatory region identified in human and chimpanzee organoids. More specifically, we identified open chromatin regions in human and chimpanzee organoids separately as described above. To compare an equal number of human and chimpanzee regions, we took the top 77,611 peaks (corresponding to the number of human consensus peaks) in each species and performed reciprocal liftOver, requiring a 50% minimum ratio of based that must remap, in order to identify their orthologous counterparts in the other species. Peaks that successfully lifted over (>99%) were merged using bedtools and re-named (i.e. mergePeak#). Count matrices were generated at these merged peaks in the species own genome, and the matrices were then joined on the common peak name. Considering the higher read coverage in human cells, we subsampled reads in human cells to equalize the medians of total number of reads mapped to the regions of interest in human and chimpanzee. This procedure was applied separately to NPCs and neurons. The resulting count matrices were binarized. We then fitted a generalized linear model for each region across all human and chimpanzee cells, with the accessibility as the response variable and species as the independent variable. Another model with the species variable replaced by a scaling coefficient was also fitted as the null model. The scaling coefficient is fixed to one for human cells and *p*c/*p*h for chimpanzee cells, where *p*c and *p*h are the average accessibility across all regions and all cells in chimpanzee and human, respectively. We compared the two models and got the p-values by using the likelihood ratio test. Regions with BH-corrected P<0.01 were defined as differentially accessible (DA) regions (Supplementary Table 9). This procedure was applied to NPCs and neurons separately to obtain DA regions in the two cell states.

### Functional and evolutionary characterization of genomic regions with differential accessibility

We performed permutations to determine if differentially accessible (DA) peaks were significantly more likely to overlap a given annotation compared to non-differentially accessible (non-DA) peaks. In more detail, we first resized all peaks to an equal length of 500bp and calculated the average accessibility of human and chimp cells in the resized DA and non-DA peaks. Peaks were then placed into average accessibility bins of 5% intervals. Given the number of DA peaks in each accessibility bin, the same number of non-DA peaks was chosen at random from the corresponding accessibility bin. The random set of non-DA peaks was then overlapped with the given annotation using bedtools *intersect*. The random sampling of non-DA peaks and annotation overlap was repeated 2000 times. For each annotation, we counted the number of times a non-DA peak permutation resulted in a higher overlap than what was observed for DA peaks. This number was divided by the number of permutations to determine significance (p<0.05).

We used fixed SNCs, organoid-specific peaks, and linked differentially expressed (DE) genes as annotations. When overlapping peaks with fixed SNCs, we restricted the analysis to include only regions that passed a stringent genome alignability filter (“map35_100%”)^44^, in which SNCs could be called. Organoid-specific peaks were defined as peaks detected in 2-month and 4-month old cerebral organoid stages, but not detected in earlier stages of differentiation (pluripotency to neuroepithelial stages). Cell state-specific peaks were those identified as differentially accessible between NPCs and neurons in either human or chimp.

To study putative effects of fixed SNCs on transcription factor binding in the accessible genomic regions, we used funseq2^79^ to scan and statistically evaluate all possible transcription factor binding motifs created by fixed SNCs in DA peaks. To generate a list of TF motifs lost on the human lineage, we used the human allele as the reference allele and the ancestral allele^44^ as the alternative allele. To generate a list of TF motifs gained on the human lineage, we flipped the state of the reference and alternative allele. This allowed us to directly compare the sequence scores of TF motifs gained or lost in humans. We subtracted the sequence score with the alternative allele from the sequence score with reference allele and performed min-max normalization. Human TF motif gains were plotted as positive values, while human TF motif losses were plotted as negative values. The genomic location of SNCs predicted to alter TF motif binding are provided in Supplementary Table 10. The alteration rate for TF motifs gained in humans was calculated by dividing the number of gains in DA peaks by the number of occurrences of that motif when scanning all organoid accessible peaks using chromVAR and the human genome sequence. The alteration rate for TF motifs lost in humans was calculated by dividing the number of losses in DA peaks by the number of occurrences of that motif when scanning all organoid accessible peaks using chromVAR and the chimpanzee genome sequence. The alteration rates of human TF gains and losses were also calculated per TF family, using TF motif family assignments obtained from^80^.

We used the macaque cerebral organoid scATAC-seq data to determine species specificity of the peaks identified as differentially accessible between human and chimpanzee (Supplementary Table 9). In brief, we counted read coverage of each accessible region we compared between human and chimp which can lift over to the macaque genome in each macaque cell. Regions failed during liftover were seen as inaccessible in all macaque cells. A random sampling of reads in human and chimpanzee cells was applied to equalize median read coverage in the three species. This procedure was applied 100 times and to the two cell states separately. Accessible probability was then calculated for the two cell states in the three species. In human and chimpanzee, averages across the 100 read-subsampling-based estimation were used. The difference of accessible probability between human and macaque (H-M), and that between chimpanzee and macaque (C-M), was then calculated for each human-chimpanzee DA peak in each cell state. The identified DA was considered as human-specific if its H-M difference is at least four times larger than the C-M difference, while its H-M difference is no less than 2%. Similar criteria was also applied to define chimp-specific DA.

To investigate potential biological processes that may be influenced by DA peaks, we used human-chimp DA peaks for each cell state (NPC or neuron) as input test regions for GREAT (version 3.0.0)^74^ with all accessible organoids peaks serving as background regions. This analysis was then carried out the same way as explained above.

### Single-nucleus and bulk RNA-Seq data generation

Cubes were dissected from prefrontal cortex from human, chimpanzee, bonobo and macaque on dry ice aiming for cubes with few curvature to obtain reproducible slicing results. Briefly, the thickness of grey matter at all facets of the cube was measured to obtain a mean gray matter thickness. The mean thickness was divided by 10 to obtain the thickness for each of the segments, whereby each of the segments consisted of several slices at 50 um thickness. Sectioning was performed in a cryostat (Microm, Thermo Fisher), with slices being alternately immersed in Trizol (Invitrogen) for bulk RNA isolation or transferred to a dry tube (low binding) for single nucleus isolation on dry ice. Segments 11 and 12 were collected as well but were considered being derived from white matter of the cortex. Samples were then stored at -80°C until further use.

For nuclei isolation from frozen tissue, all following steps were performed on ice with precooled buffers and centrifugation steps were performed at 4°C. Briefly, tissue was spun down, thawed on ice and 1 ml PBSE (PBS (Gibco), 2 mM EDTA (Life Technologies)) was added to the tissue. The tissue slices were incubated at 4°C on a shaker at 1500 rpm for a total of 45-60 min with trituration steps in between using 1000p and 200p to homogenize the tissue. Generally, segments 1-10 were used for single-nucleus experiments. Two segments were pooled to obtain sufficient material for single nucleus isolation, resulting in 5 segments per individual. To reduce batch effects and increase the number of nuclei per experiment, material from three different individuals (originating from human, chimp/bonobo and macaque respectively) was pooled for each segment. After homogenization, solutions were combined in a 5 ml tube and spun down at 900xg for 5 min. The pellet was resuspended in 1.5 ml PBSE + 1% NP-40 (BioVision), triturated 20 times using 1000p and incubated for 7 min incubation on ice. Samples were then spun down at 900xg for 5 min and resuspend in 1.5 ml PBSE + 1% BSA (Serva) two times. Samples were then spun down again at 900xg for 5 min and resuspended in PBS + 1% BSA. Before sorting, samples were filtered through a 30 um cell filter (Miltenyi Biotec) and stained using DAPI (1:1000, BD Pharmingen). Nuclei were sorted in yield sort mode (BD FACS AriaIII and BD FACS Fusion) based on a defined nuclei population by excluding debris using FSC and SSC and by sorting DAPI positive events. Nuclei were sorted in bulk into 96 well plates and spun down 5 min at 600xg to enrich for nuclei in the pellet.

For each of the pooled samples, 2 lanes on a 10X Chromium microfluidic chip were loaded if feasible, aiming for the maximum possible number of nuclei to be targeted obtained from the sorting. Single-nucleus experiments were performed using the 10X Genomics Single Cell 3’ kit v2 to encapsulate nuclei along with barcode tagged beads, generate and amplify cDNA and to generate sequencing libraries. Each pooled library was barcoded using i7 barcodes provided by 10X Genomics. cDNA and sequencing library quality and quantity were determined using Agilent’s High Sensitivity DNA Assay. Libraries were pooled and sequenced in 150bp paired-end mode on Illumina’s NovaSeq platform as provided in Supplementary table 1.

RNA isolation for bulk-RNA Seq was performed using the Direct-zol 96 RNA kit (Zymo Research) and was quantified using Agilent’s Bioanalyzer RNA 6000 Nano and Pico kit. Libraries were prepared using the NEBNext Ultra Low RNA Library Prep Kit (New England Biolabs). Library quantification was performed using Agilent’s Bioanalyzer DNA 1000 chip kit. All bulk RNA Seq libraries were pooled at equal ratios and sequenced on one lane of an Illumina NovaSeq platform in 150 bp paired-end mode.

### Processing of single-nucleus and bulk RNA-seq data from human, chimpanzee and macaque adult brains

Single-nucleus libraries were demultiplexed based on their i7 index sequences using 10x Cell Ranger (v2.1). Mapping to the human-chimp-macaque consensus genome and generation of count matrices was then performed using the same Cell Ranger, with the GENCODE v27 human annotation provided. Nuclei were assigned to species based on species specific sites using a two-step approach by separating all great ape from macaque nuclei first and subsequently assigning nuclei to either human or chimp/bonobo. Nuclei with a support of less than 80% for either of the groups were removed from further analysis. Moreover, nuclei with less than 200 and more than 6,000 genes detected, so as those with more than 5% detected transcripts being transcribed from mitochondria, were removed from further analyses.

The full single-nucleus RNA-seq data set including all species was further analyzed using Seurat (v3) (Supplementary Table 12). Single-nucleus expression values were normalized and highly variable genes were identified using a variance stabilizing function to detect the top 2000 variable genes (Supplementary Table 11). Data were then integrated by finding corresponding anchors between the species using 30 dimensions. Scaling and principal component analysis were performed using the integrated data. The top 20 principal components were used to identify neighbors of cells and clusters and to visualize the clustering using tSNE embedding. Cluster identities were assigned using unbiased identification using cluster markers by running Seurat’s FindAllMarkers function (Wilcoxon test, min.pct = 0.25, min logFC = 0.25) using non-integrated expression values, known marker genes reported elsewhere (Lake et al., PMID: 27339989, 29227469) and by cell type prediction using Seurat’s TransferData function to anchor to the published Drop-seq based human adult frontal cortex snRNA-seq data (Lake et al. Nature Biotech, PMID: 29227469). Two potential doublet clusters (c11, c19) were excluded from further analysis. For analysis of the major cell classes (excitatory neurons, inhibitory neurons, astrocytes, oligodendrocytes, oligodendrocyte precursor cells, microglia, endothelial cells) subtype clusters were combined and cell type markers recalculated using Seurat’s FindAllMarkers function (Wilcoxon test, min.pct = 0.25, min logFC = 0.25) using non-integrated expression values (Supplementary Table 13).

Since nuclei of the three species have significantly different transcriptome coverage, pseudo-nuclei were constructed for more robust transcriptome measurement, as well as for more fair and efficient comparison, using a similar procedure as described above to generate pseudocells, under the constraint of merging only nuclei from the same segment of the same sample and grouped in the same cell cluster. The probabilities of nuclei selected as pseudo-nuclei seed were 1/13 for human, 1/8 for chimpanzee and 1/10 for macaque.

Reads of the bulk RNA-seq samples were mapped to the human-chimpanzee-macaque consensus genome using STAR (v2.6.1d). The Python utility hiseq-count was used to count the numbers of uniquely mapped reads of genes annotated in GENCODE v27 human annotations. DESeq2 was used for normalization and retrieving FPKM as the expression level measurement.

To determine the laminar origin of each segment, genes with segment-dependent expression were firstly screened for each cortical cube. In brief, an ANCOVA analysis was applied to compare two models: the natural spline (df = 6) linear model with log10- transformed FPKM as the response and the segment order as the variable; the null model of expression values without any linear relationship with segments. For each of the resulted gene, its enriched segments in the cube were identified, as the segments with the gene’s expression at least one standard deviation higher than the mean across segments. Genes with enriched expression at each segment were then overlapped with the layer markers identified in ^9^. Segments with enriched genes significantly overlapping with markers of only one layer were seen as pure-layer original, others were seen as mixture of multiple layers. For each mixture segment, a quadratic-programming-based transcriptome deconvolution ^81^ was applied to determine the relative contribution of the enriched layers. A layer index was then obtained for each segment, as the average layers weighted by contributed proportion of the enriched layers.

### Estimation of cell type distribution across cortical layers and gene expression patterns in neurons across cortical layers

To estimate the cell type composition of each layer, nuclei from each sample were randomly assign to one layer, based on the layer mixture proportions estimated above. The proportion of each of the six major cell classes: excitatory neurons, inhibitory neurons, astrocytes, oligodendrocytes, oligodendrocyte precursor cells (OPCs), microglia and endothelial cells, was then calculated for nuclei assigned to each layer in human. This procedure was repeated 100 times, with the resulted average as the final estimation. The laminar distribution of each cell cluster was also estimated based on the described procedure. In addition, a subsampling procedure with replaceable manner of the same number of nuclei (n = 200) from each layer was further applied to each of the 100 nuclei layer random assignment to control differences on the detected nuclei number of each layer. To get more precise estimation of layer origins on the nuclei level for excitatory and inhibitory neurons, both of which show a distinct layer distribution pattern across different subtypes, we trained an elastic net linear regression model (alpha = 0.5) on excitatory and inhibitory neurons separately, with the sample layer indices as the training response and expression levels of the highly variable genes as the variables. To enhance model robustness, pseudo-nuclei from all the three species together were used for model trainings. The trained models were then applied to the excitatory and inhibitory pseudo-nuclei again. The predicted layer indices were used as the estimated relative laminar location of the pseudo-nuclei. The projection of the predicted layer indices to layers were done by averaging expression patterns of markers of different layers^9^.

### Differential expression analysis between human and chimpanzee cell types in adult brains and determination of their species-specificity

Due to the sparse nature of the snRNA-seq data and the unequal coverage of nuclei from different species, commonly used statistical test for differential expression analysis (e.g. Wilcoxon’s rank sum test) failed to provide reliable estimation of DE, even with the state-of-art VST normalization methods^82^. As detection rates of genes are correlated with their expression levels^82^, we therefore compared gene expression levels of the same cell type in human and chimpanzee by comparing their detection rates, using a GLM-ANCOVA analysis similar to the one described above to identify genomic regions with differential accessibility. In brief, the pseudo-nuclei expression matrix was binarized. A binomial GLM model was trained for each gene, with its detection as the response variable and species of pseudo-nuclei as the independent variable. This model was compared to the null model with the species variable replaced by a scaling coefficient. The scaling coefficient is fixed to one for human pseudo-nuclei and p_c_/p_h_ for chimpanzee pseudo-nuclei, where p_c_ and p_h_ are the average detected gene numbers across pseudo-nuclei involved in the test in chimpanzee and human, respectively.

While the described DE test was applied to four cell classes with sufficient numbers of pseudo-nuclei: excitatory neurons, inhibitory neurons, astrocytes and oligodendrocytes, the heterogeneity within the two neuron types, as well as their uneven distributions in human and chimpanzee, needed to be considered. A subsampling procedure with replaceable manner was therefore applied. In every subsampling, an equal number of pseudo-nuclei (n = 200) from each species were sampled, with pseudonuclei in clusters annotated as the cell class of interest sharing equal probability being selected. The described DE test was then applied to the sampled nuclei of this cell class. This subsampling procedure was repeated for 100 times, and DE genes of each cell class were defined as genes with significant DE (BH-corrected P<0.005) in at least 80 times of the subsampling. Additional filtering was then applied, requiring the same direction of human-chimpanzee difference on detection rates and VST-normalized expression values.

Macaque pseudo-nuclei were then introduced to investigate species specificity of the identified DE. Similar procedure sampling the same number of pseudo-nuclei from clusters annotated to be the same cell class was repeated 100 times to the macaque pseudo-nuclei. For each sampling, average VST-normalized expression values were calculated for each cell class in human, chimpanzee and macaque, with which differences between human and macaque (*d_HM_*), as well as between chimpanzee and macaque (*d_CM_*), were calculated. The identified human-chimpanzee DE was defined as human-specific if |*d_HM_*| > 4 * |*d_HC_*|. Genes with chimpanzee-specific DE were identified in the same way (Supplementary Table 14).

## EXTENDED DATA FIGURES

**Extended Data Figure 1.**
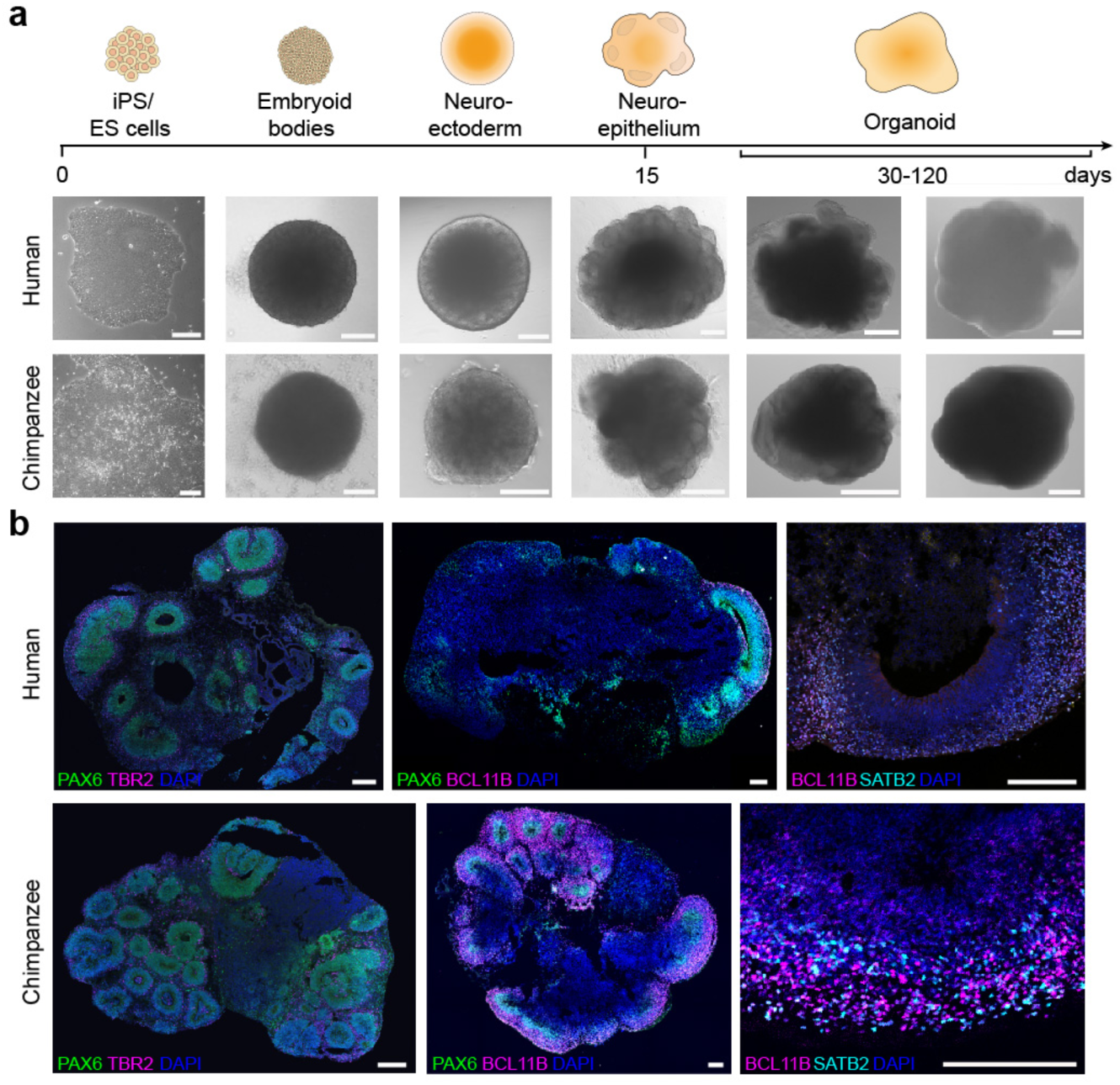
Differentiation and immunohistochemical characterization of human and chimpanzee cerebral organoids. (a) Phase contrast (iPSC stage – neuroepithelium, scale bar 200 μm; H9 for human, SandraA for chimpanzee) and bright field images (organoid stages, scale bar 1 mm; H9 and Wibj2 for human, JoC and SandraA for chimpanzee) of different stages of organoid development for human and chimpanzee organoid differentiation. (b) Immunohistochemical stainings of human (Sc102a1 and 409b2) and chimpanzee (all SandraA) organoids reveal proper formation of cortical-like regions (scale bar 200 μm).

**Extended Data Figure 2:**
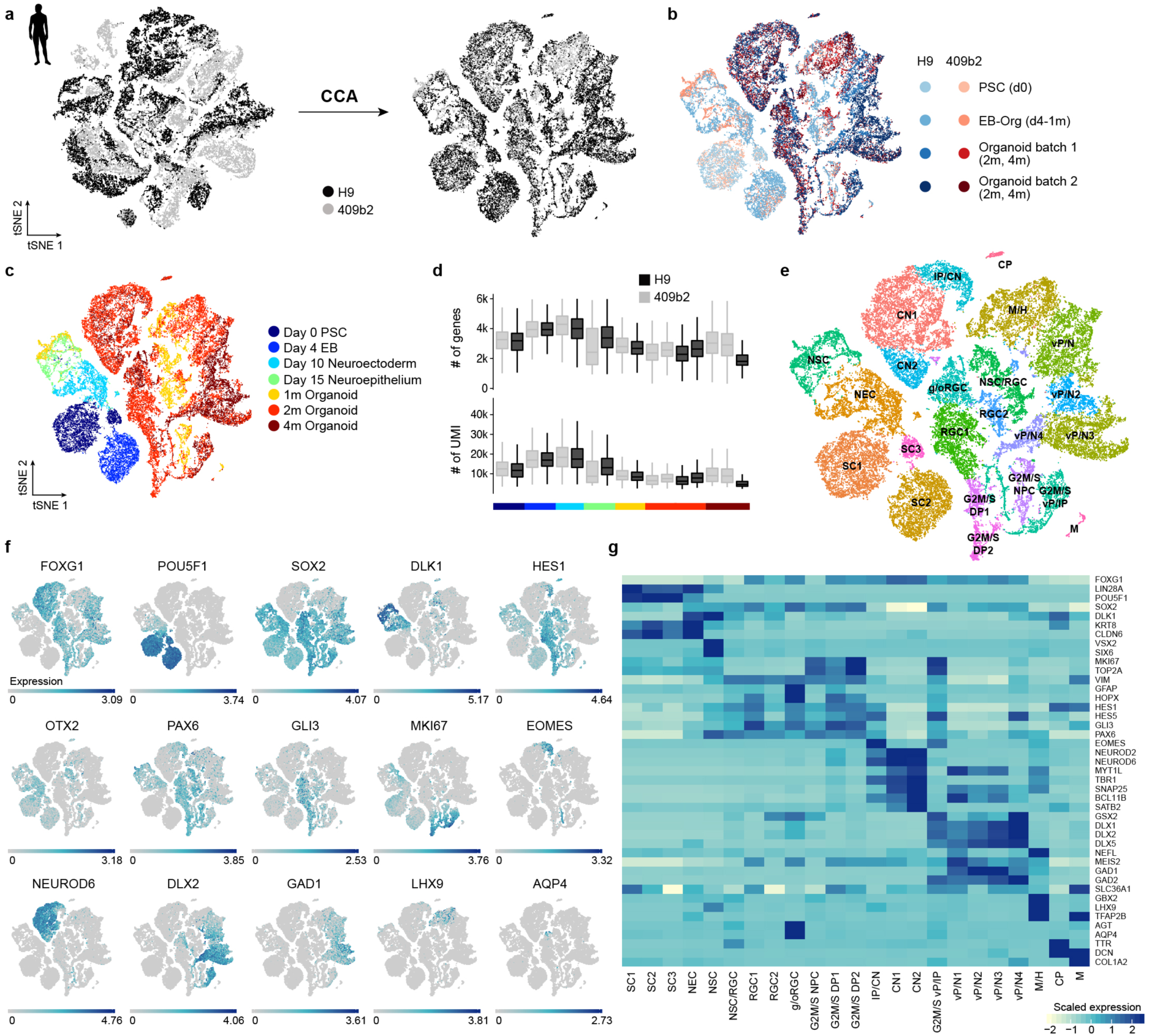
Heterogeneity analysis during human cerebral organoid development from pluripotency. (a) Cells from different human cell lines were integrated using canonical correlation analysis and visualized using t-stochastic neighbor embedding (tSNE). (b) tSNE color coded based on cell line and batch. (c) tSNE colored based on time point. Heterogeneity analysis was performed on combined cells from day 0 of differentiation to 4 month old organoids for iPSC and ESC-derived cells. (d) Distribution of number of genes and UMIs for different time points and cell lines. (e) Clustering was performed using the top 20 principal components as input for tSNE and cluster names were assigned based on expression of cluster marker genes and known marker genes. SC stem cells, NEC – neuroectoderm-like cells, NSC – neural stem cells, (g/o)RGC – (gliogenic/outer) radial glia cells, G2M/S NPC – neural progenitor cells in G2M/S phase, G2M/S DP – dorsal progenitor cells in G2M/S phase, IP – intermediate progenitor, CN – cortical neurons, G2M/S vP – ventral progenitors in G2M/S phase, M/H – midbrain/hindbrain, CP – choroid plexus, M – mesenchymal-like cells. (f) tSNE plot colored with respect to expression of selected marker genes based on non-integrated expression values. (g) Heatmap showing averaged cluster expression for representative marker genes for clusters ordered according to their differentiation time from early to later stages and regional identity from dorsal to ventral forebrain and non-forebrain cells.

**Extended Data Figure 3:**
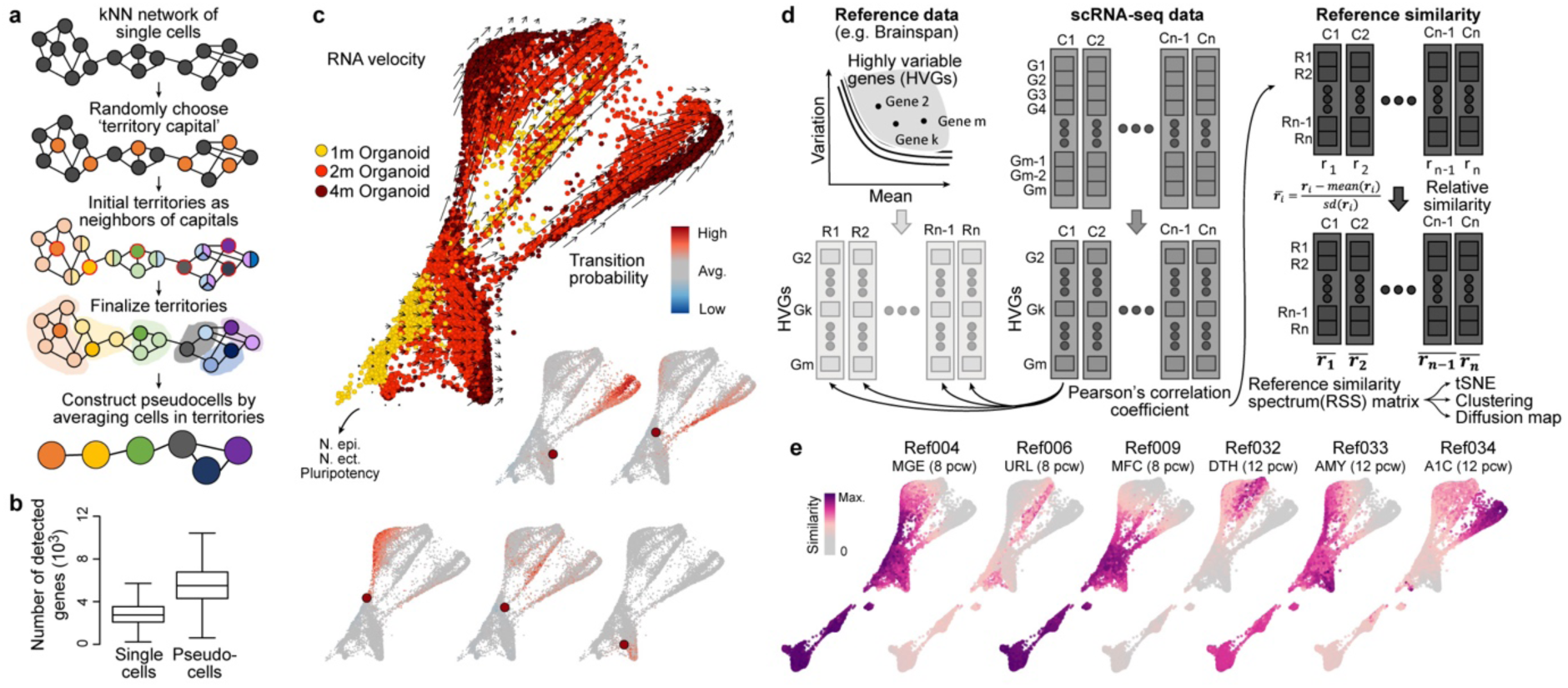
Pseudocell construction, monitoring differentiation using RNA velocity, and reference similarity spectrum calculation. (a) Pseudocells are constructed based on kNN networks of single cells. Random cells in the network are selected as seeds or ‘territory capitals’, with their neighbors as initial members belonging to the territories; cells assigned to multiple territories are randomly assigned to one to finalize pseudocell territories, with the transcriptome of each pseudocell calculated as the average transcriptome of cells in its territory. (b) Boxplots (box, interquartile range (IQR); whisker, 1.5*IQR) showing the number of detected genes significantly increased in the constructed pseudocells compared to single cells. (c) RNA velocity analysis applied to the constructed pseudocells suggests the neurogenesis trajectories of the three different neuronal branches of cortical neurons, ventral forebrain neurons and non-forebrain neurons. Velocity transition probabilities are shown for five example neural progenitor pseudocells (red). (d) Reference Similarity Spectrum (RSS) is calculated for each cell or pseudocell against RNA-seq data of 237 fetal macrodissected human brain samples in Allen Brain Atlas (Brainspan). Pearson correlation coefficients are calculated across 2,256 highly variable genes of the Brainspan data set. Correlations of one cell to different reference samples are normalized by z-transformation, which represents its normalized similarity to reference samples. The normalized correlation to each reference sample across cells represent similarity patterns of a cell to each reference sample, while the resulting normalized RSS of each cell is seen as its dimension-reduced representation. (e) SPRING plot with pseudocells colored by normalized similarities to six Brainspan reference samples.

**Extended Data Figure 4:**
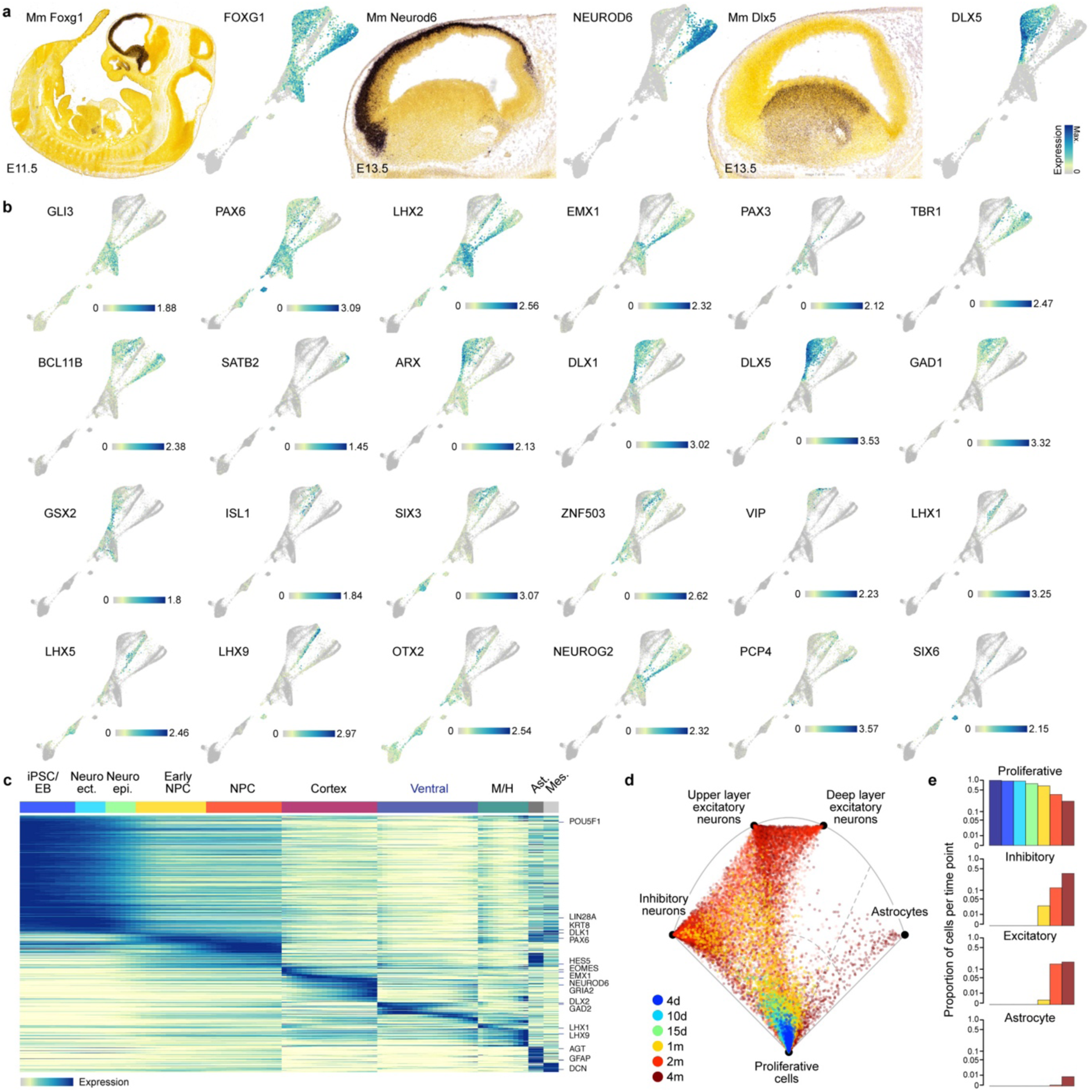
Marker gene expression during the human organoid time course. (a) In situ hybridization images from the Allen Developing Mouse Brain Atlas showing expression of Foxg1, Neurod6, and Dlx5 in the mouse developing forebrain, and human whole-trajectory SPRING plots colored by the corresponding genes. (b) Whole-trajectory SPRING plot colored by marker gene expression. (c) Pseudotemporal expression of example genes with pseudotemporal expression changes in the whole human cerebral organoid developmental trajectory. (d) Umbrella plot showing the similarity of each organoid cell to a cell “prototype” generated from a reference scRNA-seq cell atlas of the human fetal cortex^27^. (e) Plots show the proportion of organoid cells per time point that match a reference prototype.

**Extended Data Figure 5:**
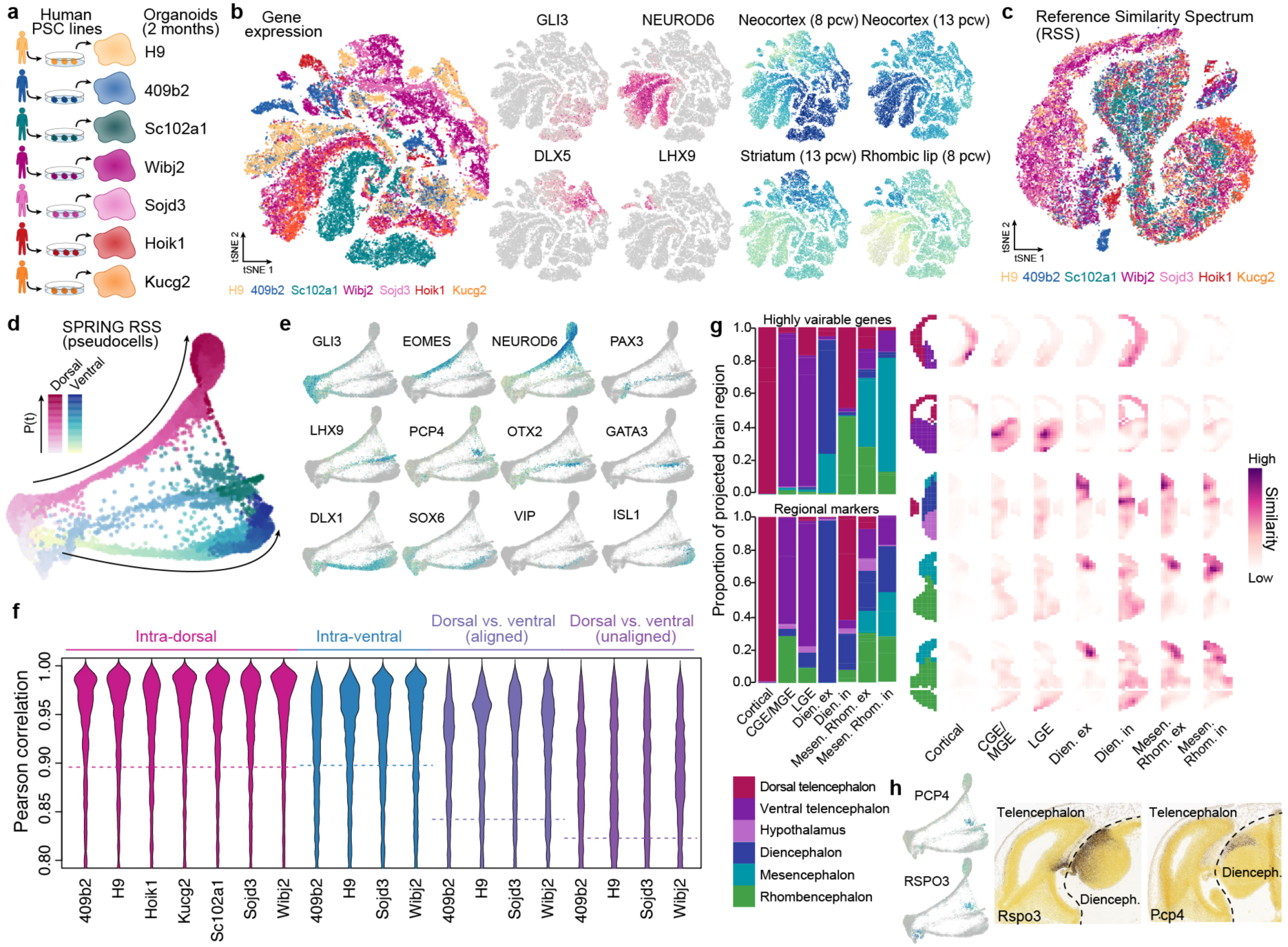
Analysis of human cerebral organoid single-cell transcriptomes from 7 individuals. (a) scRNA-seq was performed on 2-month old cerebral organoids from human ESC and six iPSC lines. (b) All data were combined and cell heterogeneity was assessed using t-distributed stochastic neighbor embedding (tSNE) with the top-20 principal components (PCs) as the input. Cells are also colored by marker gene expression and RSS. (c) tSNE plot with RSS against Brainspan fetal reference data as the input (RSS-tSNE), colored by cell lines. Cells from different lines are well integrated. (d) SPRING plot of 2-month old human organoid pseudocells, colored by neuronal trajectory branches and pseudotimes. (e) SPRING plot of 2-month old human organoid cells, colored by marker gene expression. (f) Correlations of expression trajectories of genes with pseudotime-dependent expression patterns between cortical cells from each line to the others (pink), ventral cells from each line to others (blue), and cortical and ventral cells from the same lines after or before aligning the cortical and ventral pseudotimes (purple). (g) Spatial location inference of neuron subtypes in human cerebral organoids. (Left) Barplots show proportion of cells of each cell type which show highest gene expression pattern similarity to the average expression patterns in different structures, based on the processed in situ hybridization image data (E13.5) provided in Developing Mouse Brain database of Allen Brain Atlas. Expression similarity was calculated based on highly variable genes of the scRNA-seq data (top), or regional markers defined with the ISH data (bottom left). (Right) Correlation patterns of average regional marker gene expression of each neuron subtype to voxels in five example sections (E13.5), as well as the structural annotation of the sections. (h) Expression of two marker genes of Diencephalon inhibitory neurons (PCP4, RSPO3) in the SPRING embeddings, and their spatial expression patterns in E13.5 mouse brain (data from Allen Brain Atlas).

**Extended Data Figure 6:**
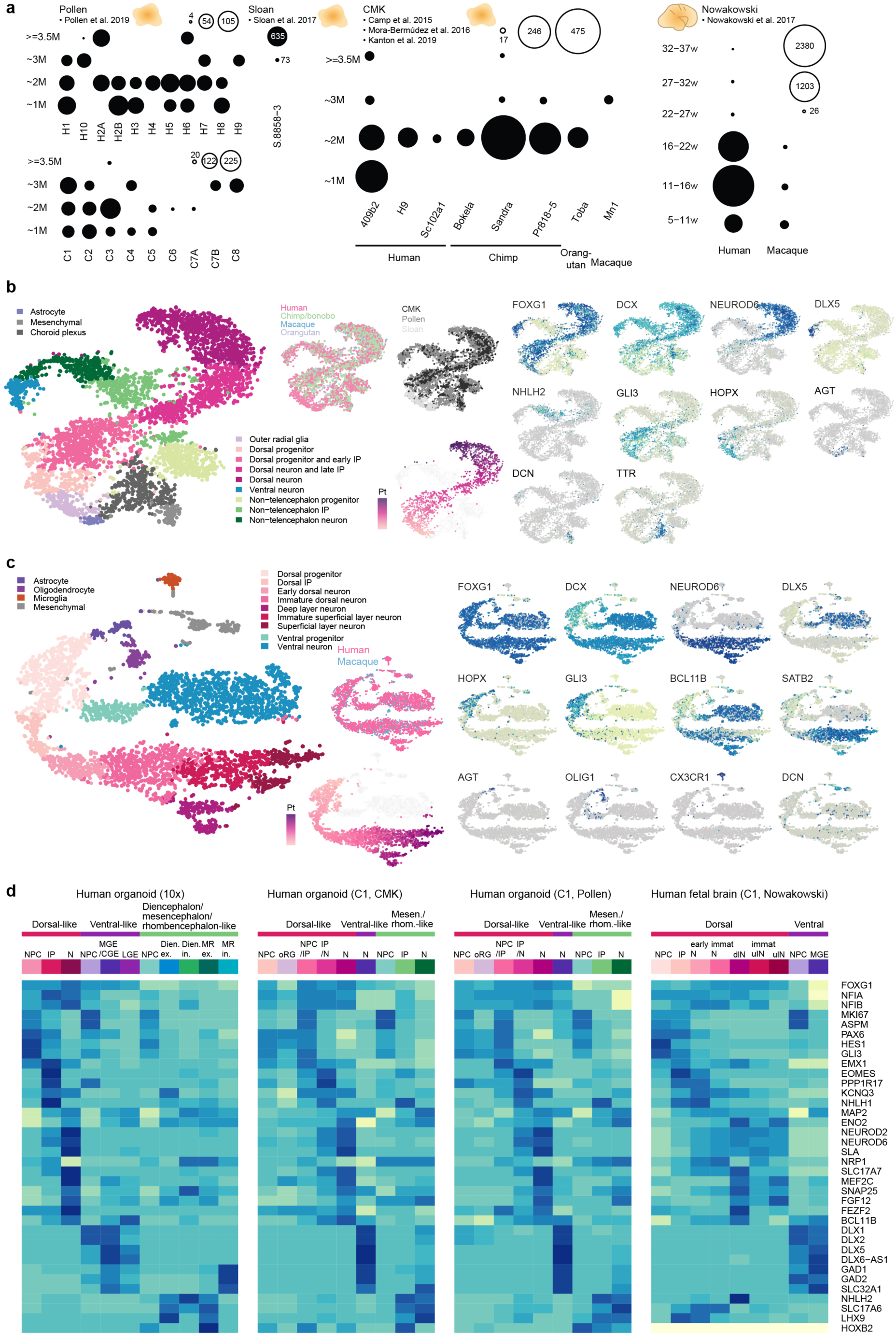
Supplementary analysis on cell type heterogeneity of cerebral organoids and fetal cortical tissues based on scRNA-seq data from Fluidigm C1. (a) Overview of the Fluidigm C1 scRNA-seq data. Each dot represents a cerebral organoid or fetal brain sample from one cell line or species at a certain age, with its size showing the number of cells measured. The left panel shows organoid sample information as published in Pollen et al. 2019 (excluding redundant cells from Camp et al. 2015 and Mora-Bermudez et al. 2016), including the data initially published in Sloan et al. 2017. The middle panel shows organoid sample information generated in Camp et al. 2015, Mora-Bermudez et al. 2016, and in this manuscript. The right panel shows fetal prefrontal cortex sample information reported in Nowakowski et al. 2017. (b) All cerebral organoid data were combined and cell heterogeneity was assessed using t-distributed stochastic neighbor embedding (tSNE) with the reference similarity spectrum (RSS) profiles to the fetal Brainspan data as the input. Cells are colored by cell type/cluster, species, institutions generating the data, dorsal trajectory pseudotimes, and marker gene expression. (c) tSNE plots for all fetal brain data to assess cell heterogeneity, with the RSS profiles to the fetal Brainspan references as the input. Cells are colored by cell type/cluster, species, dorsal excitatory neuron trajectory pseudotimes, and marker gene expression (d) Heatmap showing marker gene expression patterns across different cell types in the droplet-based organoid scRNA-seq data generated in this manuscript and the above described C1-based scRNA-seq data.

**Extended Data Figure 7:**
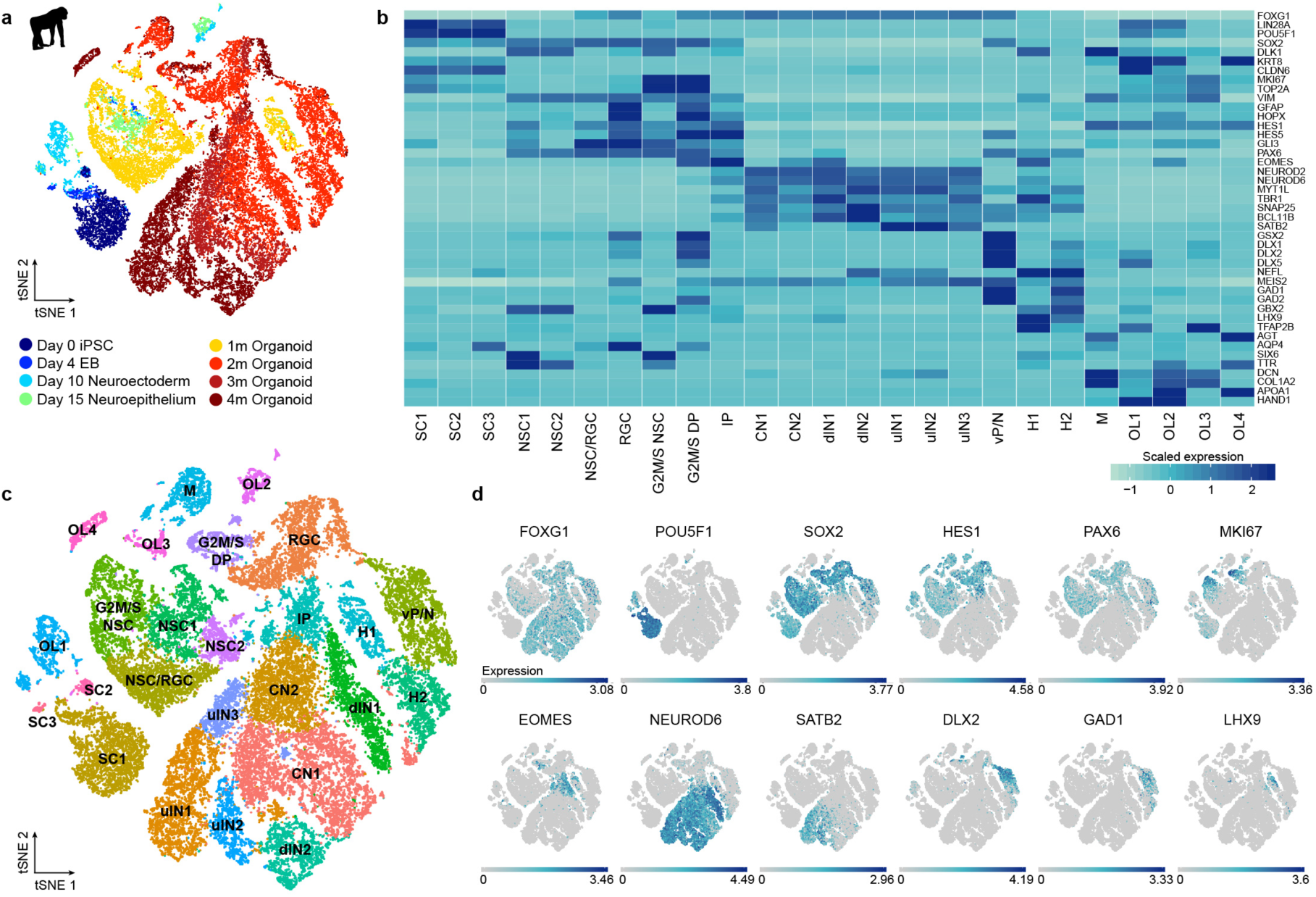
Heterogeneity analysis during chimp cerebral organoid development from pluripotency. (a) Heterogeneity analysis for iPSC-derived chimpanzee cells from day 0 of differentiation to 4 months of organoid development for one cell line. (b) Heatmap visualizing averaged cluster expression for marker genes with columns ordered based on differentiation progress from early to late time points and regional identity sorted from dorsal to ventral forebrain to non-forebrain cells and non-ectodermal-derived cells. (c) Cluster identification and t-stochastic neighbor embedding using the top 15 principal components for clustering. Cluster assignment was based on cluster markers as well as expression patterns of known marker genes. SC – stem cells, NSC - neural stem cells, RGC – radial glia cells, G2M/S DP – dorsal progenitors in G2M/S phase, IP – intermediate progenitors, CN – cortical neuron, dlN – deep layer neuron, ulN – upper layer neurons, vP/N – ventral progenitor/neuron, H – hindbrain, M – mesenchymal-like cells, OL – off lineage cells, MIC - microglia. (d) tSNE plots colored based on gene expression of representative marker genes used to assign cluster identities.

**Extended Data Figure 8:**
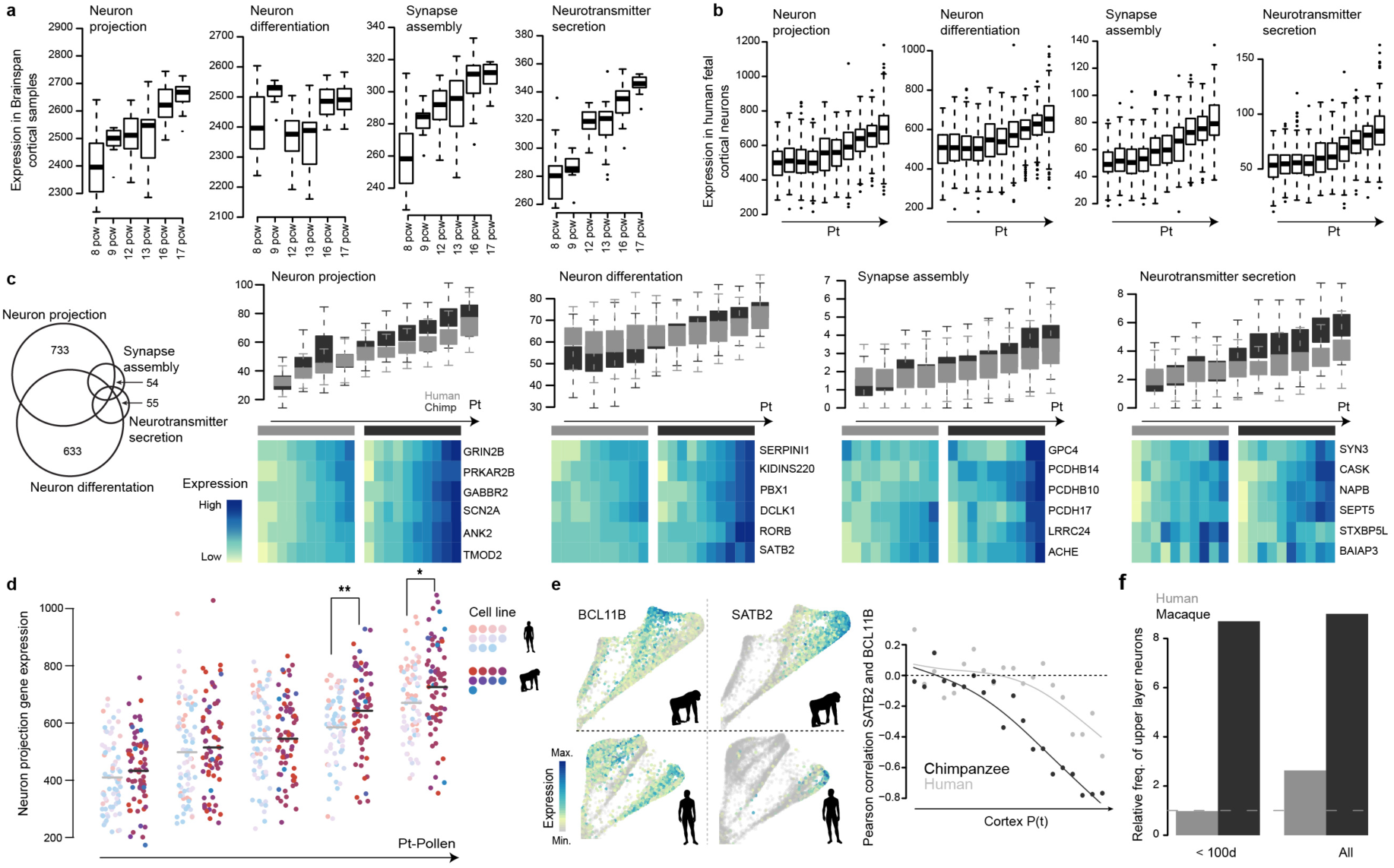
Supplementary analysis on timing difference of neuron maturation in human and chimpanzee cerebral organoids. (a) Boxplots (interquartile range with minimum and maximum, outliers removed) showing sum expression levels (in RPKM) of genes with Gene Ontology annotation ‘neuron projection’, ‘neuron differentiation’, ‘synapse assembly’ and ‘neurotransmitter secretion’ in Brainspan fetal cortical samples aged from 8 post conception weeks (pcw) to 17 pcw. (b) Boxplots showing sum expression levels of the same gene lists in fetal human dorsal excitatory neurons along the estimated developmental pseudotimes (Nowakowski et al. 2017 data set). (c) Boxplots showing sum expression levels of genes with specific annotation to only one of the four GO terms in human and chimpanzee along the cortical pseudotimes. Heatmaps showing expression of example genes from the GO terms for human and chimp along pseudotime bins. The Venn diagram on the left shows the overlap of genes related to the four GO terms. (d) Distribution of neuron projection scores of human and chimpanzee cortical cells reported in Pollen et al. 2019 along the cortical pseudotimes. Each dot represents one cell, and is colored by the organoid cell line. Light colors represent human cell lines, and dark colors represent chimpanzee ones. Two-sided Wilcoxon’s rank sum test (*: P<0.1; **: P<0.01). (e) Observed timing difference of upper-deeper layer specification in human and chimpanzee cerebral organoids from 10X genomics data generated in this study. The left panel shows expression of cortical deep (BCL11B, left) and upper (SATB2, right) layer marker genes projected onto the chimpanzee (top) and human (bottom) SPRING plot. BCL11B and SATB2 become anti-correlated in their pseudotemporal expression profile in both human and chimpanzee (right), while the onset of anticorrelation happens earlier in chimpanzee than in human. (f) Abundance of upper layer neurons relative to deeper layer neurons in human and macaque fetal prefrontal cortex samples in Nowakowski et al. 2017 grouped by early time points (<100 days old) or all time points combined.

**Extended Data Figure 9:**
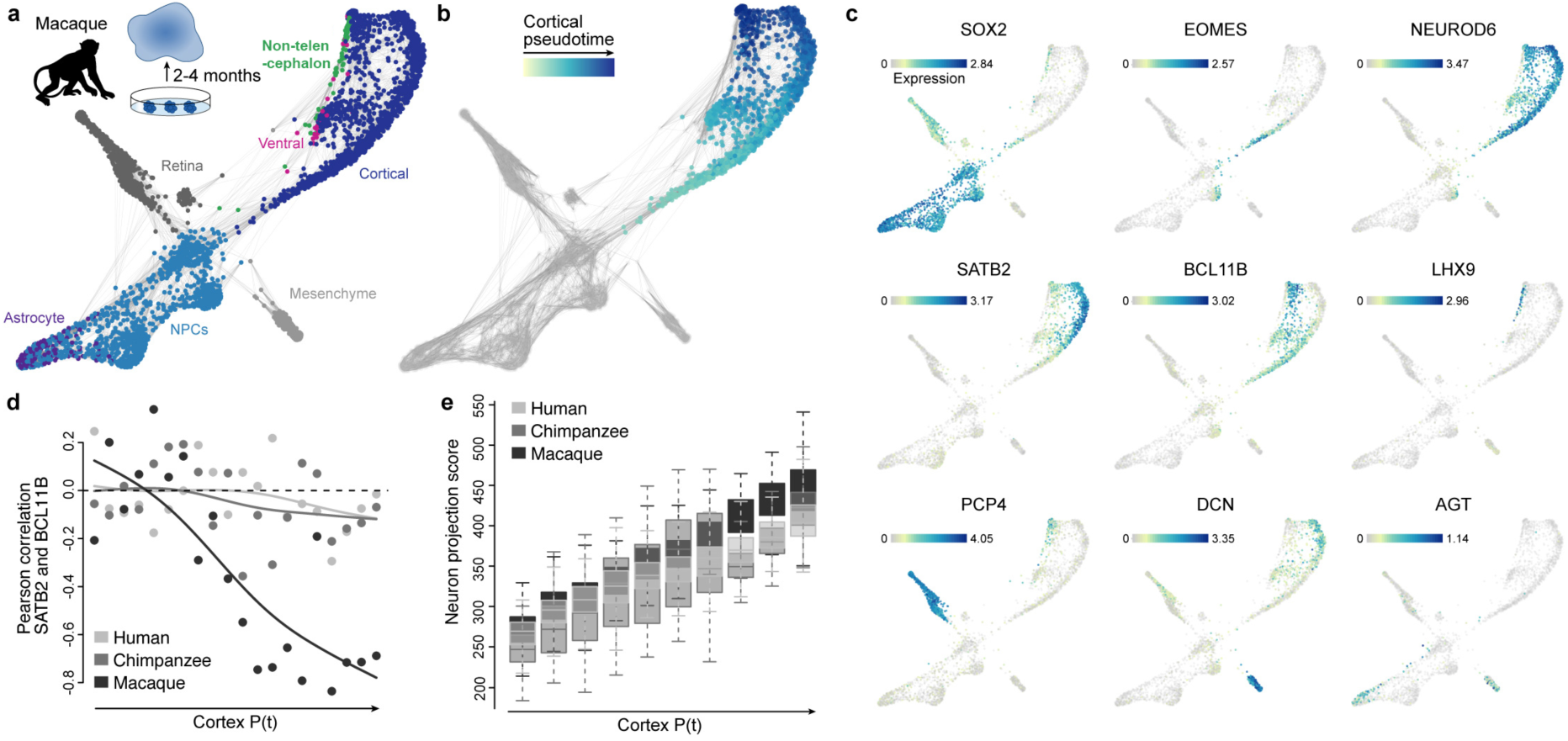
Single-cell RNA-seq analysis of macaque cerebral organoids. (a) scRNA-seq was performed on 2 to 4-month cerebral organoids from a macaque iPSC line. The SPRING plot of pseudocells was constructed with the top 20 PCs as the input. The heterogeneity analysis suggests multiple cell types in the macaque organoids, including cortical neurons, NPCs, astrocytes and other cell types such as retina and mesenchyme-like cells. (b) SPRING plot colored by pseudotimes of cortical pseudocells, which are the pseudocells’ quantiles of DC1 of the cortical pseudocells diffusion map. (c) SPRING plot colored by marker gene expression. (d) The onset of anti-correlation between SATB2 and BCL11B occurs earlier along the macaque pseudotime, relative to human and chimpanzee, when focusing on the 2-month cerebral organoids. (e) Boxplots (box, interquartile range (IQR); whisker, 1.5*IQR) showing the neuron projection scores in human, chimpanzee and macaque along the unaligned cortical pseudotimes.

**Extended Data Figure 10:**
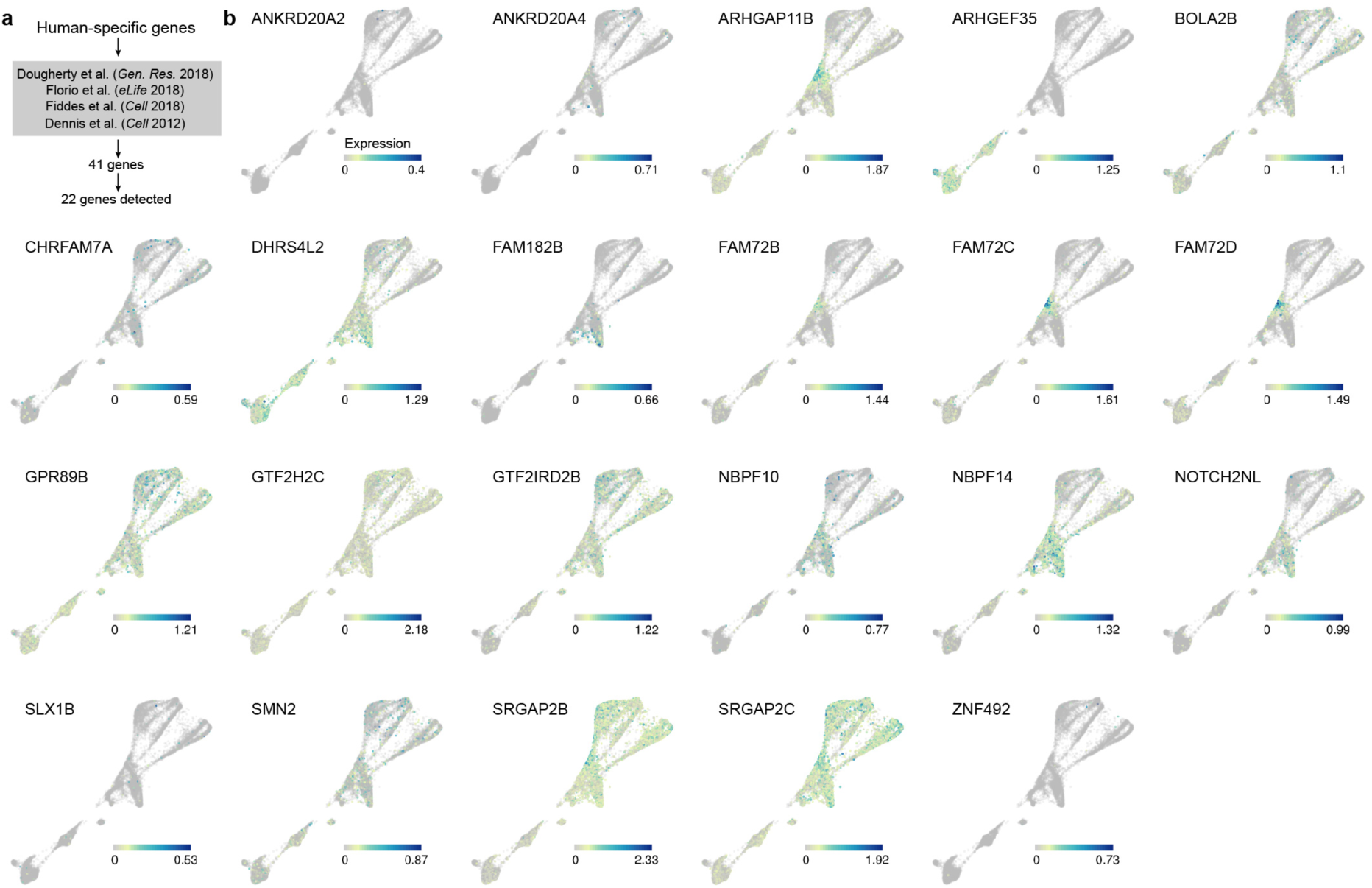
Expression of human-specific genes along human organoid development from pluripotency. (a) Reported human-specific genes were collected from four previous publications, and the expression of these genes was analyzed along the human organoid developmental trajectory from pluripotency to 4 month old organoids. (b) SPRING plot colored by expression of the human-specific genes.

**Extended Data Figure 11:**
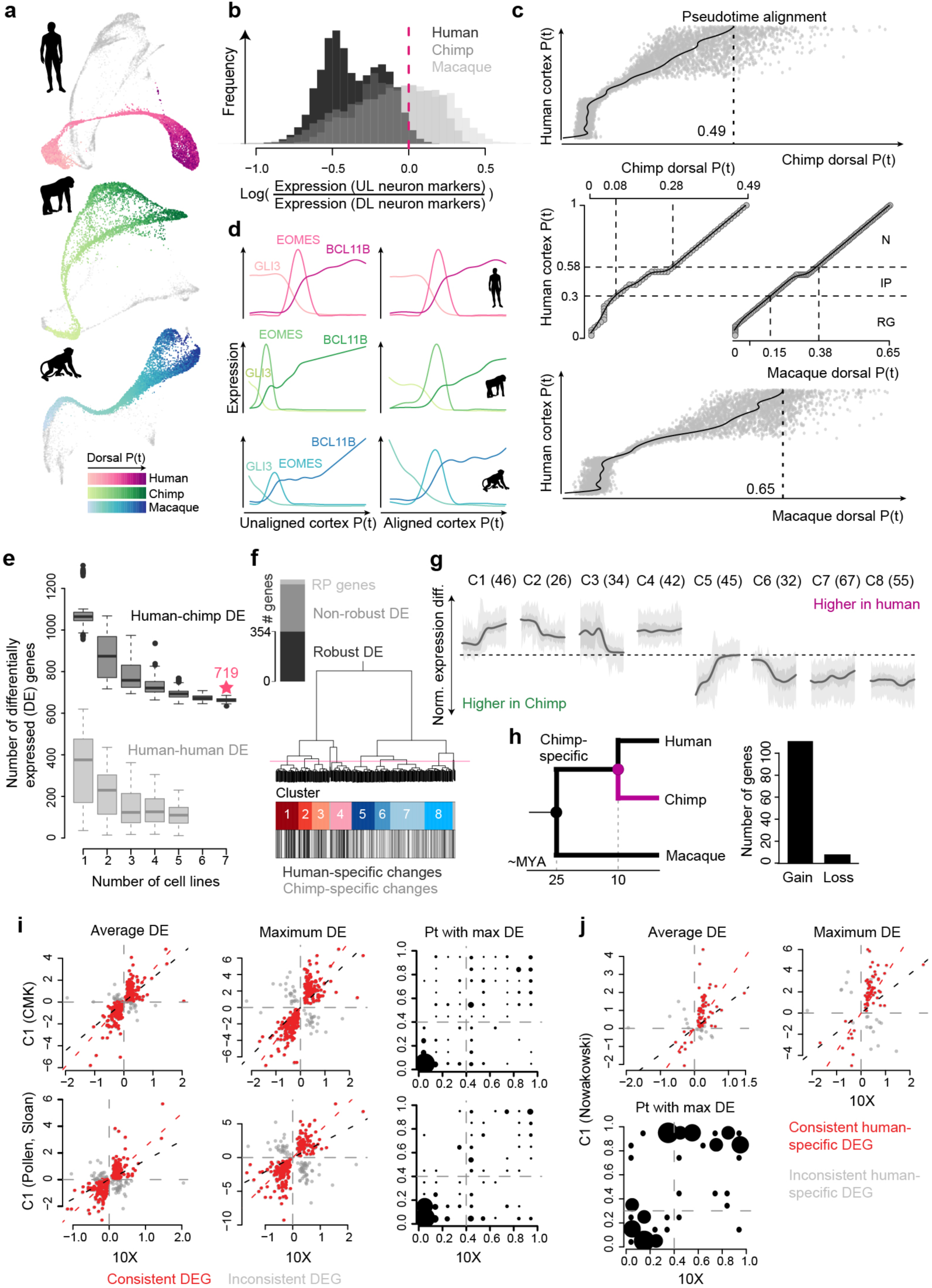
Pseudotime alignment between primates and differential expression between human and chimpanzee. (a) SPRING plots visualizing the kNN networks of human and chimpanzee pseudocells, and macaque cells, which represent neural progenitors (NPC) and neurons of different brain regions. Cortical NPCs and neurons are colored by their pseudotimes. (b) Ratios of upper layer (UL) to deeper layer (DL) neuron marker expression in human (black), chimpanzee (dark grey) and macaque (light grey) organoids. The dashed line indicates the cutoff applied to human pseudocells to filter out those representing UL neurons. (c) Truncated dynamic time warping (dtw)-based alignment was applied to align human, chimpanzee and macaque cortical pseudotime courses. Two support vector regression models were trained to predict chimpanzee (upper) and macaque (lower) pseudotimes of human pseudocells. A constrained B-splines regression model was fitted to determine the trimming point at the chimpanzee and macaque pseudotime courses, respectively. An end-to-end dtw-based alignment was applied to the human pseudotime course to the trimmed chimpanzee and macaque pseudotime courses for the final alignments (middle). (d) Pseudotemporal expression profiles of GLI3, EOMES and BCL11B along the human, chimpanzee and macaque cortical pseudotimes, before (left) and after (right) the pseudotime alignment procedures. (e) Robustness and false positive rate of differential pseudotemporal expression between human and chimpanzee based on the number of cell lines involved in the analysis with constrained replaceable pseudocell subsampling. In each subsampling, pseudocells representing cells from a certain number of human lines were sampled in a replaceable manner to recapitulate pseudocell distribution along pseudotime course of the chimpanzee pseudocells. Differential expression (DE) analysis was applied to compare all chimpanzee pseudocells and the sampled human pseudocells to estimate robustness to cell line numbers (dark grey boxes), and to compare sampled human pseudocells to human pseudocells from another two lines sampled with the same procedure to estimate false positive rate (light grey boxes). Boxplots: box, interquartile range (IQR); line, 1.5*IQR; dots, outliers. (f) Robustly detected human-chimpanzee DE genes (robust DE genes) are defined as the non-ribosomal genes which were detected as DE genes in at least 80% of the subsampling-based human-chimpanzee DE analysis using any number of human lines (black). The dendrogram shows the hierarchical clustering of robust DE genes, based on their human-chimpanzee pseudotemporal DE patterns along the aligned pseudotimes of cortical organoid pseudocells, resulting in eight clusters of robust DE genes. (g) Pseudotemporal differential expression patterns between human and chimpanzee of the eight clusters of genes along the pseudotimes of cortical organoid pseudocells with 50% and 95% confidence intervals shown in dark and light grey, respectively. (h) Number of differentially expressed genes in chimpanzee vs. human and macaque comparison grouped by gain or loss of expression in chimpanzees. A gain of expression specifically in chimpanzees is more likely than a loss of expression pattern conserved in the other primates. (i) Comparison of the reported human-chimpanzee pseudotemporal DE based on 10X Genomics data with the Fluidigm C1-based scRNA-seq data of human and chimpanzee cerebral organoids. The two rows show the results based on C1 data generated in this manuscript and combined with data from Camp et al. 2015^17^, Mora-Bermudez et al. 2016^19^, Pollen et al. 2019^21^. The first two columns show estimated human-chimpanzee DE directionality and magnitude in the reported droplet-based scRNA-seq data and the C1-based measurement, with the first column presenting the generalized DE along the whole cortical pseudotimes, and the second column presenting the maximum DE along the pseudotimes. The red dots represent consistent DE genes, which have consistent DE directionalities in the two data sets. The right panel shows pseudotime intervals with the largest human-chimpanzee DE in the two data sets in comparison to the consistent DE genes. Dot sizes represent frequencies. (j) Comparison of the estimated human-macaque DE directionality and magnitude of the human-specific DE genes using human and macaque fetal prefrontal cortex scRNA-seq data^21, 27^.

**Extended Data Figure 12:**
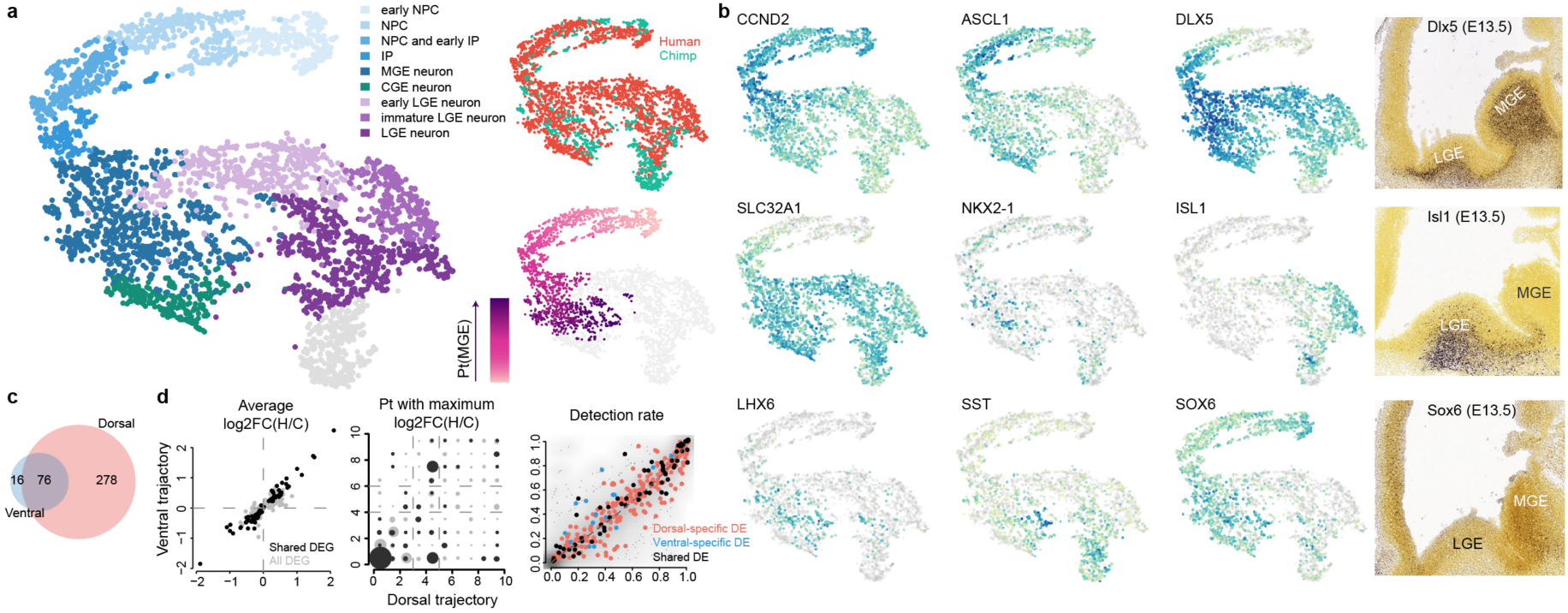
Organoid ventral forebrain cell heterogeneity and differential expression between human and chimpanzee. (a) Ventral forebrain cell heterogeneity in organoids was investigated by tSNE embeddings, with RSS profiles of human and chimpanzee ventral pseudocells combined as the input. Pseudocell clusters were annotated based on marker gene expression. Pseudocells were also colored by species and diffusion map based on medial ganglionic eminence (MGE) neuron developmental pseudotimes. (b) tSNE plots colored by marker gene expression, and in situ hybridization images from the Allen Developing Mouse Brain Atlas showing expression of Dlx5, Isl1 and Sox6 in the mouse developing ventral forebrain (E13.5). (c) Human-chimpanzee ventral DE genes are largely shared along the dorsal forebrain developmental trajectories. (d) Human-chimpanzee DE directionalities and magnitudes, and DE genes detection rates on the two trajectories. DE directionalities and magnitudes are consistent on the dorsal and MGE trajectories, with most of the shared DE genes showing the highest human-chimpanzee expression divergence at NPC. DE genes specifically detected on one trajectory have tendency of higher detection rates on the trajectory with detected human-chimpanzee DE.

**Extended Data Figure 13:**
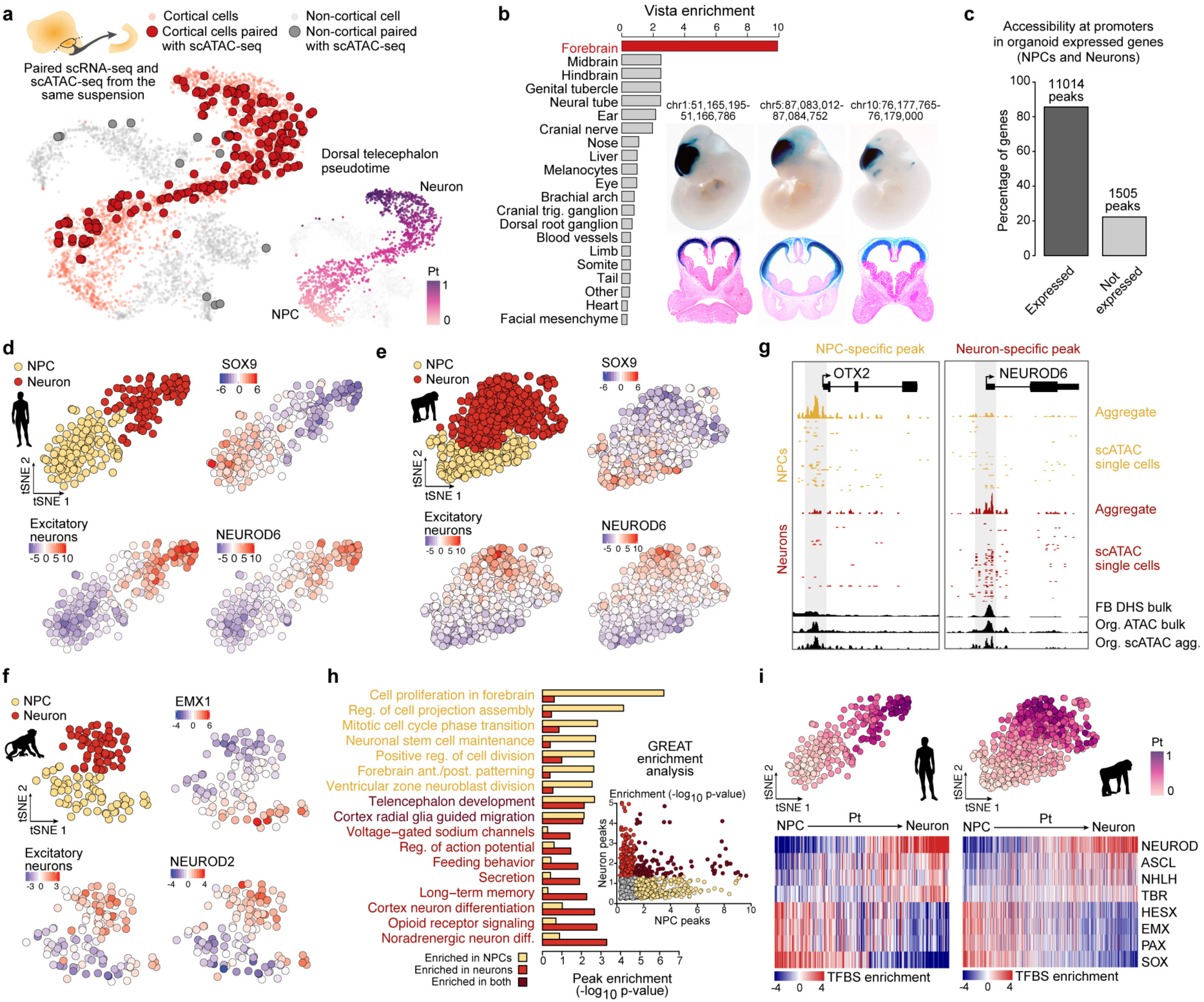
Chromatin accessibility in cerebral organoids. (a) t-SNE projection of highly variable gene expression in Fluidigm C1-based scRNA-seq data of cerebral organoids. Cortical cells are colored red, with larger points corresponding to cells with paired expression and chromatin accessibility data (data generated from the same cell suspension). 94.4% of cells with paired data are cortical, validating the cortical origins of the dissected cerebral organoid regions. (b) Cerebral organoid accessible peaks are significantly and highly enriched for overlapping human VISTA enhancers active in the forebrain relative to all other tissues (left). Three representative human VISTA enhancers with validated expression in E11.5 mouse forebrain that overlap cerebral organoid peaks are shown (out of 268 such enhancers). (c) Barplot showing the percentage of genes with accessible chromatin at the promoter of genes that are expressed or not expressed in human cerebral organoids. (d-f) t-SNE projection of bias-corrected deviations in accessibility for 7-mers within organoid scATAC-seq peaks per cell, with cells color coded by cell state (NPC, neuron) for human (d), chimpanzee (e), and macaque (f). Binding motif deviation Z-scores for representative transcription factors are shown, as well as deviation Z-scores for overlapping differentially accessible (DA) snATAC-seq peaks in mouse developing forebrain excitatory neurons^71^. (g) Signal intensity tracks of aggregated and individual single-cell chromatin accessibility data per cell state in human organoids at a NPC-specific promoter peak (left) and a neuron-specific promoter peak (right). For comparison, cerebral organoid bulk ATAC-seq chromatin accessibility data and human fetal brain bulk DNase-seq is shown. (h) Peaks identified as DA between NPC and neurons in human organoids were used as input for gene ontology enrichment analysis using the tool GREAT. Barplot showing the enrichment of representative enriched biological process gene ontology (GO) terms associated with human NPC DA peaks (gold) or human neuron DA peaks (light red) relative to all human organoid accessible peaks. Each point in the scatter plot represents a GO term and is colored by their enrichment in NPCs (yellow), neurons (red), both (dark red), or neither (grey). (i) tSNE plots colored by pseudotime, and heatmaps showing binding motif deviation Z-scores for selected transcription factors (rows) in all cells (columns) ordered in pseudotime for human (left) and chimpanzee (right).

**Extended Data Figure 14:**
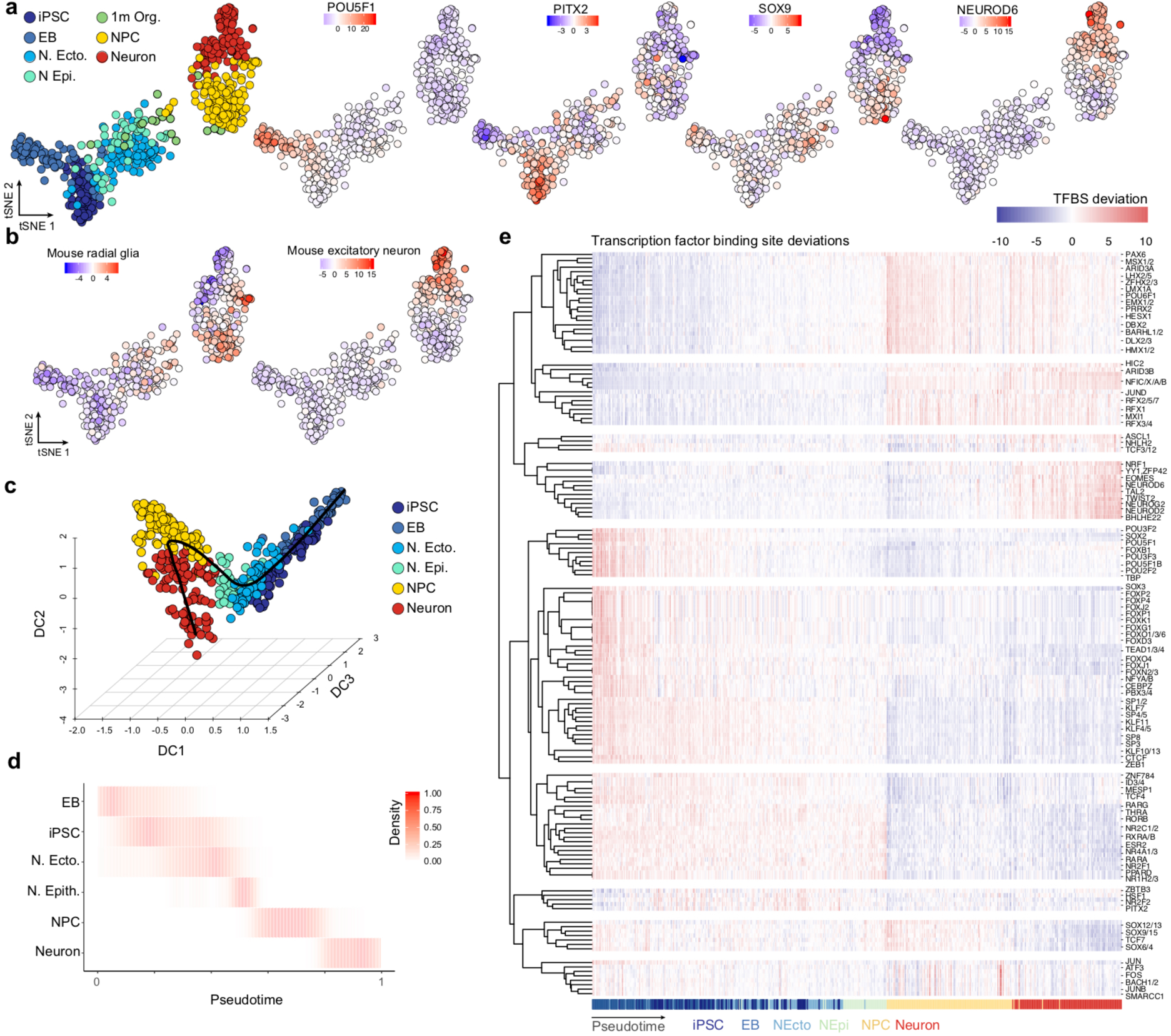
Chromatin accessibility during development from pluripotency to cortex formation in human cerebral organoids. (a) t-SNE projection of bias-corrected deviations in accessibility for 7-mers within scATAC-seq peaks per cell, with cells color coded by time point, and organoid data color coded by cell state (NPC, neuron). Binding motif deviation Z-scores for representative transcription factors are shown to the right. (b) t-SNE plot with cells colored by their deviation Z-score for overlapping differentially accessible snATAC-seq peaks from mouse developing forebrain^71^ radial glia cells (left) or excitatory neurons (right). (c) Diffusion map projection using the top 250 differentially accessible peaks per time point/cell state. The principle curve fit through the cells is shown as a black line. (d) Proportion of cells scaled by row is shown for each time point/cell state over pseudotime. (e) Heatmap representing the deviation Z-score of TF motifs that vary over the time course plotted for each cell across pseudotime.

**Extended Data Figure 15:**
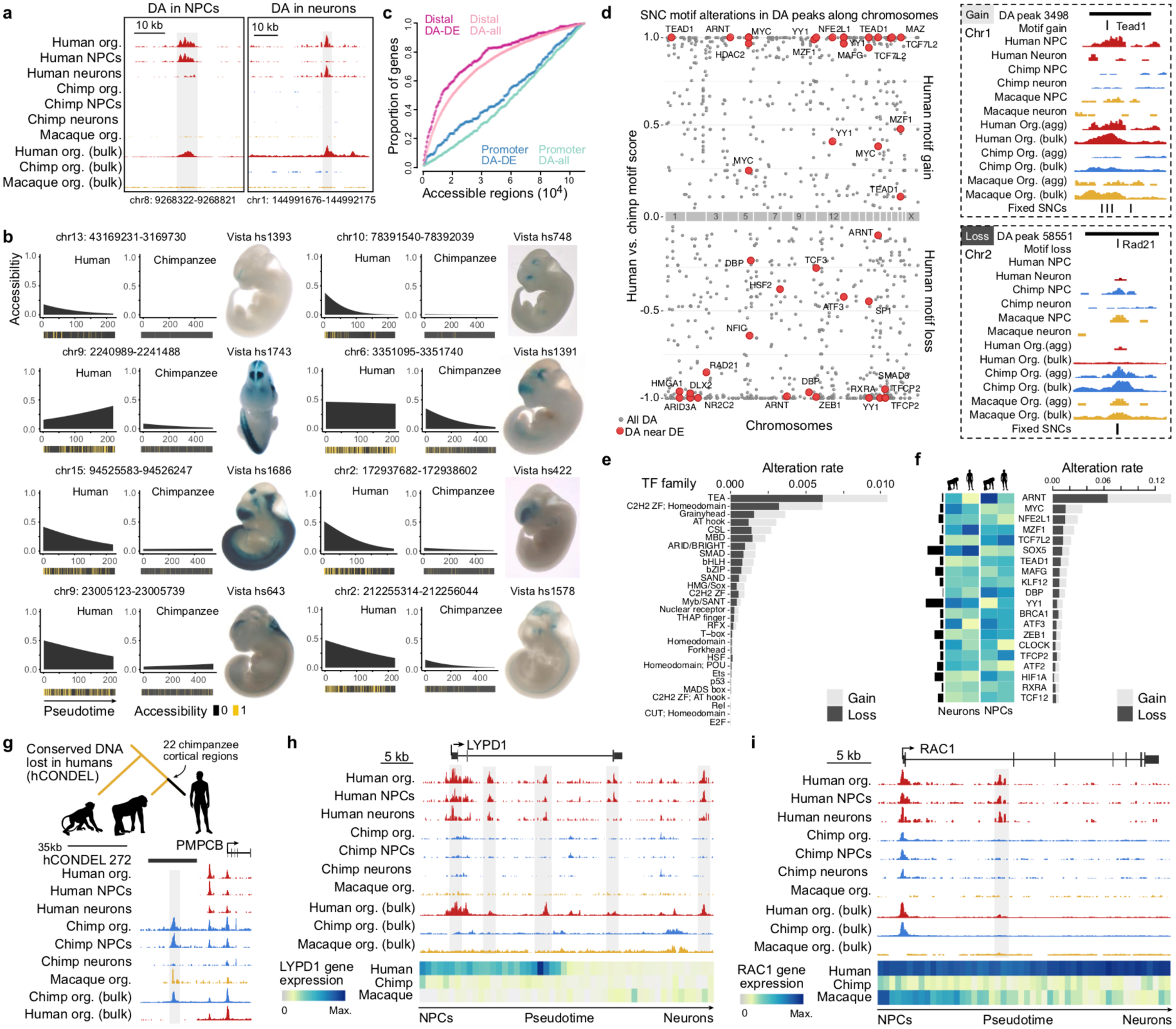
Chromatin accessibility differences in human and chimpanzee cerebral organoids. (a) Signal intensity tracks of aggregated single-cell and bulk chromatin accessibility data from human, chimpanzee, and macaque at a human-specific NPC-specific differentially accessible (DA) peak (left) and a human-specific neuron-specific DA peak (right). (b) The 8 most significant human-chimp organoid DA peaks containing a fixed SNC and accessible only in the cerebral organoid stage that overlap a VISTA human enhancer with validated activity in the developing mouse forebrain (out of 68 such cases). For each DA peak, the accessibility across pseudotime is shown for human and chimpanzee with heatmaps depicting cells where the peak is accessible (yellow) or inaccessible (black). The expression pattern of the overlapping VISTA enhancer in E11.5 mouse embryos is shown to the right. (c) Shown are the proportion of differentially expressed (DE) genes (dark color) or all expressed genes as background (light color) with a human-chimp organoid DA peak overlapping the promoter region (blue) or is distal to the promoter region (pink). The plot shows that DE genes between human and chimpanzee organoids are more likely to have a nearby DA peak than background. (d) Fixed SNCs predicted to significantly alter transcription factor binding within human-chimp organoid DA peaks, with the name of the altered motif shown for peaks linked to DE genes (red points). On the right, signal intensity tracks for a human motif gain (top) and human motif loss (bottom) within a human-chimp DA peak. (e) Altered transcription factor motifs grouped by family plotted for their alteration rate, which is the number of times a family member’s motif is altered in human-chimp organoid DA peaks divided by the number of times it’s detected in all accessible organoid peaks. (f) 20 transcription factors with the highest alteration rate, which is the number of times a motif is altered in human-chimp organoid DA peaks divided by the number of times it’s detected in all accessible organoids peaks. Heatmaps show their expression level in human and chimpanzee NPCs and neurons, with the bars to the left representing the average expression level across NPCs and neurons. (g) Example of an accessible peak in chimpanzee and macaque that overlaps a computationally-verified, non-polymorphic human conserved deletion (hCONDEL). (h-i) Signal intensity tracks of aggregated single-cell or bulk chromatin accessibility data from human, chimpanzee and macaque for two genes, (h) LYPD1 and (i) RAC1, that have higher expression specifically in humans. Gene expression is shown in heatmaps (below).

**Extended Data Figure 16:**
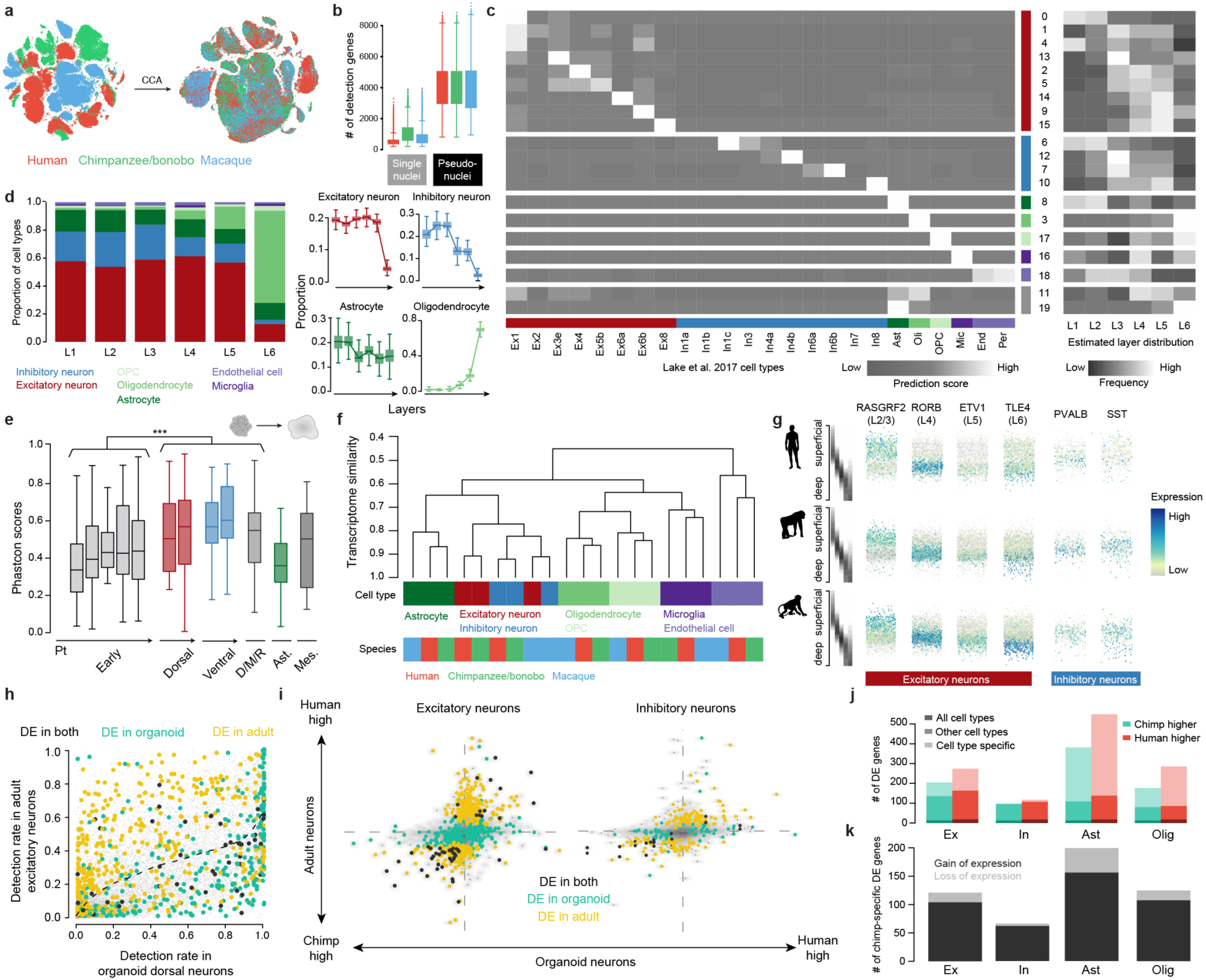
Supplementary analysis of human, chimpanzee and macaque adult brain single-nucleus RNA-seq. (a) The snRNA-seq data of adult brains in human, chimpanzee/bonobo and macaque were integrated using Seurat v3. (b) Boxplots (box, interquartile range (IQR); whisker, 1.5*IQR) showing the number of detected genes in single nuclei and pseudonuclei. (c) Heatmap showing the average prediction scores of each of the 20 identified clusters to each of the cell types reported by Lake et al. 2017, as well as their estimated distributions in different cortical layers in humans. Clusters are grouped in major cell classes. (d) Cell type composition of layers and layer distribution of cell types in human. (Left) Stacked bars showing the estimated cell type composition of different layers. (Right) Boxplots (box, interquartile range (IQR); whisker, 1.5*IQR) showing the estimated proportion per layer for four cell classes: excitatory neurons, inhibitory neurons, astrocytes and oligodendrocytes. (e) Genomic conservation based on average phastCon scores of developmental stage markers from iPSCs to neurons in human cerebral organoids (***: two-sided Wilcoxon’s rank sum test, P<0.0001). (f) Hierarchical clustering of the average transcriptome of seven cell classes in the three species. (g) Expression of layer markers (RASGRF2, RORB, ETV1, TLE4) in excitatory neurons and inhibitory neuron subtype markers (PVALB, SST) in inhibitory neurons, along the predicted laminar origin of the pseudonuclei in human, chimpanzee/bonobo and macaque. (h) Comparison of gene detection rates in organoid dorsal neurons and adult excitatory neurons, with human-chimpanzee DE genes in adult excitatory neurons colored in yellow, DE genes in organoid dorsal neurons colored in green, and shared DE genes colored in black. The dashed curve shows the fitted relationship between the two systems using all genes. Area below the curve represents higher detection rate in organoid neurons than adult neurons, and that above the curve represents higher detection rate in adult neurons. (i) Comparison of human-chimpanzee DE (left) between organoid dorsal neurons and adult excitatory neurons, as well as (right) between organoid ventral MGE neurons and adult inhibitory neurons. Densities are shown as grey scale shadows, with human-chimpanzee DE genes highlighted (yellow: DE only in adult; green: DE only in organoids; black: DE in both). (j) Number of human and chimp differentially expressed genes for cell classes based on all cell types, a subset of cell types and specific cell types. (k) Number of chimpanzee-specific DE genes across cell classes. The majority of the chimpanzee-specific DE genes have gain of expression (dark) rather than loss of expression (light).

## REFERENCES

1. Sousa, A. M. M., Meyer, K. A., Santpere, G., Gulden, F. O. & Sestan, N. Evolution of the Human Nervous System Function, Structure, and Development. Cell 170, 226–247, doi:10.1016/j.cell.2017.06.036 (2017).

2. Geschwind, D. H. & Rakic, P. Cortical evolution: judge the brain by its cover. Neuron 80, 633–647, doi:10.1016/j.neuron.2013.10.045 (2013).

3. Martin, R. D. Relative brain size and basal metabolic rate in terrestrial vertebrates. Nature 293, 57–60 (1981).

4. Paabo, S. The human condition-a molecular approach. Cell 157, 216–226, doi:10.1016/j.cell.2013.12.036 (2014).

5. Preuss, T. M. The human brain: rewired and running hot. Annals of the New York Academy of Sciences 1225 Suppl 1, E182–191, doi:10.1111/j.1749-6632.2011.06001.x (2011).

6. Somel, M., Liu, X. & Khaitovich, P. Human brain evolution: transcripts, metabolites and their regulators. Nat Rev Neurosci 14, 112–127, doi:10.1038/nrn3372 (2013).

7. Lui, J. H., Hansen, D. V. & Kriegstein, A. R. Development and evolution of the human neocortex. Cell 146, 18–36, doi:10.1016/j.cell.2011.06.030 (2011).

8. Florio, M. & Huttner, W. B. Neural progenitors, neurogenesis and the evolution of the neocortex. Development 141, 2182–2194, doi:10.1242/dev.090571 (2014).

9. He, Z. et al. Comprehensive transcriptome analysis of neocortical layers in humans, chimpanzees and macaques. Nature neuroscience 20, 886–895, doi:10.1038/nn.4548 (2017).

10. Sousa, A. M. M. et al. Molecular and cellular reorganization of neural circuits in the human lineage. Science 358, 1027–1032, doi:10.1126/science.aan3456 (2017).

11. Somel, M. et al. Transcriptional neoteny in the human brain. Proceedings of the National Academy of Sciences of the United States of America 106, 5743–5748, doi:10.1073/pnas.0900544106 (2009).

12. Brawand, D. et al. The evolution of gene expression levels in mammalian organs. Nature 478, 343–348, doi:10.1038/nature10532 (2011).

13. Konopka, G. et al. Human-specific transcriptional networks in the brain. Neuron 75, 601–617, doi:10.1016/j.neuron.2012.05.034 (2012).

14. Zhu, Y. et al. Spatiotemporal transcriptomic divergence across human and macaque brain development. Science 362, doi:10.1126/science.aat8077 (2018).

15. Lancaster, M. A. et al. Cerebral organoids model human brain development and microcephaly. Nature 501, 373–379, doi:10.1038/nature12517 (2013).

16. Marchetto, M. C. et al. Differential L1 regulation in pluripotent stem cells of humans and apes. Nature 503, 525–529, doi:10.1038/nature12686 (2013).

17. Camp, J. G. et al. Human cerebral organoids recapitulate gene expression programs of fetal neocortex development. Proceedings of the National Academy of Sciences of the United States of America 112, 15672–15677, doi:10.1073/pnas.1520760112 (2015).

18. Otani, T., Marchetto, M. C., Gage, F. H., Simons, B. D. & Livesey, F. J. 2D and 3D Stem Cell Models of Primate Cortical Development Identify Species-Specific Differences in Progenitor Behavior Contributing to Brain Size. Cell stem cell 18, 467–480, doi:10.1016/j.stem.2016.03.003 (2016).

19. Mora-Bermudez, F. et al. Differences and similarities between human and chimpanzee neural progenitors during cerebral cortex development. eLife 5, doi:10.7554/eLife.18683 (2016).

20. Kronenberg, Z. N. et al. High-resolution comparative analysis of great ape genomes. Science 360, doi:10.1126/science.aar6343 (2018).

21. Pollen, A. A. et al. Establishing Cerebral Organoids as Models of Human-Specific Brain Evolution. Cell 176, 743-756 e717, doi:10.1016/j.cell.2019.01.017 (2019).

22. Amiri, A. et al. Transcriptome and epigenome landscape of human cortical development modeled in organoids. Science 362, doi:10.1126/science.aat6720 (2018).

23. Quadrato, G. et al. Cell diversity and network dynamics in photosensitive human brain organoids. Nature 545, 48–53, doi:10.1038/nature22047 (2017).

24. Birey, F. et al. Assembly of functionally integrated human forebrain spheroids. Nature 545, 54–59, doi:10.1038/nature22330 (2017).

25. Weinreb, C., Wolock, S. & Klein, A. M. SPRING: a kinetic interface for visualizing high dimensional single-cell expression data. Bioinformatics 34, 1246–1248, doi:10.1093/bioinformatics/btx792 (2018).

26. Miller, J. A. et al. Transcriptional landscape of the prenatal human brain. Nature 508, 199–206, doi:10.1038/nature13185 (2014).

27. Nowakowski, T. J. et al. Spatiotemporal gene expression trajectories reveal developmental hierarchies of the human cortex. Science 358, 1318–1323, doi:10.1126/science.aap8809 (2017).

28. La Manno, G. et al. RNA velocity of single cells. Nature 560, 494–498, doi:10.1038/s41586-018-0414-6 (2018).

29. Bakken, T. E. et al. A comprehensive transcriptional map of primate brain development. Nature 535, 367–375, doi:10.1038/nature18637 (2016).

30. Renner, M. et al. Self-organized developmental patterning and differentiation in cerebral organoids. EMBO J, doi:10.15252/embj.201694700 (2017).

31. Marchetto, M. C. et al. Species-specific maturation profiles of human, chimpanzee and bonobo neural cells. eLife 8, doi:10.7554/eLife.37527 (2019).

32. Leigh, S. R. Brain growth, life history, and cognition in primate and human evolution. American journal of primatology 62, 139–164, doi:10.1002/ajp.20012 (2004).

33. Gould, S. J. Ontogeny and Phylogeny. (Harvard University Press, 1977).

34. Dougherty, M. L. et al. Transcriptional fates of human-specific segmental duplications in brain. Genome research 28, 1566–1576, doi:10.1101/gr.237610.118 (2018).

35. Florio, M. et al. Evolution and cell-type specificity of human-specific genes preferentially expressed in progenitors of fetal neocortex. eLife 7, doi:10.7554/eLife.32332 (2018).

36. Fiddes, I. T. et al. Human-Specific NOTCH2NL Genes Affect Notch Signaling and Cortical Neurogenesis. Cell 173, 1356–1369 e1322, doi:10.1016/j.cell.2018.03.051 (2018).

37. Dennis, M. Y. et al. Evolution of human-specific neural SRGAP2 genes by incomplete segmental duplication. Cell 149, 912–922, doi:10.1016/j.cell.2012.03.033 (2012).

38. Florio, M. et al. Human-specific gene ARHGAP11B promotes basal progenitor amplification and neocortex expansion. Science 347, 1465–1470, doi:10.1126/science.aaa1975 (2015).

39. Roadmap Epigenomics, C. et al. Integrative analysis of 111 reference human epigenomes. Nature 518, 317–330, doi:10.1038/nature14248 (2015).

40. Schep, A. N., Wu, B., Buenrostro, J. D. & Greenleaf, W. J. chromVAR: inferring transcription-factor-associated accessibility from single-cell epigenomic data. Nature methods 14, 975–978, doi:10.1038/nmeth.4401 (2017).

41. Visel, A., Minovitsky, S., Dubchak, I. & Pennacchio, L. A. VISTA Enhancer Browser--a database of tissue-specific human enhancers. Nucleic acids research 35, D88–92, doi:10.1093/nar/gkl822 (2007).

42. Villar, D. et al. Enhancer evolution across 20 mammalian species. Cell 160, 554–566, doi:10.1016/j.cell.2015.01.006 (2015).

43. Reilly, S. K., et al. Evolutionary genomics. Evolutionary changes in promoter and enhancer activity during human corticogenesis. Science 347, 1155–1159, doi:10.1126/science.1260943 (2015).

44. Prufer, K. et al. The complete genome sequence of a Neanderthal from the Altai Mountains. Nature 505, 43–49, doi:10.1038/nature12886 (2014).

45. Lindblad-Toh, K. et al. A high-resolution map of human evolutionary constraint using 29 mammals. Nature 478, 476–482, doi:10.1038/nature10530 (2011).

46. Prabhakar, S., Noonan, J. P., Paabo, S. & Rubin, E. M. Accelerated evolution of conserved noncoding sequences in humans. Science 314, 786, doi:10.1126/science.1130738 (2006).

47. Gittelman, R. M. et al. Comprehensive identification and analysis of human accelerated regulatory DNA. Genome research 25, 1245–1255, doi:10.1101/gr.192591.115 (2015).

48. McLean, C. Y. et al. Human-specific loss of regulatory DNA and the evolution of human-specific traits. Nature 471, 216–219, doi:10.1038/nature09774 (2011).

49. Tekinay, A. B. et al. A role for LYNX2 in anxiety-related behavior. Proceedings of the National Academy of Sciences of the United States of America 106, 4477–4482, doi:10.1073/pnas.0813109106 (2009).

50. Reijnders, M. R. F. et al. RAC1 Missense Mutations in Developmental Disorders with Diverse Phenotypes. American journal of human genetics 101, 466–477, doi:10.1016/j.ajhg.2017.08.007 (2017).

51. Stuart, T. et al. Comprehensive integration of single cell data. bioRxiv, 460147, doi:10.1101/460147 (2018).

52. di Gregorio, M. C. et al. pH sensitive tubules of a bile acid derivative: a tubule opening by release of wall leaves. Physical chemistry chemical physics : PCCP 15, 7560–7566, doi:10.1039/c3cp00121k (2013).

53. Halevi, S. et al. Conservation within the RIC-3 gene family. Effectors of mammalian nicotinic acetylcholine receptor expression. The Journal of biological chemistry 278, 34411–34417, doi:10.1074/jbc.M300170200 (2003).

54. Kilpinen, H. et al. Common genetic variation drives molecular heterogeneity in human iPSCs. Nature 546, 370–375, doi:10.1038/nature22403 (2017).

55. Thomson, J. A. et al. Embryonic stem cell lines derived from human blastocysts. Science 282, 1145–1147 (1998).

56. Okita, K. et al. An efficient nonviral method to generate integration-free human-induced pluripotent stem cells from cord blood and peripheral blood cells. Stem Cells 31, 458–466, doi:10.1002/stem.1293 (2013).

57. Picelli, S. et al. Smart-seq2 for sensitive full-length transcriptome profiling in single cells. Nature methods 10, 1096–1098, doi:10.1038/nmeth.2639 (2013).

58. Kang, H. M. et al. Multiplexed droplet single-cell RNA-sequencing using natural genetic variation. Nature biotechnology, doi:10.1038/nbt.4042 (2017).

59. Butler, A., Hoffman, P., Smibert, P., Papalexi, E. & Satija, R. Integrating single-cell transcriptomic data across different conditions, technologies, and species. Nature biotechnology 36, 411–420, doi:10.1038/nbt.4096 (2018).

60. Angerer, P. et al. destiny: diffusion maps for large-scale single-cell data in R. Bioinformatics 32, 1241–1243, doi:10.1093/bioinformatics/btv715 (2016).

61. He, Z., Bammann, H., Han, D., Xie, G. & Khaitovich, P. Conserved expression of lincRNA during human and macaque prefrontal cortex development and maturation. Rna 20, 1103–1111, doi:10.1261/rna.043075.113 (2014).

62. Buenrostro, J. D., Giresi, P. G., Zaba, L. C., Chang, H. Y. & Greenleaf, W. J. Transposition of native chromatin for fast and sensitive epigenomic profiling of open chromatin, DNA-binding proteins and nucleosome position. Nature methods 10, 1213–1218, doi:10.1038/nmeth.2688 (2013).

63. Buenrostro, J. D. et al. Single-cell chromatin accessibility reveals principles of regulatory variation. Nature 523, 486–490, doi:10.1038/nature14590 (2015).

64. Renaud, G., Stenzel, U. & Kelso, J. leeHom: adaptor trimming and merging for Illumina sequencing reads. Nucleic acids research 42, e141, doi:10.1093/nar/gku699 (2014).

65. Renaud, G., Stenzel, U., Maricic, T., Wiebe, V. & Kelso, J. deML: robust demultiplexing of Illumina sequences using a likelihood-based approach. Bioinformatics 31, 770–772, doi:10.1093/bioinformatics/btu719 (2015).

66. Li, H. et al. The Sequence Alignment/Map format and SAMtools. Bioinformatics 25, 2078–2079, doi:10.1093/bioinformatics/btp352 (2009).

67. Consortium, E. P. An integrated encyclopedia of DNA elements in the human genome. Nature 489, 57–74, doi:10.1038/nature11247 (2012).

68. Zhang, Y. et al. Model-based analysis of ChIP-Seq (MACS). Genome biology 9, R137, doi:10.1186/gb-2008-9-9-r137 (2008).

69. Robinson, J. T., et al. Integrative genomics viewer. Nature biotechnology 29, 24–26, doi:10.1038/nbt.1754 (2011).

70. Pliner, H. A. et al. Cicero Predicts cis-Regulatory DNA Interactions from Single-Cell Chromatin Accessibility Data. Molecular cell 71, 858–871 e858, doi:10.1016/j.molcel.2018.06.044 (2018).

71. Preissl, S. et al. Single-nucleus analysis of accessible chromatin in developing mouse forebrain reveals cell-type-specific transcriptional regulation. Nature neuroscience 21, 432–439, doi:10.1038/s41593-018-0079-3 (2018).

72. Suzuki, K. et al. Targeted gene correction minimally impacts whole-genome mutational load in human-disease-specific induced pluripotent stem cell clones. Cell stem cell 15, 31–36, doi:10.1016/j.stem.2014.06.016 (2014).

73. Mayer, C. et al. Developmental diversification of cortical inhibitory interneurons. Nature 555, 457–462, doi:10.1038/nature25999 (2018).

74. McLean, C. Y. et al. GREAT improves functional interpretation of cis-regulatory regions. Nature biotechnology 28, 495–501, doi:10.1038/nbt.1630 (2010).

75. Quinlan, A. R. & Hall, I. M. BEDTools: a flexible suite of utilities for comparing genomic features. Bioinformatics 26, 841–842, doi:10.1093/bioinformatics/btq033 (2010).

76. Cagan, A. et al. Natural Selection in the Great Apes. Molecular biology and evolution 33, 3268–3283, doi:10.1093/molbev/msw215 (2016).

77. Peyregne, S., Boyle, M. J., Dannemann, M. & Prufer, K. Detecting ancient positive selection in humans using extended lineage sorting. Genome research, doi:10.1101/gr.219493.116 (2017).

78. Chintalapati, M., Dannemann, M. & Prufer, K. Using the Neandertal genome to study the evolution of small insertions and deletions in modern humans. BMC evolutionary biology 17, 179, doi:10.1186/s12862-017-1018-8 (2017).

79. Fu, Y. et al. FunSeq2: a framework for prioritizing noncoding regulatory variants in cancer. Genome biology 15, 480, doi:10.1186/s13059-014-0480-5 (2014).

80. Lambert, S. A. et al. The Human Transcription Factors. Cell 172, 650–665, doi:10.1016/j.cell.2018.01.029 (2018).

81. Yu, Q. & He, Z. Comprehensive investigation of temporal and autism-associated cell type composition-dependent and independent gene expression changes in human brains. Scientific reports 7, 4121, doi:10.1038/s41598-017-04356-7 (2017).

82. Hafemeister, C. & Satija, R. Normalization and variance stabilization of single-cell RNA-seq data using regularized negative binomial regression. bioRxiv, 576827, doi:10.1101/576827 (2019).

