## Supplementary Materials for "Single-cell genomic atlas of great ape cerebral organoids uncovers human-specific features of brain development"

### SUPPLEMENTARY TABLES

Supplementary Table 1: Overview of single-cell genomic experiments

Supplementary Table 2: Highly variable genes used in SPRING and Reference Similarity Spectrum analyses

Supplementary Table 3: Metadata annotations of human cells from scRNA-seq

Supplementary Table 4: Cluster marker genes from heterogeneity analysis of human whole trajectory data

Supplementary Table 5: Metadata annotations of chimpanzee cells from scRNA-seq

Supplementary Table 6: Cluster marker genes from heterogeneity analysis of chimpanzee whole trajectory data

Supplementary Table 7: Metadata annotations of differentially expressed genes between human, chimpanzee, and macaque in cerebral organoids

Supplementary Table 8: Metadata annotations of cells from scATAC-seq

Supplementary Table 9: Metadata annotations of differentially accessible peaks between human and chimpanzee cerebral organoids

Supplementary Table 10: Metadata annotations of peaks overlapping human conserved deletions and single nucleotide changes in differentially accessible peaks

Supplementary Table 11: Highly variable genes used in adult brain single-nuclei RNA-seq data integration from human, chimpanzee and macaque

Supplementary Table 12: Metadata annotations of nuclei and pseudonuclei from snRNA-seq

Supplementary Table 13: Major cell class marker genes of human, chimpanzee and macaque adult brain

Supplementary Table 14: Metadata annotations of differentially expressed genes between human, chimpanzee and macaque in adult brains
